# Estimating the additive genetic variance for relative fitness from changes in allele frequency

**DOI:** 10.1101/2025.05.13.653886

**Authors:** Manas Geeta Arun, Aidan Angus-Henry, Darren J. Obbard, Jarrod D. Hadfield

## Abstract

The rate of adaptation is equal to the additive genetic variance for relative fitness (*V*_*A*_) in the population. Estimating *V*_*A*_ typically involves obtaining suitable measures of fitness on a large number of individuals with known pairwise relatedness. Such data are hard to collect and the results are often sensitive to the definition of fitness used. Here, we present a new method for estimating *V*_*A*_ that does not involve making measurements of fitness on individuals, but instead tracks changes in the genetic composition of the population. First, we show that *V*_*A*_ can readily be expressed as a function of the genome-wide diversity/linkage disequilibrium matrix and genome-wide expected change in allele frequency due to selection. We then show how independent experimental replicates can be used to infer the expected change in allele frequency due to selection and then estimate *V*_*A*_ via a linear mixed model. Finally, using individual-based simulations, we demonstrate that our approach yields precise and accurate estimates over a range of biologically plausible scenarios.

**Article summary:** Conventional approaches for estimating the heritable component of fitness variation (*V*_*A*_) have steep methodological, statistical, and even definitional challenges. Here, the authors present a new method that overcomes many of these issue by modelling *V*_*A*_ using selection-induced changes to a population’s genetic composition. The authors develop novel mathematical theory and an inference approach that uses independent experimental populations derived from the same ancestral population. Individual based simulations show that this method provides unbiased and precise estimates of *V*_*A*_. This opens the door for future studies investigating the genomic distribution of *V*_*A*_, a key factor driving Darwinian evolution.

## Introduction

Despite its simplicity, the fundamental theorem of natural selection (FTNS) (Fisher, 1930, 1958) is arguably one of the most central results in evolutionary biology, providing a concise mathematical statement of how quickly a population is expected to adapt. It describes the per generation gain in the mean fitness of a population as a result of natural selection, assuming the ‘environment’ (including variables intrinsic to the population, such as density or allele frequencies) are held constant (Ewens, 1989; Frank and Slatkin, 1992). The crucial insight the FTNS provides is that the rate of this increase in mean fitness is exactly equal to the additive genetic variance for relative fitness (*V*_*A*_) in the population (Burt, 1995; Grafen, 2015). Consequently, estimating *V*_*A*_ is one of the ‘*holy grails*’ of evolutionary genetics (Walsh, 2022), despite the FTNS being criticised for its inability to predict the actual gain in mean fitness (Price, 1972; Ewens, 1989, 2024) (also see Edwards (1994) for an historical review of the debate around FTNS).

A number of attempts have been made to measure *V*_*A*_ in wild populations. Typ-ically, these have involved long term studies on natural populations in which the lifetime reproductive success of a large number of individuals has been measured. Combining these fitness data with information on the relatedness among individuals in a (generalised) linear mixed model approach yields estimates of *V*_*A*_ (Kruuk, 2004). This is far from straightforward in natural populations, as it can be notoriously difficult to tease apart additive genetic effects from common environmental effects, such as parental effects (Kruuk and Hadfield, 2007; Shaw and Shaw, 2014). In addition, wild study systems are rarely closed, meaning emigration can be misinterpreted as mortality and offspring sired outside the study area can be overlooked. Furthermore, many studies on wild populations lack genetic pedigrees, and as a consequence may miss substantial fitness variation acquired through undetected polygamy (Vedder *et al*., 2011; Charmantier and Sheldon, 2006). In addition to these biases, estimates from wild populations also tend to come with considerable uncertainty. Burt (1995) reviewed studies estimating *V*_*A*_ in three species of plants and three species of animals, and found that most estimates of *V*_*A*_ were not significantly different from zero. They argued that the upper bound for estimates of *V*_*A*_ could be as high as 0.3, but most likely less than 0.1. Consistent with this, and using a larger data set, Hendry *et al*. (2018) reported that estimates of *V*_*A*_ varied between zero and 0.85, with the vast majority of estimates (73%) being less than 0.2. Overall, the mean 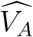 across studies was 0.08. In a recent meta-analysis, Bonnet *et al*. (2022) applied Bayesian quantitative genetic methods to data obtained from 19 long term studies on wild vertebrate populations. They reported a meta-analytic mean *V*_*A*_ of 0.185 across studies, considerably higher than those obtained by Burt (1995) and Hendry *et al*. (2018). Surprisingly, estimates of *V*_*A*_ in populations of Spotted Hyenas (*Crocuta crocuta*), as well as two of the three populations of Blue Tits (*Cyanistes caeruleus*) were higher than 0.4. This is a remarkable finding, since it suggests that growth rates in these populations should increase nearly 1.5 fold every generation due to selection, provided the environment remains constant. All three meta-analyses investigating *V*_*A*_ (Burt, 1995; Hendry *et al*., 2018; Bonnet *et al*., 2022) have detected substantial variability between study systems in their estimates of *V*_*A*_. Although most of this variability will be sampling error, the best estimate suggests the variability in actual *V*_*A*_ across populations is also large with a standard deviation of 0.11, although there is substantial uncertainty about its exact value (95% credible intervals: 0.01–0.26) (Bonnet *et al*., 2022).

Measuring *V*_*A*_ in the laboratory is considerably more straightforward, and involves either quantitative genetic breeding designs such as diallel crosses and full-sib half-sib experiments (Falconer and Mackay, 1996; Lynch and Walsh, 1998), or experimental techniques such as hemiclonal analysis (Abbott and Morrow, 2011). While the controlled environment of the laboratory can, to a large extent, help overcome some of the challenges faced by field studies, laboratory environments often lack important features that are likely to generate fitness variation, such as parasites, predators, and competitors. Therefore, it is not entirely clear if laboratory estimates of *V*_*A*_ are particularly relevant. Furthermore, many laboratory studies standardise their fitness measurements in a way that makes it hard to infer estimates of *V*_*A*_ (eg. Ruzicka *et al*. (2019)) or work with absolute fitness but then fail to report *V*_*A*_ (eg. Singh *et al*. (2023)). However, these studies do suggest that genetic variance for fitness is likely highly dependent on the specific environment in which fitness is assayed (eg. Punzalan *et al*. (2014)). In one of the few laboratory studies that estimated the genetic variance for *relative* fitness, Martinossi-Allibert *et al*. (2018) reported estimates from isofemale lines of the bean beetle, *Acanthoscelides obtectus*, that were highly sensitive to evolutionary history, assay environment, and sex, being higher in males (0.13-0.42) than in females (0.013-0.056). A compromise between the precision of the laboratory environment and the biotic and abiotic complexity encountered by wild populations can be achieved by working with experimental populations established in the field, an approach especially tractable in annual plants. Working with field populations of the annual legume *Chamaecrista fasciculata*, Kulbaba *et al*. (2019) found estimates of *V*_*A*_ that varied considerably among populations and years. Interestingly, many of their estimates were appreciably larger, with the largest estimate (calculated from Table 1 in the correction) being 3.05.

**Table 1:** The ranges for the rates for non-neutral mutations used in the history phase of the full simulations implemented in msprime (*µ*_*msp*_) (the coalescent simulation) and SLiM (*µ*_*SLiM*_) (the forward simulation of the history), along with the ranges for the resulting number of non-neutral sites segregating in the population at the end of the history phase 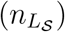 in simulations with different values of the absolute degree of dominance (|*κ*|), the mean of the gamma distribution from which effect sizes for log absolute fitness were sampled for non-neutral mutations (*E*[|*η*|]), and the ratio of the rate of beneficial mutations to the rate of deleterious mutations in the history phase (*µ*_*ben*_ : *µ*_*del*_). Note that the ranges for the number of selected loci, 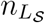, are only shown for the simulations where the map length in the history phase was 0.5 morgan.

| $\vert \kappa\vert $ | $E[\vert \eta\vert ]$ | $\mu_{ben} : \mu_{del}$ | $\mu_{msp}$ | $\mu_{SLiM}$ | $n_{L_S}$ |
| --- | --- | --- | --- | --- | --- |
| 0 | 0.02 | 0 | $5.56 \times 10^{-8} - 5.56 \times 10^{-7}$ | $5.56 \times 10^{-7} - 5.56 \times 10^{-6}$ | 10058 – 63755 |
| 0 | 0.02 | 0.0002 | $5.56 \times 10^{-8} - 5.56 \times 10^{-7}$ | $5.56 \times 10^{-7} - 5.56 \times 10^{-6}$ | 9742 – 63797 |
| 0 | 0.02 | 0.02 | $2.0 \times 10^{-8} - 2.0 \times 10^{-7}$ | $2.0 \times 10^{-7} - 2.0 \times 10^{-6}$ | 3127 – 29999 |
| 0 | 0.03 | 0 | $3.6 \times 10^{-8} - 3.6 \times 10^{-7}$ | $3.6 \times 10^{-7} - 3.6 \times 10^{-6}$ | 6565 – 42097 |
| 0 | 0.06 | 0 | $1.8 \times 10^{-8} - 1.6 \times 10^{-7}$ | $1.8 \times 10^{-7} - 1.6 \times 10^{-6}$ | 2998 – 18011 |
| 0.5 | 0.03 | 0 | $7.2 \times 10^{-8} - 3.24 \times 10^{-7}$ | $7.2 \times 10^{-7} - 3.24 \times 10^{-6}$ | 13738 – 49299 |
| 0.75 | 0.03 | 0 | $7.5 \times 10^{-8} - 2.07 \times 10^{-7}$ | $7.5 \times 10^{-7} - 2.07 \times 10^{-6}$ | 16870 – 38695 |

A common difficulty for current field and laboratory approaches is that, while Darwinian fitness is a deceptively intuitive concept, there is little consensus on its precise definition. In fact, it has been argued that the appropriate definition of fitness can vary depending on the context (Hendry *et al*., 2018). In the absence of a universal definition, empiricists can only use a measure of fitness deemed most appropriate for their study system, and it is reasonable to assume that estimates of *V*_*A*_ are likely to be sensitive to the definition of fitness used. A useful illustration of this point is provided by two studies that estimated *V*_*A*_ in a wild population of Red Deer (*Cervus elaphus*), using largely overlapping datasets, but markedly different definitions of fitness (Kruuk *et al*., 2000; Foerster *et al*., 2007). Kruuk *et al*. (2000) defined fitness as the total number of progeny produced by an individual in its lifetime and estimated *V*_*A*_ to be 0.1, whereas Foerster *et al*. (2007) employed an alternative definition of fitness that measured an individual’s contribution to population growth (Coulson *et al*., 2006) and obtained the appreciably higher estimate of 0.64.

Some of the definitional difficulties of measuring *V*_*A*_ can be overcome by measuring *V*_*A*_ as the rate of adaptation, rather than comparing measures of fitness among relatives. One of the earliest attempts to explore this idea was made by Fowler *et al*. (1997) (also see Gardner *et al*. (2005)) in laboratory populations of *Drosophila melanogaster*. Using balancer chromosomes that allow recombination to be suppressed on the third chromosome, Fowler *et al*. (1997) could track the frequency trajectories of newly introduced wild-type third chromosomes over the course of 43 weeks. By modelling fitness using genotype frequency data for a number of different wild-type third chromosomes (Barton and Partridge, 2000), they demonstrated the presence of substantial *V*_*A*_ on the third chromosome. More recently, in a landmark study, Buffalo and Coop (2019) developed a method to estimate the amount of genome-wide allele frequency change that can be attributed to selection. The linchpin of their theory is the idea that linked selection induces across-generation covariances in allele frequency change at neutral loci, since associations between these neutral loci and their respective non-neutral backgrounds persist across generations. This new theoretical framework has the potential to pave the way for a powerful empirical tool to detect genomic signatures of linked selection (Buffalo and Coop, 2020; Simon and Coop, 2024). Of particular relevance here, Buffalo and Coop (2019) also show that their method can be used to obtain estimates of *V*_*A*_, albeit under some potentially restrictive assumptions.

In this study, we present an alternative theoretical framework that relates *V*_*A*_ to genome-wide changes in allele frequency. Using mathematical identities only, we show how *V*_*A*_ can be obtained from an initial linkage disequilibrium (LD) matrix and expected changes in allele frequency due to selection, without making any assumptions about patterns of gene action or the relationships between genotype fitnesses and genotype frequencies. Our approach, like that of Buffalo and Coop (2019), relies on temporal genomic data and does not necessitate measuring fitness in individuals. However, in contrast to the ‘bottom-up’ population genetic approach of Buffalo and Coop (2019), we use a ‘top-down’ quantitative genetic approach which simplifies and generalises some aspects of the problem.

Our aim here is three-fold. First, we derive our central theoretical result from first principles. Second, we develop the statistical machinery required to apply our result to real biological data, and validate it with individual based simulations that permit dominance effects, but assume random mating and an absence of epistasis. Third, we make a detailed comparison of our method with that of Buffalo and Coop (2019).

## Materials and methods

Table 3 summarises the notation used in the following text and the subsection ‘Assumptions retained in our approach’ (see below) summarises the assumptions involved in our approach.

### Outline of the theory

We consider a population consisting of *N* diploid individuals. We assume that there are *n*_*L*_ segregating biallelic loci in the population. Let *c*_*k,i*_ and *α*_*i*_ represent the proportion of copies of an arbitrarily chosen reference allele at locus *i* in individual *k* and Fisher’s average effect (Fisher, 1941) for relative fitness at locus *i*, respectively. The widely accepted mathematical definition of the *α*’s are the regression coefficients obtained from a multiple regression of the *c*’s on relative fitness, *w* (Fisher, 1941; Lee and Chow, 2013). The vector ***α*** can be expressed as

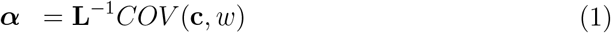

where **c** is the vector of predictors representing the *c*’s at all loci for an individual and **L** is a symmetric *n*_*L*_ *× n*_*L*_ matrix whose *i*^*th*^ (diagonal) element is the variance in the *c*’s at locus *i* computed over individuals, while the *ij*^*th*^ (off-diagonal) element describes the covariance between the *c*’s at locus *i* and locus *j* computed over individuals.

The breeding value for the relative fitness of individual *k* is then

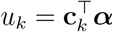

and the additive genetic variance for relative fitness is the variance of this quantity across individuals:

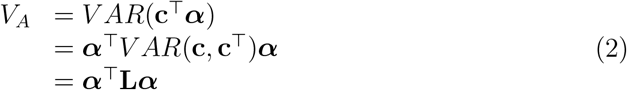

This follows from the fact that at any given point in time, *α*’s are constant across individuals. In the absence of mutation and meiotic drive, the allele frequency in parents is transmitted to offspring without bias, such that the vector of expected change in allele frequencies due to selection can be expressed as Robertson’s covariance (Robertson, 1966; Price, 1970; Queller, 2017):

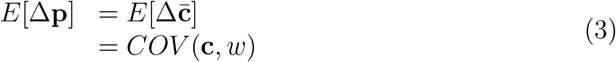

where the expectation is taken over the evolutionary process.

Substituting Equation 1 into Equation 3 gives *E*[Δ**p**] = **L*α***, which is the mul-tivariate analogue of Equation 10 in Kirkpatrick *et al*. (2002). Combining this with Equation 2 yields,

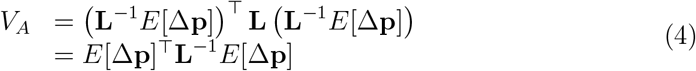

Equation 4 is a general result and involves no assumptions about the patterns of dominance or epistasis for fitness, or about patterns of mating. An intuitive explanation of why *V*_*A*_ can be calculated this way is to note Fisher’s Fundamental Theorem states that *V*_*A*_ is equal to the (partial) increase in mean fitness caused by evolutionary change through natural selection. Equation 2 can be expressed as a sum of *α*_*i*_*E*[Δ*p*_*i*_] over all loci, *i*:

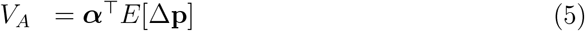

In this, *α* represents the proportional change in the population mean fitness that is caused by a unit change in allele frequency at a locus (Fisher, 1941; Kojima, 1959; Lee and Chow, 2013). Therefore, multiplying *α* by the actual change caused by natural selection, *E*[Δ*p*], we get the proportional change in mean fitness caused by evolutionary change by natural selection at that locus (see Eq 51.4 Price, 1972, also). If we then add these changes at every locus in the genome, we obtain the total proportional change in mean fitness due to evolutionary change by natural selection, and hence *V*_*A*_.

### Extending our approach to practical situations

Our theoretical approach assumes ***α*** or *E*[Δ**p**] are known. In reality, neither can be directly observed and must be inferred from data on observed allele frequency change, Δ**p**. Since Δ**p** will vary around *E*[Δ**p**] due to genetic drift, *E*[Δ**p**] must be inferred using replicate observations of Δ**p**. Since time cannot be replayed, we infer *E*[Δ**p**] through experimental replicates (see Buffalo and Coop (2020) also). Our theoretical model also assumes **L** is known, but since it is hard to generate experimental replicates without at least one round of reproduction, we condition on **L**_0_ (**L** in a generation prior to the first generation over which allele frequency change is measured). In what follows we will refer to the population at time zero (generation 0) as the ‘base population’. Our aim is then to approximate the additive genetic variance for fitness in the base population *V*_*A*_(0) as 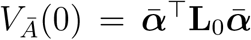, where 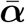 is the mean vector of average effects averaged over time and replicates. Note that if average effects are constant then *V*_*Ā*_(0) = *V*_*A*_(0), but if the average effects vary then *V*_*A*_(0) will in general exceed *V*_*Ā*_(0) which can be interpreted as the additive genetic *covariance* in fitness between replicates/time points for a population with genotypic structure equal to that in the base population (see Supplementary information S2). It is important to note that conventional methods for estimating *V*_*A*_ – such as those relying on pedigreed individuals – also estimate additive genetic *covariances* among environments in which relatives are assayed (Vehviläinen *et al*., 2008) and these covariances equal *V*_*A*_ only when the average effects remain constant.

We also allow allele frequency changes to be measured over multiple generations, rather than a single generation. Thus, 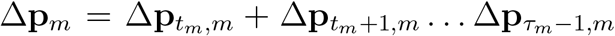 is the observed change in allele frequency from generation *t*_*m*_ to *τ*_*m*_ in replicate *m*, with Δ**p**_*t,m*_ being the change from time *t* to *t* + 1. Note that *t*_*m*_ (i.e. the generation in which we start recording allele frequency changes in replicate *m*) can be any positive integer. If replicate populations are derived from the base population with exactly one round of reproduction, *t*_*m*_ would be 1 for all *m. τ*_*m*_ can be any integer greater than *t*_*m*_. We first note that the total change in allele frequency in replicate *m* between times *t*_*m*_ and *τ*_*m*_ is

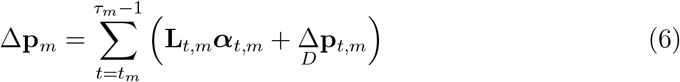

where **L**_*t,m*_***α***_*t,m*_ = *E*[Δ**p**_*t,m*_] is the expected change in allele frequency due to selection in replicate *m* between generation *t* and *t* + 1 and 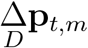 is the change due to drift. It is important to note that **L**_*t,m*_***α***_*t,m*_ captures responses to both direct and indirect selection. **L** will vary over time and we decompose **L** at a particular time/replicate into a part that can be predicted by **L**_0_ and the action of drift and recombination, and a part that cannot be predicted. To do this we decompose **L** into **L**^*′*^ and **L**^*′′*^, following the notation of Buffalo and Coop (2019). **L**^*′*^ represents the (co)variances in the *c*’s computed over the haploid genomes of all 2*N* gametic contributions that constitute the population. Thus, the diagonal elements of **L**^*′*^ are proportional to gametic (gene) diversity and the off-diagonals are proportional to gametic-phase disequilibrium. On the other hand, **L**^*′′*^ represents the (co)variances that arise due to alleles within the different gametic contributions of a genotype, and thus the diagonal elements are proportional to the additive coefficients of HardyWeinberg disequilibrium (Bulmer (1980) and Chapter 12 in Weir and Cockerham (1989)), and the off-diagonal elements are proportional to the nongametic-phase disequilibrium. Given this decomposition, the dynamics of **L** under drift and recombination are (See Supplementary information S1 for details and Hill and Robertson (1968) and Santiago and Caballero (1998)):

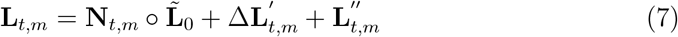

where *∘* is the Hadamard (i.e. element-wise) product, 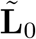 is the weighted sum of 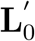 and 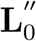, with the *ij*^*th*^ element of 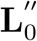 weighted by *r*_*ij*_*/*(1−*r*_*ij*_) and **N**_*t,m*_ is a matrix with the *ij*^*th*^ element being 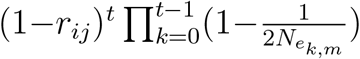 is the recombination rate between the two loci and 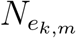 is the effective population size in generation *k* in replicate *m*. 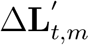 is a stochastic term that represents the accumulated change in **L**^*′*^ between generations 0 and *t* in replicate *m* that cannot be predicted. 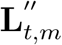 is the matrix of nongametic-phase disequilbria that arises in generation *t* in replicate *m*. In the absence of selection, 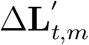 and 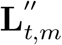 have zero expectation. In the following, it will be useful to designate 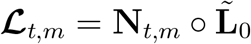 as the predicted **L**_*t,m*_, for *t >* 0, conditional on 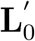 and 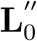. Also, we represent the deviation of **L**_*t,m*_ from this prediction as 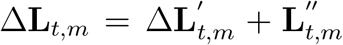. It will also be useful to denote 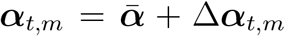 where Δ***α***_*t,m*_ is the deviation of the average effects (in generation *t* in replicate *m*) from the mean of the average effects across time and replicates. We can then decompose the change due to selection into two terms:

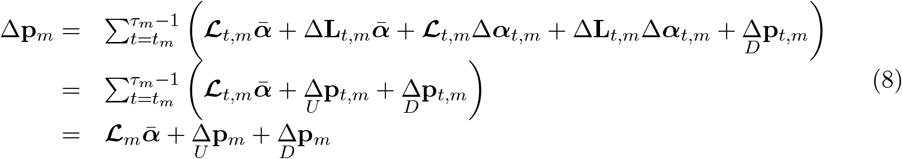

Here quantities subscripted with just *m* are the sums of the relevant quantity over *t* (for example, 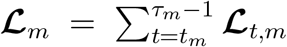). The predictable change due to selection is 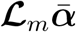 and is similar to that derived for a single locus in Santiago and Caballero (1998). The unpredictable change due to selection caused by stochastic changes in **L** and/or ***α*** is 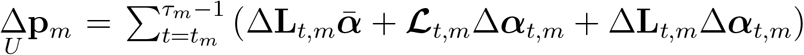. The total change due to drift is 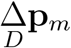. The 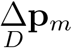 have zero expectation and are independent across replicates (Buffalo and Coop, 2019). In what follows we also assume the 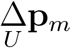 have zero expectation, although we believe there are two primary mechanisms by

which this assumption can fail. First, the model for the predicted changes in **L** (i.e. **ℒ**_*m*_) may be inaccurate such that the Δ**L**_*t,m*_ have non-zero expectation. Since **ℒ**_*m*_ is derived under the assumptions of drift and recombination only, directional changes in **L** induced by selection is an obvious mechanism. However, if the genetic architecture of fitness is sufficiently polygenic, such that selection coefficients associated with individual loci are small, selection-induced changes in **L** may be safely ignored. In other words, selection-induced changes in **L** may cause the total allele frequency change due to selection to be negligibly different from 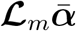, the expected allele frequency change due to selection. Second, the product Δ**L**_*t,m*_Δ***α***_*t,m*_ may have nonzero expectation if changes in the average effects are correlated with changes in patterns of diversity/LD. Again, selection-induced changes in Δ**L**_*t,m*_ is a possible mechanism. Additionally, unless all gene action is additive, changes in **L** also induce changes in ***α*** since then average effects depend on allele frequencies (Supplementary information S8).

However, calculating change in allele frequency over a single generation (i.e. *τ*_*m*_ − *t*_*m*_ = 1) would minimise any bias since any change in **L** due to selection should be minimised and the approximations that follow will be more accurate. Although the approximations will hold better when *τ*_*m*_ − *t*_*m*_ = 1, increasing *τ*_*m*_ − *t*_*m*_ (i.e. calculating change in allele frequency over multiple generations) will increase power since the changes in allele frequency due to selection will be larger.

Assuming both 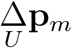 and 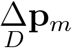 have zero expectation we can derive the mean and covariance structure (computed over the evolutionary process) of allele frequency change in a replicate as:

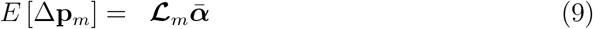

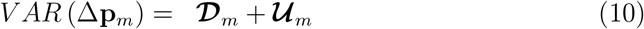

where **𝒟**_*m*_ and **𝒰**_*m*_ are the covariances due to the cumulative action of drift and the unpredictable response to selection, respectively. The drift (co)variances have expectation equal to 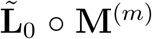 where 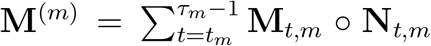 and the *ij*^*th*^ element of **M**_*t,m*_ is 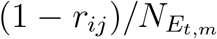 (Supplementary information S2). Note that since the predictable response to selection includes both direct and indirect selection, the relevant effective population size for the drift covariance in allele frequency does not include the effects of linked selection and for this reason we denote it as *N*_*E*_ rather than *N*_*e*_. Using this notation, *N*_*E*_ = 4*N/*(2 + *V*_*o*_) where *N* is the census population size and *V*_*o*_ is the variance in offspring number in the absence of additive genetic fitness variation (Wright, 1938). Although the dynamics of **L** (Equation 7, also see Supplementary information S1) were derived under drift and recombination in the absence of selection, and hence *N*_*e*_ = *N*_*E*_, the dynamical equations for **L** will better approximate reality when an effective population size that incorporates all excess variation in fitness is used. Consequently, we use *N*_*e*_ and *N*_*E*_ to distinguish the effective population sizes that are relevant for stochastic changes in **L** and allele frequency, respectively.

There is no easy form for the covariance due to the cumulative unpredictable response to selection and we simply denote it as **𝒰**_*m*_ (Supplementary information S2). While the drift terms will be independent across replicates, we also need to assume that 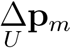 are independent across replicates for the covariance between replicates to be zero. An obvious source of non-independence would be if replicates are not initiated from individuals independently generated from Generation 0 individuals (i.e. *t*_*m*_ ≠ 1 for at least one *m*). If this were the case, the Δ**L** will be dependent across replicates due to the shared changes in **L** from generation 0 to the generation from which the replicates are derived. This could be minimised by initialising replicates as early as possible (ideally at *t*_*m*_ = 1).

### Inference outline

In the previous section, the mean average effects, 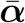, are treated as fixed and expectations and variances are taken over the evolutionary process, including variation in the average effects over time and replicates. When making inferences, we treat the mean average effects as random variables rather than fixed quantities (See Supplementary information S3 and Gianola *et al*. (2009) for a discussion on what this implies) and derive expectations and (co)variances taken over both the evolutionary process and the distribution of 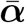. Using the laws of total expectation and (co)variance we obtain:

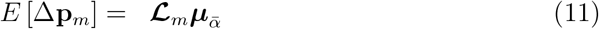

and

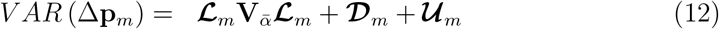

where 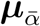 and 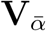 and are the mean and covariance structure of the mean average effects respectively. Critically, when deconditioning on 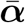 the cross-replicate covariances become non-zero:

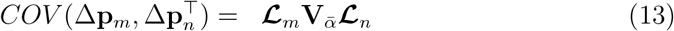

where *m* and *n* are a pair of replicates.

In Supplementary information S3 we determine permissible models for 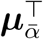 and 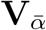. Of those, we identify the sensible biological model for the mean average effect:

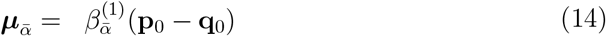

where **p**_0_ is the frequency of the reference alleles at each locus in the base population and **q**_0_ = **1** − **p**_**0**_. For the variance of the average effects, we identify the model

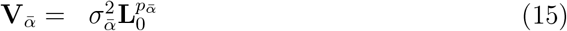

although in our analysis of simulated data we set the off-diagonal elements of **L**_0_ to zero in the above equation, such that the variance in average effects is simply a power function of the genetic diversities (Zeng *et al*., 2018). When 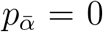 average effects are independent of genetic diversity, and when 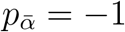 average effects scale inversely with genetic diversity. 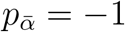 is a common assumption in many related approaches (e.g. Yang *et al*., 2011) and under this assumption 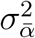 is the average contribution of a locus to the additive genic variance.

By applying sum of squares theory (Searle, 2006, page 355) to Equation 2 we can obtain the (posterior) expectation of *V*_*Ā*_(0) after averaging over the distribution of average effects (Supplementary information S3 and Gianola *et al*. (2009)):

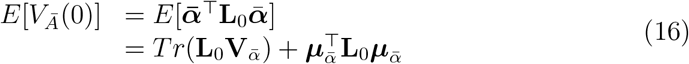

where we aim to estimate 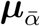 and 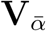 through Equations 11–13 using multiple evolutionary replicates starting from a common base population.

Rather than working with the allele frequency changes directly, we project them on to a new (reduced) basis and denote this new vector of changes as 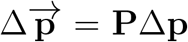 where **P** is some projection matrix. We chose a projection that collapses allele frequency changes into the non-null subspace of **L**_0_, since *V*_*Ā*_(0) only depends on this subspace (Supplementary information S4 and Supporting Information in de Los Campos *et al*. (2015)). To do this, let **U**_**L**_ be the eigenvectors of **L**_0_ with non-zero eigenvalues and then the drift covariance in the reduced subspace is 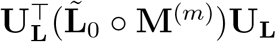. If we have **U**_2_ and **D**_2_ as the eigenvectors and a diagonal matrix of square-rooted eigenvalues of this matrix, then the projection matrix 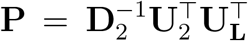 results in projected allele frequency changes that are identically and independently distributed under drift in the reduced subspace only (see Supplementary information S4).

The mean and covariance of the projected allele frequency changes due to predictable selection are:

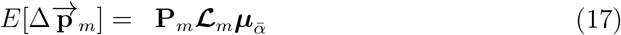

and

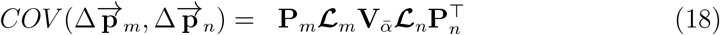

With 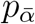 known, the model is a linear mixed model with covariance structure due to the predictable response to selection being proportional (by a factor 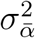 that is to be estimated) to

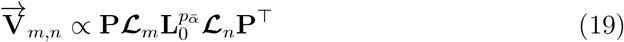

between replicates *m* and *n*. When **ℒ** ^(*m*)^ = **ℒ** ^(*n*)^ for all *n* and *m*, then this can more easily be fitted by incorporating locus as a random effect with the above covariance structure. The vector of expected values (shown for replicate *m*) is also proportional to

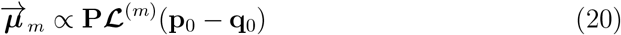

with the associated coefficient 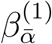 to be estimated.

We estimate the parameters of the model by treating it as a separable linear mixed model problem (Richards, 1961). Conditional on a value of 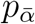 the model is linear mixed model and the conditional (restricted) maximum likelihood can be obtained from asreml (Butler *et al*., 2023). In order to maximise the (unconditional) likelihood we use the R function optim to find the value of 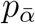 that results in the highest conditional likelihood. Note that the residual variance should equal one if there is no unpredictable response to selection.

Although linear (mixed) models generate unbiased estimates of 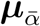, the quadratic form 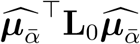 will be upwardly biased by sampling error in 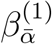. To correct for this bias, we use the inverse Hessian (conditional on the best estimate of 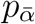) to get an approximate expression for the sampling variance 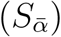 of 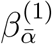, and use this in order to get an improved estimate of the quadratic form as follows (see Supplementary information S5 for a general derivation):

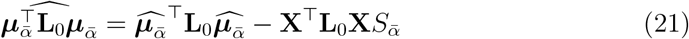

where **X** is the fixed effect design matrix.

In addition, allele frequency changes will rarely be known without error and will most likely be estimated using a pool-seq approach. In such cases, it is necessary to add an additional covariance structure into the model that accommodates the estimation error. In Supplementary information S6 we derive the covariance structure for projected allele-frequency change (in replicate *m*) as:

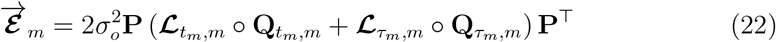

where element *ij* of a **Q** matrix is the number of reads that span site *i* and *j* divided by the product of the coverages at the two sites. When *i* = *j, Q*_*i,j*_ is simply the inverse of the coverage. When individuals have a constant probability of being sampled, 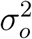 is expected to be one, but we treat it is a free parameter in order to capture any overdispersion (McCullagh and Nelder, 1989).

### Comparison with Buffalo and Coop (2019)

To connect our work with Buffalo and Coop (2019) (henceforth ‘B&C’) it will be useful to express the covariance matrix **L** in terms of a diagonal matrix of standard deviations, **B**, (half the square-root of the genetic diversities under random mating) and the correlation matrix, **R**, such that **L** = **BRB**. We can then split the response to selection in generation *t* into two parts:

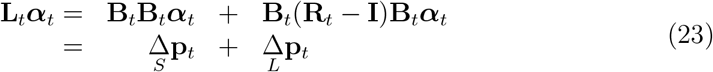

where the first term is due to direct selection at the loci, and the second term is due to linkage-disequilibria with other selected loci. It will also be useful to distinguish the additive *genetic* variance (*V*_*A*_(*t*)) from the additive *genic* variance (*V*_*a*_(*t*)) in generation *t*:

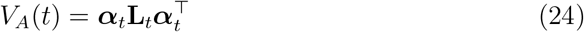

versus

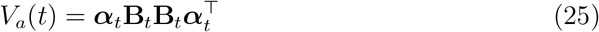

with the distinction being that the additive *genetic* variance (*V*_*A*_(*t*)) captures the contributions of both genetic diversities at individual loci and the linkage disequilibria between pairs of loci, while the additive *genic* variance (*V*_*a*_(*t*)) captures the contributions of genetic diversities only.

We can also think about the additive genetic/genic covariance in fitness between time points *t* and *τ* for a population with genetic structure equal to that in generation *t*:

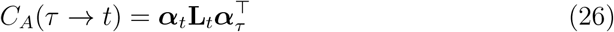

and

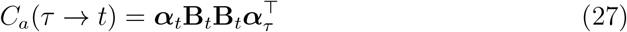

These will differ from *V*_*A*_(*t*) or *V*_*a*_(*t*) when the average effects at generation *τ* are different from those at generation *t*.

In the theory section above, we assumed **L**_*t*_ and ***α***_*t*_ were known and so the change due to both direct and linked selection are fixed quantities, as is *V*_*A*_(*t*). In contrast, B&C condition on **B**_*t*_ and ***α***_*t*_, and treat **R**_*t*_ as a random variable. Consequently, in B&C, the change due to direct selection and *V*_*a*_(*t*) are fixed, as in our approach, since genetic diversities and average effects are known. However, the change due to indirect selection and *V*_*A*_(*t*) is random since the linkage-disequilibrium is unknown. In our inference section we acknowledge that ***α*** cannot be directly observed, but develop a model for the distribution of *α*’s, where estimates of *V*_*A*_(*t*) can be made after marginalising ***α***. Similarly, in the approach of B&C the ***α***’s and the elements of **B** for selected sites are also not directly observed, but they make a number of strong assumptions that effectively allow the joint distribution of the average effects and selected site diversities to be marginalised.

In Supplementary information S7 we work through the derivation of B&C using our own notation and retaining a full multi-locus treatment. For easy comparison with the present work, here we summarise both the explicit and implicit assumptions underlying the general approach of B&C:

- A) Allele frequency changes are only measured over a single generation.
- B) There is no direct selection on the loci for which allele frequency change is measured.
- C) The reference allele at a neutral locus is chosen arbitrarily.
- D) The signed linkage-disequilibrium between selected sites is zero, which precludes processes such as Hill-Robertson interference. Under this assumption *V*_*A*_ = *V*_*a*_.
- E) Changes in **R** are due to recombination alone selection and drift are absent.
- F) Nongametic-phase linkage disequilibrium is absent.
- G) There is no relationship between the additive genic variation at a selected site and its LD with neutral sites, as measured through *R*_*t,ij*_*R*_*τ,ij*_ where *i* is a neutral locus and *j* a selected locus. Note that since genetic diversity determines the additive genic variation and LD (even measured as a correlation) is constrained by the genetic diversities at the two loci, this is unlikely to be met (Sved and Hill, 2018).
- H) The average effects are constant, i.e. ***α***_*t*_ = ***α***_*τ*_ if estimates are to be interpreted as the additive genic variance (or additive genetic variance, if Assumption D is met) in generation *τ* . If they are not constant, it is the additive genic/genetic covariance between generations *t* and *τ* that is measured.
- I) The initial expected LD-structure between selected and neutral loci, 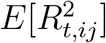, is approximated from an expression for 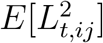, which is itself derived under mutation-recombination-drift balance (i.e. no selection). This assumption is partly relaxed in the Appendix by assuming the expected LD-structure between selected and neutral loci is equal to the observed LD-structure between all loci, not distinguishing between selected and neutral loci (Assumption I-b). However, in many cases the genetic diversity at selected loci will be less than that found at (or assumed for) neutral loci and so 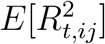 will likely be smaller than assumed (Sved and Hill, 2018).
- J) The ratio of genetic diversity in generation *τ* to genetic diversity in generation *t* (*ϕ*_*t,τ*_), is constant across all selected loci. Under this assumption, dividing the estimate of *V*_*a*_(*τ*) by *ϕ*_*t,τ*_ gives an estimate of *V*_*a*_(*t*).
- K) The recombination rate between two sites *g* base pairs apart is given by 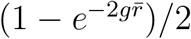 (i.e. Haldane’s (1919) mapping function) where 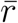 is the average crossover rate per site per generation.
- L) Neutral and selected loci are distributed uniformly and independently across the genome such that the distance between them has a triangular distribution.
- M) Since the genetic diversity at selected sites is not measured, it is assumed that *ϕ*_*t,τ*_ is equal to the average ratio of genetic diversity in generation *τ* to genetic diversity in generation *t* across all neutral loci.

When applying the theory to data, the method of B&C requires two additional assumptions:

- Ib) The average LD between selected and neutral sites is equal to the average LD observed between all sites: see Assumption I) above.
- N) The (co)variance in allele frequency changes divided through by the average genetic diversity is a good approximation for the (co)variance of weighted allele frequency changes where the weights are the inverse of the square root of the genetic diversities.
- O) If allele frequencies are estimated from a sample, then the estimation error is binomially distributed.

### Assumptions retained in our approach

Our approach works with diploid populations throughout, and assumes non-overlapping generations. While deriving our main theoretical result (Equation 4), we further assume an absence of meiotic drive and mutation. When extending our approach to practical situations and in our inference approach, we relax most of the assumptions made by B&C, but not all.

- Assumption E is only partly relaxed - the change in **R** does not assume a lack of drift but it does assume a lack of selection.
- We also make Assumption H if we choose to interpret *V*_*Ā*_(0) as as an additive genetic variance (i.e. *V*_*A*_(0)) rather than an additive genetic covariance over replicate/time-points.
- Assumptions E & H result in the unpredictable response to selection being zero. While we do not make this assumption, we do assume that the unpredictable responses to selection are independent across replicates. However, when applying the theory to data we assume that the within-replicate (residual) (co)variances are driven by drift only (rather than an unpredictable response to selection) and that the drift (co)variances are equal to their expected values (conditional on **L**_0_). In reality, the within-replicate covariances are likely larger, due to the unpredictable response to selection. This assumption might not be too severe if the unpredictable response to selection results in within-replicate (co)variances that are proportional to those under pure drift, although the residual variance may be higher than expected.
- In the absence of a recombination map, Assumption K may also be applied in our method.
- If **L**_0_ cannot be partitioned into gametic-phase and nongametic-phase contri-butions, assumption F might also be made by substituting **L**_0_ for 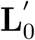 in 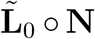 and 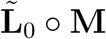.
- By relaxing some of the more extreme assumptions of B&C we are instead forced into making assumptions about the mean and covariance structure of the average effects of projected loci when making inferences.

### Multilocus simulations

To validate our method and inference approach, we performed multilocus simulations using msprime (version 1.2.0) (Kelleher *et al*., 2016) and SLiM (version 4.2.2) (Haller and Messer, 2023). The code for the simulations and the downstream analyses performed in R (version 4.3.3) is available on GitHub (https://github.com/manas-ga/Va_simulations) and includes the R library, Vw. We first describe the structure of the simulations in general terms and then discuss the specifics of our parameter choices in the context of the scaling used (see ‘Simulation parameters’ below). While these simulations do incorporate dominance effects, it is important to acknowledge that they do not include epistatic interactions between loci. They also assume random mating and non-overlapping generations. Furthermore, it is also important to note that genetic variation in fitness in our simulations was maintained, primarily, by drift-recombination-mutation-selection equilibrium, and our simulations do not capture other mechanisms of maintenance of genetic variation for fitness such as balancing selection or frequency-dependent selection.

We simulated a single, contiguous 1 million base-pair long genomic region. Our simulations had two distinct phases: the first phase simulated the history of an ancestral population and the second phase simulated a typical evolve and resequence experiment with independent replicate experimental populations derived from a base population drawn from the ancestral population.

### Model for fitness

We used an additive model across loci for log absolute fitness, *log*(*W*), as in Buffalo and Coop (2019). This choice was motivated by the fact that the variance in *log*(*W*) is approximately equal to the variance in relative fitness (see Appendix 1 in Lynch and Walsh (1998)). Across-locus additivity implies that the genotypic value of log fitness for individual *k* is equal to 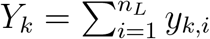, where *y*_*k,i*_ represents the genotypic contribution made by locus *i* to the log absolute fitness of individual *k*. Within loci, we employed the classical quantitative genetic fitness scheme (pp. 67 in Lynch and Walsh (1998)) adapted for *proportions* (*c*_*k,i*_’s) (as opposed to counts) of reference alleles within individuals:

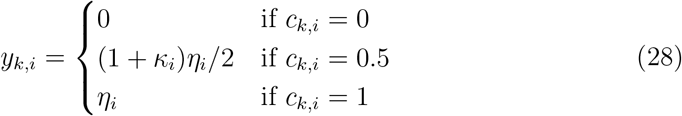

where *κ* determines the degree of dominance. Note that *κ* = 0 implies additivity and *κ* = −1 complete recessivity. The average effect for log absolute fitness at locus *i* is then given by 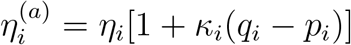 (see Equation 4.10b in Lynch and Walsh (1998)). The average effects for (relative) fitness (*α*’s) are well approximated by the *η*^(*a*)^’s when they are small in magnitude, but higher-order approximations are required when the *η*^(*a*)^’s are large (see Supplementary information S8). To obtain the log fitness of each individual we added a noise term drawn from a standard normal distribution (mean = 0, variance = 1) to the genotypic value. The absolute fitness of each individual was then obtained by exponentiating the individual’s log fitness.

We sampled the *η*’s from a distribution comprising a weighted mixture of three distributions: (1) a point mass at *η* = 0 representing neutral mutations, (2) a reflected gamma distribution with shape = 0.3 and scale = *η*_*scale*_ (typically 0.066, unless specified otherwise) representing deleterious mutations, and (3) a gamma distribution with shape = 0.3 and scale = *η*_*scale*_ (typically 0.066, unless specified otherwise) representing beneficial mutations. Thus, for both deleterious and beneficial mutations, the mean absolute *η* (i.e. *E*[|*η*|]) (scale *×* shape) was, typically, 0.02. The ratio of the frequency of beneficial to deleterious mutations, and *η*_*scale*_ varied among simulations. Below, we refer to the distribution of non-neutral *η*’s (i.e. omitting mixture component (1) described above) as the ‘distribution of fitness effects’ (DFE), but note that this is in fact the distribution of non-zero effects on log fitness.

### Phase 1 (‘history phase’): Simulating the history of an ancestral population

We first used a neutral coalescent simulation implemented in msprime (Kelleher *et al*., 2016) to construct genealogies for 2,500 diploid genomes (i.e. *N*_*e*_ = 2,500). To initialise a (non-equilibrium) set of selected loci, we then simulated mutations at a rate *µ*_*msp*_, using the pyslim package to attach fitness effects (*η*) drawn randomly from the non-neutral part of the distribution described above. Since derived mutations are rare, and the DFE is predominantly deleterious, this will generate a positive relationship between the reference allele frequency and the fitness effects *η*’s (and therefore the *α*’s, i.e. the average effects for relative fitness), such that 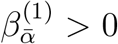 but there should be no relationship between genetic diversities and the *η*’s (i.e. 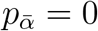). We implemented the fitness model described above by using SLiM’s (Haller and Messer, 2023) default recipe for polygenic selection (in the Wright-Fisher mode). SLiM code for implementing dominance effects in a quantitative genetic framework was adapted from Schaal *et al*. (2022). To reach mutation-selection-drift balance, we then let this population of 2,500 individuals evolve forward in time with selection for 25,000 generations. Non-neutral mutations drawn from the same DFE were allowed to occur in this period at a rate given by *µ*_*SLiM*_ . At generation 25,000, as the alleles reach mutation-selection-drift balance, we expect 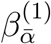 to have become more positive and 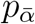 to have become more negative, better reflecting a real population undergoing selection.

In generation 25,000, we sampled *N*_0_ diploid individuals (typically 1000, unless specified otherwise) from this population, which then go on to become the base population in the next phase of the simulation. At this stage, we generated complete genomes for the *N*_0_ individuals in the base population by using pyslim to add neutral mutations to the tree sequence recorded so far. To obtain our target number of loci, *n*_*L*_ (typically 65,000, unless specified otherwise), we set the neutral mutation rate to be 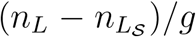 where 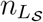 is the number of non-neutral segregating sites already present and *g* is the total branch length of the recorded tree sequence. We recorded the phased genotype of each parent at each locus, allowing us to construct **L**_0_ and 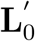.

### Phase 2 (‘experiment phase’): Simulating an evolve and resequence experiment

In the second phase of our simulations, again implemented in SLiM, we first allowed the base population to undergo one round of reproduction without selection to establish replicate experimental populations (typically 1,000 individuals in each of 10 replicates, unless specified otherwise). Next, we allowed each of these populations to evolve forward in time (typically three generations, unless specified otherwise) with selection as in the history phase. Since our goal was to estimate *V*_*A*_ in the base population, we restricted our analyses only to the set of loci segregating in the base population. Any new mutations in subsequent generations would, in all likelihood, occur at loci outside this set. Therefore, we did not simulate new mutations during the experiment phase. For each of the independent replicate populations, we recorded the genome-wide vector of allele frequencies in each generation of the experiment.

### Simulation parameters

#### Varying the true levels of V_A_

We varied the true (i.e. simulated) levels of *V*_*A*_ from *ca*. 0.01 to 0.1 by varying the number of non-neutral segregating sites 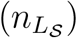 in the base population using a range of rates of non-neutral mutations in the history phase in both msprime (*µ*_*msp*_) and SLiM (*µ*_*SLiM*_). For example, in simulations with *E*[|*η*|] = 0.02 (i.e. *η*_*scale*_ = 0.066) and no beneficial mutations, we varied *µ*_*msp*_ between 5.56 *×* 10^−9^ and 5.56 *×* 10^−8^ and *µ*_*SLiM*_ between 5.56*×*10^−7^ and 5.56*×*10^−6^, which resulted in *n*_*L*_ varying between *ca*. 10,000 and 64,000 in the base population (for other scenarios see Table 1). We set *µ* _*msp*_ to be an order of magnitude smaller than *µ*_*SLiM*_ in most simulations because otherwise all individuals would have had a fitness of zero at the end of the coalescent part of the history phase. This extreme mutation load arises because deleterious alleles, having previously evolved under neutrality, would segregate at fairly high frequencies. However, the choice of *µ*_*msp*_ is unlikely to significantly affect the composition of the base population in most simulations, since the non-neutral genetic diversity of the base population of the experiment phase was primarily determined by the driftrecombination-mutation-selection equilibrium reached over the course of the 25,000 generation long forward simulation, and therefore primarily dependent on *µ*_*SLiM*_ .

#### Number of segregating sites (n_L_)

The requirement for handling large *n*_*L*_ × *n*_*L*_ matrices (e.g.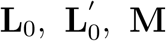, and **N**, etc.) imposed an upper bound of around 70,000 on *n*_*L*_ to permit the analysis of simulations in parallel. Under most scenarios the entire target range of simulated *V*_*A*_ (i.e. approximately 0.01 to 0.1) could be achieved without 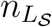 exceeding 65,000. Therefore, in general, we set *n*_*L*_ to 65,000 (but see ‘Simplified simulations’ below). However, under a small minority of scenarios – for example, at higher map lengths in the history phase (Figure 5B) or under strong dominance (Figure 3) – generating a larger *V*_*A*_ required significantly more than 65,000 non-neutral segregating sites. Consequently, in such cases, we had to discard simulations in which 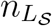 was greater than 65,000 which generally happened at the higher mutation rates and, therefore, the higher levels of *V*_*A*_.

#### Map length of the simulated region

There are two recombination-related parameters likely to influence the performance of our method: (1) the *V*_*A*_ per map length, and (2) the density of segregating sites per unit map length. Since our simulations were scaled down in an important way – namely, they assumed that *V*_*A*_ was limited to only about 65,000 segregating sites as opposed to millions of sites in real populations – it was not possible to simultaneously set parameters (1) and (2) close to their realistic values. Therefore, we varied the effective map length from 0.001 to 2 morgans which spans scenarios where either the total map length of the simulated region is comparable to a typical *D. melanogaster* autosome (∼ 0.5 morgan) or the density of segregating sites is comparable (∼ 0.065 morgan). Exact settings are detailed in the ‘Reference parameters’ section below. The typical *D. melanogaster* map-length was determined from an effective crossover rate of 10^−8^ (Comeron *et al*., 2012; Wang *et al*., 2023) and a 50Mb chromosome. The typical site-density was determined under the assumption that there are one million segregating sites per morgan, which is reasonably consistent with a recent evolveand-resequence study employing *D. melanogaster* (Bitter *et al*. (2024): 827,200 and 893,465 sites on Chromosome 2 and 3 respectively).

Since we only simulated a population of 2,500 individuals in the history phase, which is ∼500 times smaller than the estimate of *N*_*e*_ in *D. melanogaster* (Campos *et al*., 2013; Campos and Charlesworth, 2019), we scaled our map-length upwards by a factor of 500 to achieve the target effective map-length described above (Campos and Charlesworth (2019) but see Dabi and Schrider (2025) and Ferrari *et al*. (2025) for potential issues). On the other hand, since the experiment phase simulates a realistic evolve-and-resequence experiment with *N*_*e*_ *≈* 1, 000, this scaling was not applied to map lengths in the experiment phase (see ‘Reference parameters’ below). In what follows, the map-lengths implemented in the simulations are reported rather than the target effective map-lengths.

#### Degree of dominance

In the simulations with dominance effects (i.e. *κ* ≠ 0) we always modelled deleterious alleles to be recessive. In practical terms, this meant that *κ*_*i*_ was set to be positive when *η*_*i*_ was positive, and *κ*_*i*_ was set to be negative when *η*_*i*_ was negative. For simplicity, in a given simulation, we assumed |*κ*_*i*_ | to be the same across all loci, although the actual dominance deviation, *d*_*i*_ = *κ*_*i*_*η*_*i*_ would be locus-specific. We simulated two different levels of |*κ*| : (1) |*κ*| = 0.5, and (2) |*κ*| = 0.9. These correspond to *h* = 0.25 and *h* = 0.05, respectively, when the heterozygous fitness is 1 − *hs*. While *h* = 0.25 is consistent with the theoretical expectation from a fitness landscape based model (Manna *et al*., 2011), *h* = 0.05, where every deleterious allele is almost completely recessive is probably extreme. Since full simulations with dominance were up to two orders of magnitude slower than their additive counterparts, we switched on dominance effects only in the last 5,000 generations of the history phase (i.e. from generation 20,001) as well as in the experiment phase. Relative to additive simulations, full simulations with dominance also required significantly more segregating sites to generate a given level of *V*_*A*_. To generate sufficient *V*_*A*_ (i.e. between *ca*. 0.01 and 0.1) while keeping 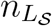 below 67,500, we simulated deleterious mutations from a gamma distribution with a slightly larger mean effect on log fitness (i.e. E[|*η*|] = 0.03) in the history phase and set |*κ*| to either 0.5 or 0.75. To test our method against extreme dominance (i.e. |*κ*| = 0.9), we performed additional full simulations in which we simulated the entire history phase under additivity, and switched on dominance effects only in the experiment phase. While the resulting genetic composition was far from the expected equilibrium, deleterious alleles were expected to be segregating at substantially lower frequencies than in the simplified simulations.

#### Reference parameters

Our aim was to investigate the sensitivity of our method to the following parameters: (1) map length in the history phase (0.5 morgan, 5 morgans, 50 morgans, 250 morgans), (2) map length in the experiment phase (0.01 morgan, 0.2 morgan, 2 morgans), (3) number of replicate populations in the experiment phase (3, 5, 10), (4) the population size of each of the replicate populations (100, 500, 1,000), (5) number of generations over which allele frequency changes were recorded in the experiment phase (1, 3, 5), (6) the ratio of the rates of beneficial to deleterious mutations (0%, 0.02 %, 2%), and (7) the mean of the gamma distribution from which non-neutral *η*’s were sampled, *E*[| *η*|] (0.02, 0.03, 0.06), and (8) the degree of dominance, *κ* (0, 0.5, 0.9).

Rather than vary all parameters in a fully factorial design, we selected a reference parameter set and explored the sensitivity of the method by changing each parameter in turn. The reference parameter set was (1) a map length of 0.5 morgan in the history phase (2) a map length of 2 morgans in the experiment phase, (3) 10 replicate populations in the experiment phase, (4) a population size of 1,000 in each of the replicate populations in the experiment phase, (5) 3 generations over which allele frequency changes were recorded in the experiment phase, (6) no beneficial mutations in the history phase, (7) *E*[| *η*|] = 0.02, and (8) complete additivity (i.e. *κ* = 0). The reference parameter set was chosen to best reflect evolve-and-resequence experiments using *D. melanogaster*.

Note that we varied *E*[| *η*|] by using three different values (0.066, 0.09, and 0.2) for the scale of the gamma distribution (*η*_*scale*_) from which the *η*’s for non-neutral mutations were drawn while keeping the shape parameter fixed to 0.3. This meant that the coefficient of variation was fixed to 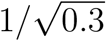.

### Simplified simulations

In addition to the full simulations described above, we performed a set of simplified, proof-of-principle simulations to test the logic of our method. These simulations were different from the full simulations in three ways:

1. The history phase was highly abbreviated – the forward simulation implemented in SLiM lasted only a single generation without any selection. This meant that the ancestral population – from which replicate experimental population were founded – had evolved entirely under neutrality.
2. Rather than attach *η*’s to the derived alleles generated with msprime, we attached the *η*’s randomly with respect to either of the two alleles segregating at the locus. Amendments (1) and (2) generate a scenario where there is no relationship between allele frequency and *η*: both 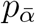 and 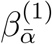 are expected to be zero.
3. Given that the ancestral population had evolved without selection, these simplified simulations had considerably higher genetic diversities compared to the full simulations. This meant that fewer segregating non-neutral sites 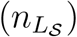 were required to achieve true levels of *V*_*A*_ between 0.01 and 0.1. Therefore, in these simulations, we set *n*_*L*_ to be 3,000, much lower than the 65,000 used for the full simulations. We achieved this by varying *µ*_*msp*_ between 3 *×* 10^−9^ and 2.35 *×* 10^−8^. *µ*_*SLiM*_ was set to 0 in these simulations. This resulted in 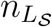 at the end of the history phase being between 160 and 1,902. We then added neutral mutations using a suitable rate as described above such that *n*_*L*_ was expected to be 3,000.

### Simulating pool-seq

To investigate the consequences of sampling noise around true allele frequencies in real data, we reanalysed the reference sets for full and simplified simulations with allele frequencies in the experiment phase obtained via simulated pool-seq. Specifically, we investigated the sensitivity of our approach to the pool-seq coverage (i.e. the expected number of reads spanning a given locus) and the degree of overdispersion in the number of individuals sampled. We assumed that the number of reads mapping to individual *k* followed a Poisson distribution with mean *λ*_*x,k*_, which in turn is sampled from a log-normal distribution with parameters *µ*_*x*_ and *V*_*x*_. Varying *V*_*x*_ allowed us to control the degree of heterogeneity in the probabilities with which individuals are sampled. For a given level of coverage and read length, we determined the expected number of reads mapping to an individual (*E*[*λ*_*x*_]). For a given value of *V*_*x*_ and read-length, using the expression for the mean of a lognormal distribution (*E*[*λ*_*x*_] = *exp*(*µ*_*x*_ + *V*_*x*_*/*2)), we chose *µ*_*x*_ to achieve the target coverage: *µ*_*x*_ = *log*(*E*[*λ*_*x*_]) − *V*_*x*_*/*2. We mapped reads to the genomes of various individuals in the population by sampling the starting positions for each read from a uniform distribution.

To select the length of the reads in these simulations, we first note that the expected number of segregating sites spanned by a 150 base-pair read in a recent evolve-and-resequence study (Bitter *et al*., 2024) (827,200 segregating sites on chromosome 2 which is ∼50 Mb in length) is approximately 2.5. To achieve this, we fixed the read length to be 800 while analysing simplified simulations (∼ 3,000 segregating sites) and 37 when analysing full simulations (∼ 65,000 segregating sites). We used four different combinations of coverage and *V*_*x*_: (1) coverage = 100x, *V*_*x*_ = 0; (2) coverage = 500x, *V*_*x*_ = 0; (3) coverage = 1000x, *V*_*x*_ = 0); (4) coverage = 1000x, *V*_*x*_ = *log*(2). Note that *V*_*x*_ = 0 corresponds to a situation where individuals are sampled without overdispersion.

### Comparison with B&C

To compare the precision and the accuracy of our approach to that of B&C, we performed additional simplified and full simulations. Although originally designed for the covariance in allele frequency change between multiple time points, B&C’s method can be readily adapted to allele frequency changes recorded over a single generation in multiple independent evolutionary replicates (see Equation S59 and Buffalo and Coop (2020)). We set simulation parameters to their reference values (see above) with a few modifications. First, to make comparisons of biases clearer, we reduced noise by employing 50 replicate populations, and second, allele frequency change in the experiment phase was recorded over a single generation. We also ran these simulations at four different levels of map length in the history phase: 0.5 morgan, 5 morgans, 50 morgans, and 100 morgans.

To investigate the consequences of Assumptions B, Ib, and N, we implemented B&C’s method using six different approaches. For clarity, we label the six approaches using a code (see Table 2) that indicates with superscripts whether an assumption is required (+) or not required (0) for an approach to work. To test Assumption B, we recorded allele frequency changes at either all segregating sites (*B*^+^) or neutral sites only (*B*^0^). To test Assumption Ib, we used either the average LD between all sites (*Ib*^+^) or the average LD between selected sites with neutral sites (*Ib*^0^). And finally, we either divided the (co)variance in allele frequency change throughout by twice the average genetic diversity (*N* ^+^; analogous to Equation 16 in B&C) or used the (co)variance of weighted allele frequency change, where the weights are the inverse of the square root of twice the genetic diversity (*N* ^0^).

**Table 2:** We applied B&C’s method to our simulations using six different approaches. The requirement of Assumptions B, Ib, and N is indicated using superscripts (“+” when required and “0” when not required). We used allele frequency change (Δ**p**) at either all segregating sites (Assumption B required) or at neutral segregating sites only (Assumption B not required). We computed average LD using either the LD between all segregating sites (Assumption Ib required) or the LD between selected and neutral sites (Assumption Ib not required). We used either the (co)variance in Δ**p** divided throughout by twice the average genetic diversity (Assumption N required) or the (co)variance in weighted Δ**p** (i.e. 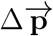) where the weights are the square roots of twice the genetic diversity at each site (Assumption N not required).

| **Approach** | **$\Delta p$** | **Average LD between** | **$Cov(\Delta \mathbf{p}_m, \Delta \mathbf{p}_n)$** |
| --- | --- | --- | --- |
| $B^+Ib^+N^+$ | all sites | all sites | divided throughout by $(\bar{p}_m\bar{q}_m + \bar{p}_n\bar{q}_n)/2$ |
| $B^0Ib^+N^+$ | neutral sites | all sites | divided throughout by $(\bar{p}_m\bar{q}_m + \bar{p}_n\bar{q}_n)/2$ |
| $B^0Ib^0N^+$ | neutral sites | selected & neutral sites | divided throughout by $(\bar{p}_m\bar{q}_m + \bar{p}_n\bar{q}_n)/2$ |
| $B^+Ib^+N^0$ | all sites | all sites | replaced by $Cov(\Delta \vec{\mathbf{p}}_m, \Delta \vec{\mathbf{p}}_n)$ |
| $B^0Ib^+N^0$ | neutral sites | all sites | replaced by $Cov(\Delta \vec{\mathbf{p}}_m, \Delta \vec{\mathbf{p}}_n)$ |
| $B^0Ib^0N^0$ | neutral sites | selected & neutral sites | replaced by $Cov(\Delta \vec{\mathbf{p}}_m, \Delta \vec{\mathbf{p}}_n)$ |

**Table 3:** Notation.

| Symbol | Description |
| --- | --- |
| $\mathbf{B}$ | Diagonal matrix of standard deviations for the $c$'s |
| $c_{k,i}$ | Number of reference alleles at locus $i$ for individual $k$ divided by 2. |
| $C_A(t \rightarrow \tau)$ | Additive genetic covariance for relative fitness between generation $t$ and $\tau$ for a population with genetic structure equal to that in generation $t$. |
| $C_a(t \rightarrow \tau)$ | Additive genic covariance for relative fitness between generation $t$ and $\tau$ for a population with genetic structure equal to that in generation $t$. |
| $\mathbf{D}_2$ | Diagonal matrix of square-rooted eigenvalues of $\mathcal{D}$ |
| $\mathcal{D}_m$ | Matrix that gives the covariance in allele frequency changes between generation $t_m$ and $\tau_m$ in replicate $m$ due to drift. |
| $\mathcal{E}_m$ | Matrix that gives the covariance in allele frequency change estimation errors in replicate $m$. |
| $\mathbf{F}_{t\tau}$ | $\mathbf{W}_{t\tau}$ but with the diagonals set to zero. |
| $g$ | Total branch length of the tree sequence recorded in the history phase of the simulations |
| $\mathbf{H}_{t\tau}$ | $\mathbf{W}_{t\tau}$ but with the off-diagonals set to zero. |
| $\mathbf{L}_{t,m}$ | Covariance matrix of $c$'s at time $t$ in replicate $m$. |
| $\mathbf{L}'_{t,m}$ | Covariance matrix of $c$'s at time $t$ in replicate $m$ due to being on the same gametic contribution. |
| $\mathbf{L}''_{t,m}$ | Covariance matrix of $c$'s at time $t$ in replicate $m$ due to being on different gametic contributions. |
| $\Delta \mathbf{L}'_{t,m}$ | The stochastic change in $\mathbf{L}'$ between time zero and $t$ in replicate $m$. |
| $\tilde{\mathbf{L}}_0$ | Weighted sum of $\mathbf{L}'_0$ and $\mathbf{L}''_0$ with weights for element $ij$ being 1 and $z_{0,ij}/\zeta_{0,ij}$ respectively. |
| $\mathcal{L}_{t,m}$ | A matrix that gives, when post-multiplied by $\boldsymbol{\alpha}$, the predictable change in allele frequency due to selection between generation $t_m$ and $t_m + 1$ in replicate $m$. |
| $\mathcal{L}_m$ | A matrix that gives, when post-multiplied by $\boldsymbol{\alpha}$, the predictable change in allele frequency due to selection between generation $t_m$ and $\tau_m$ in replicate $m$. |
| $\mathbf{M}_{t,m}$ | Matrix of weights for $\tilde{\mathbf{L}}_0$ that gives the covariance in allele frequency changes due to drift between time $t > 0$ and $t + 1$ in replicate $m$. |
| $\mathbf{M}_m$ | Matrix of weights for $\tilde{\mathbf{L}}_0$ that gives the covariance in allele frequency changes due to drift between time $t_m > 0$ and $\tau_m$ in replicate $m$ |
| $n_L$ | Number of loci |
| $n_{LS}$ | Number of non-neutral segregating sites |
| $N_{t,m}$ | Census population size at time $t$ in replicate $m$ |
| $N_{e_{t,m}}$ | Variance effective population size at time $t$ in replicate $m$ |
| $N_{E_{t,m}}$ | Variance effective population size at time $t$ in replicate $m$ ignoring the impact of linked-selection |
| $\mathbf{N}_{t,m}$ | Matrix of weights for $\tilde{\mathbf{L}}_0$ that gives the expected $\mathbf{L}$ at time $t > 0$ in replicate $m$. |
| $\mathbf{N}_m$ | Matrix of weights for $\tilde{\mathbf{L}}_0$ that gives the sum of the expected $\mathbf{L}$ from time $t_m > 0$ to $\tau_m$ in replicate $m$. |
| $\mathcal{N}$ | Used as a subscript to indicate the set of neutral loci. |
| $O_{ij}$ | Number of pool-seq reads spanning sites $i$ and $j$. If $i = j$, $O_{ij}$ is the coverage. |
| $p_{\bar{\alpha}}$ | Parameter that takes $\mathbf{L}_0$ to some power |
| $\mathbf{p}_{t,m}$ | Vector of reference allele frequencies at time $t$ in replicate $m$. |
| $\Delta \mathbf{p}_{t,m}$ | Vector of reference allele frequency changes between time $t$ and $t + 1$ in replicate $m$. |
| $\Delta \mathbf{p}_m$ | Vector of reference allele frequency changes between time $t_m$ and $\tau_m$ in replicate $m$. |
| $\mathbf{P}$ | Projection matrix for allele frequencies. |
| $\mathbf{q}_{t,m}$ | Vector of alternate allele frequencies at time $t$ in replicate $m$. |
| $\mathbf{Q}_m$ | Matrix that is proportional to the covariance in allele frequency change estimation errors in replicate $m$. |
| $r_{ij}$ | Recombination rate between loci $i$ and $j$. |
| $R_{t,ij}$ | Correlation in allele count between locus $i$ and $j$ at time $t$ (Note the use of the uppercase to distinguish from the recombination rate $r$). |
| $\mathbf{R}_+$ | Matrix of recombination probabilities. |
| $\mathbf{R}_-$ | Matrix of non-recombination probabilities. |
| $\mathbf{R}$ | Correlation matrix of the $c$'s. |
| $\mathbf{S}_{\bar{\alpha}}$ | Sampling covariance matrix for the parameters of the regression on the mean average effects, $\beta_{\bar{\alpha}}$. |
| $\mathcal{S}$ | Used as a subscript to indicate the set of selected loci. |
| $t_m$ | Time at which allele frequencies are first measured in replicate $m$. |
| $\mathbf{U}$ | Eigenvectors of $\mathbf{L}_0$. |
| $\mathbf{U}_{\mathbf{L}}$ | Eigenvectors of $\mathbf{L}_0$ with non-zero eigenvalues. |
| $\mathbf{U}_2$ | Eigenvectors of $\mathcal{D}$ |
| $\mathcal{U}_m$ | Matrix that gives the covariance in allele frequency changes between generation $t_m$ and $\tau_m$ in replicate $m$ due to the unpredictable response to selection. |
| $V_a(t)$ | Additive genic variance for relative fitness at time $t$ |
| $V_A(t)$ | Additive genetic variance for relative fitness at time $t$ |
| $V_{\bar{A}}(t)$ | Additive genetic covariance between replicate/time-points for relative fitness in a population with genotypic composition equal to that at time $t$. |
| $V_x$ | Variance of the normal distribution from which $\log(\lambda_x)$'s are sampled |
| $\mathbf{V}_{\bar{\alpha}}$ | Covariance matrix for the mean average effects. |
| $w$ | Relative fitness |
| $W$ | Absolute fitness |
| $\mathbf{W}_{t\tau}$ | A matrix with the $ij^{th}$ element equal to $R_{t,ji}R_{\tau,ki}(b_{\tau,i}/b_{t,i})$ |
| $\mathbf{X}$ | Design matrix for the regression of the mean average effects on $\mathbf{p}_0 - \mathbf{q}_0$. |
| $y_{k,i}$ | Genotypic contribution made by locus $i$ to the log absolute fitness ($\log(W)$) of individual $k$ |
| $Y_k$ | Genotypic value of log fitness for individual $k$ |
| $z_{t,ij}$ | $(1 - r_{t,ij})(1 - \frac{1}{2N_{e_t}})$ |
| $\boldsymbol{\alpha}_{t,m}$ | Vector of average effects for relative fitness at time $t$ in replicate $m$. |
| $\bar{\alpha}$ | Vector of mean average effects for relative fitness. |
| $\Delta\alpha_{t,m}$ | Vector of deviations of the average effects for relative fitness at time $t$ in replicate $m$ from the global mean. |
| $\beta_{\bar{\alpha}}^{(1)}$ | Slope of the regression of the mean average effects on $\mathbf{p}_0 - \mathbf{q}_0$. |
| $\zeta_{t,ij}$ | $r_{t,ij}(1 - \frac{1}{2N_{e_t}})$ |
| $\eta_i$ | Difference between the genotypic contributions to $\log(W)$ by the reference and non-reference homozygotes at locus $i$ |
| $\eta_i^{(a)}$ | Average effect for $\log(W)$ at locus $i$ |
| $\eta_{scale}$ | Scale of the gamma distribution from which $\eta$'s were sampled in the history phase of the simulations |
| $\kappa_i$ | Degree of dominance at locus $i$ for log absolute fitness. |
| $\lambda_{x,k}$ | Mean of the Poisson distribution from which the number of reads mapping to individual $k$ are sampled while simulating pool-seq |
| $\mu_{msp}$ | Mutation rate for non-neutral mutations in the coalescent part of the history phase of the simulations |
| $\mu_{SLiM}$ | Mutation rate for non-neutral mutations in the forward part of the history phase of the simulations |
| $\mu_x$ | Mean of the normal distribution from which $\log(\lambda_x)$'s are sampled |
| $\boldsymbol{\mu}_{\bar{\alpha}}$ | Vector of expected values for the mean average effects. |
| $\sigma_{\bar{\alpha}}^2$ | Proportionality constant that relates $\mathbf{V}_{\bar{\alpha}}$ to $\mathbf{L}_0^{p_{\bar{\alpha}}}$. |
| $\sigma_o^2$ | Parameter for scaling sampling (co)variances for allele frequencies: values greater than 1 indicate overdispersion. |
| $\tau_m$ | Time at which allele frequencies are finally measured in replicate $m$. |
| $\phi_{t,\tau}$ | The ratio of genetic diversity in generation $\tau$ to genetic diversity in generation $t$ assumed constant across all selected loci. |
| → | Used above a symbol to indicate it is on the projected space. |
| $\wedge$ | Used above a symbol to indicate an estimate. |

## Results

### Simplified simulations

We begin by discussing results from the simplified simulations in which we expect there to be no relationship between allele frequencies and *α*’s. Under our reference parameter set (no dominance, map length in history = 0.5 morgan, map length in the experiment = 2 morgans), our method provided precise and unbiased estimates of *V*_*A*_ throughout the simulated range (0.01– 0.1) (Figure 1a). Furthermore, the estimates of 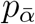 and 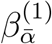 were centred around 0 as expected (Figure 1b-c), and the residual variance was marginally above 1 (Figure 1d). Next, we investigated how our estimates of *V*_*A*_ were affected by changing (1) map length of the genomic region being simulated, (2) population size of each replicate population in the evolve and resequence experiment, (3) number of replicate populations, and (4) number of generations over which allele frequency changes were recorded in the experiment. Estimates of *V*_*A*_ remained unbiased as the map length in the experiment became smaller, although estimates became noisier, particularly at higher values of *V*_*A*_ (Figure 2a). As expected, estimates also became noisier as the number of individuals, replicate populations, or generations became smaller (Figure 2b-d). However, estimates appeared upwardly biased when the number of replicate populations or generations was small (Figure 2c-d). Estimates of *V*_*A*_ from simplified simulations incorporating dominance effects in the fitness model were generally precise, but exhibited a small upward bias when dominance effects were large in magnitude (Figure 3a). As the degree of dominance increased, the estimates of 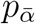 and 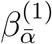 became increasingly more negative and the residual variance became larger (Figure 3b-d).

**Figure 1:**
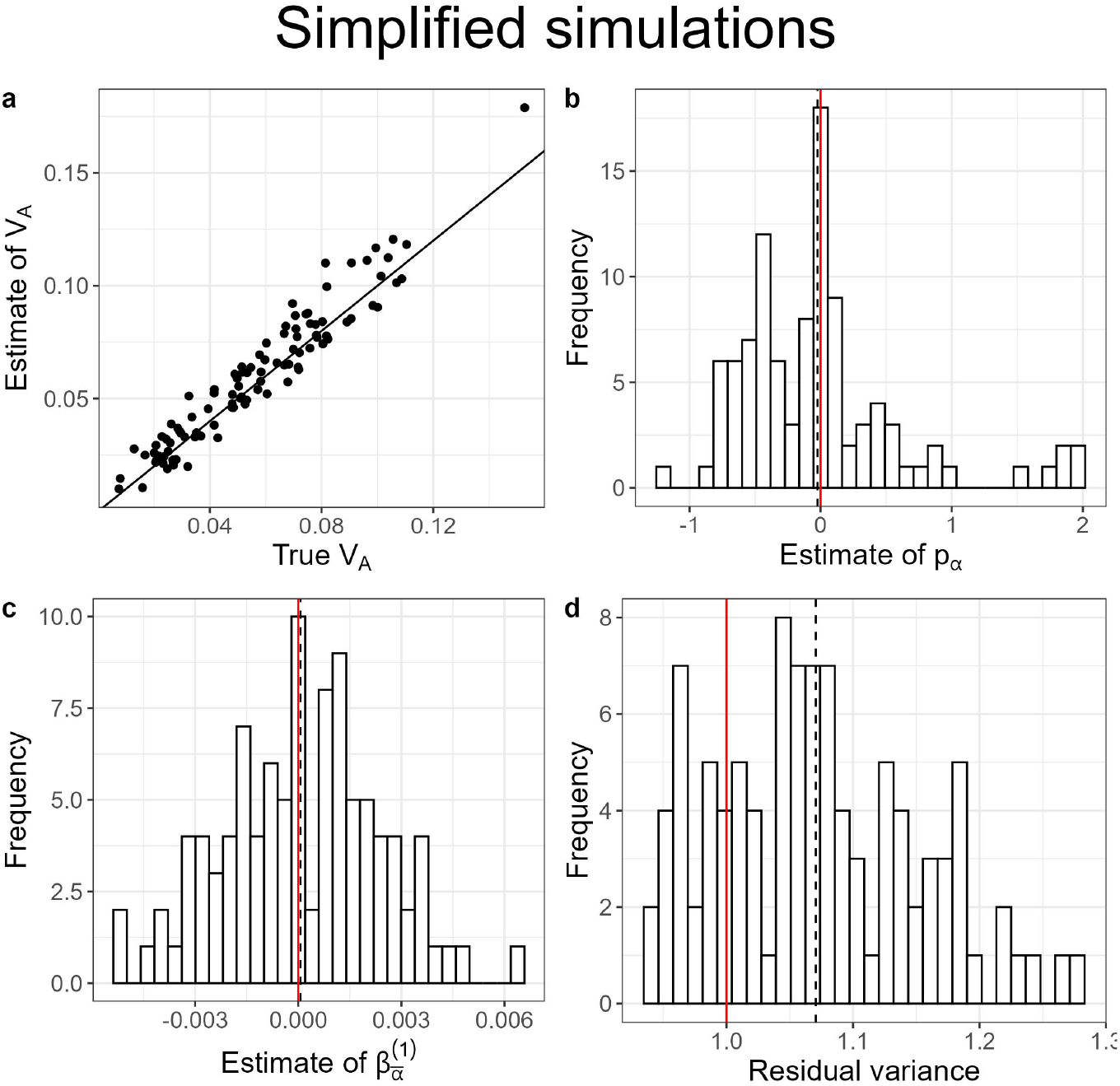
Results of simplified simulations (map length in the history phase = 0.5 morgan, map length in the experiment phase = 2 morgans, number of replicate populations = 10, population size = 1,000, number of generations = 3, the mean of the gamma distribution from which effect sizes for log absolute fitness were sampled for non-neutral mutations (*E*[|*η*|]) = 0.02, and no dominance (i.e. *κ* = 0)). (a) A scatter plot of estimates of *V*_*A*_ vs true values of *V*_*A*_. The solid black line indicates the 1:1 line. The inference of *V*_*A*_ was obtained by modelling the mean and the (co)variance of the average effects for relative fitness as 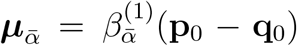 and 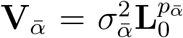, respectively. (b)-(d) Histograms of the estimates of 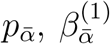, and the residual variance, respectively. The vertical red lines indicate null expectations (0 for 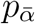 and 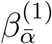, and 1 for the residual variance). The black dashed lines indicate the means of the respective distributions.

**Figure 2:**
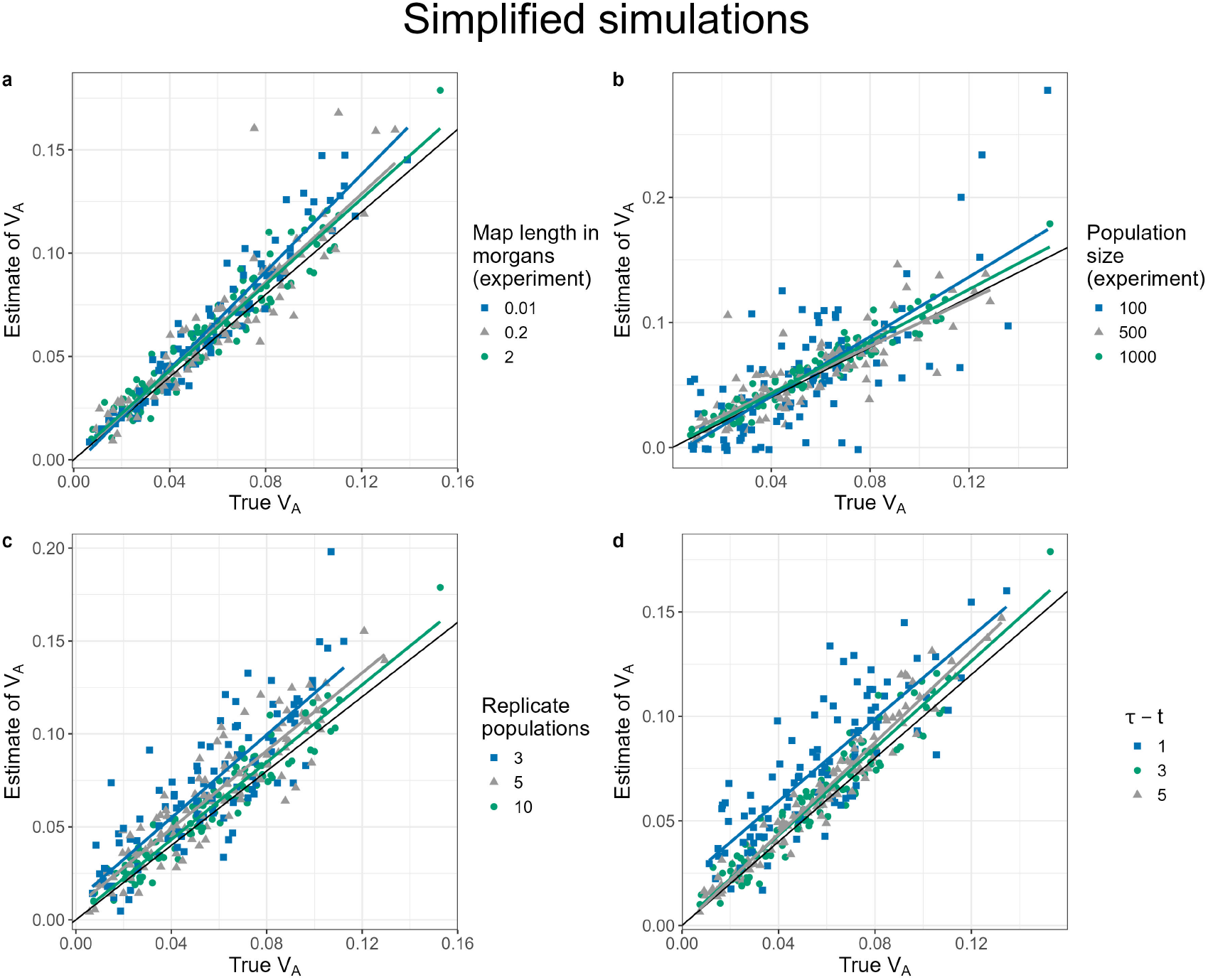
Scatter plots of estimates of *V*_*A*_ vs true values of *V*_*A*_ for simplified simulations at different levels of (a) map length in the experiment, (b) population size of each replicate population in the evolve and resequence experiment, (c) number of replicate populations, and (d) number of generations over which allele frequency changes were recorded in the experiment. In each case, other than the parameter to be varied, the other parameters were fixed at their default values: the mean of the gamma distribution from which effect sizes for log absolute fitness were sampled for non-neutral mutations (*E*[|*η*|]) = 0.02, map length in the history phase = 0.5 morgan, map length in the experiment phase = 2 morgans, number of replicate populations = 10, population size = 1,000, number of generations = 3, and no dominance (*κ* = 0)). The solid black line indicates the 1:1 line. The coloured lines represent regression lines for estimates of *V*_*A*_ vs true values of *V*_*A*_.

### Full simulations

In our full simulations we let the ancestral population evolve forward in time with selection for 25,000 generations before simulating the experiment. In these simulations, we expect the population to be at drift-recombination-mutation-selection equilibrium, such that *p*_*α*_ is negative and 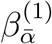 is positive. Results from our reference set (no dominance, map length in history = 0.5 morgan, map length in the experiment = 2 morgans) suggest that not only does our method provide precise and unbiased estimates of *V*_*A*_ (Figure 4a), but it does so by correctly estimating the signs of *p*_*α*_ and 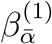 (Figure 4b-c). As before, estimates of *V*_*A*_ became slightly noisier at shorter map lengths during the experiment (Figure 5a). The quality of our estimates of *V*_*A*_ was not significantly affected by adding beneficial mutations during the history phase (Figure 5c). On the other hand, our estimates were downwardly biased at larger map lengths in the history phase (Figure 5b, also see Supplementary Figure 1) and when non-neutral loci had on average larger fitness effects (*E*[|*η* |]) (i.e. when *η*_*scale*_ was larger) (Figure 5d). The bias at larger *E*[|*η* |] was likely driven by the loss of additive genic variance at large-effect loci during the experiment (Supplementary Figure 2a-d).

When we simulated the history phase under additivity and switched on dominance only in the experiment phase, we obtained fairly reliable estimates of *V*_*A*_ (Supplementary Figure 4). However, switching on dominance in the last 5,000 generations of the history phase resulted in estimates that were significantly upwardly biased, especially when dominance effects were strong (Figure 3e). However, at the relatively low map length in the history phase (0.5 morgan) used in these simulations, the additive genetic variance for fitness was substantially lower than the additive genic variance (Supplementary Figure 5). This suggested the build up of unusually high negative linkage disequilibria between recessive deleterious alleles, generating pseudo-overdominance (Ohta and Kimura, 1970; Abu-Awad and Waller, 2023), a phenomenon typically observed in regions of extremely low recombination (Salson *et al*., 2025). At higher levels of map length in the history phase (5 morgans), our method provided unbiased estimates of *v*_*A*_ (Figure 3f).

**Figure 3:**
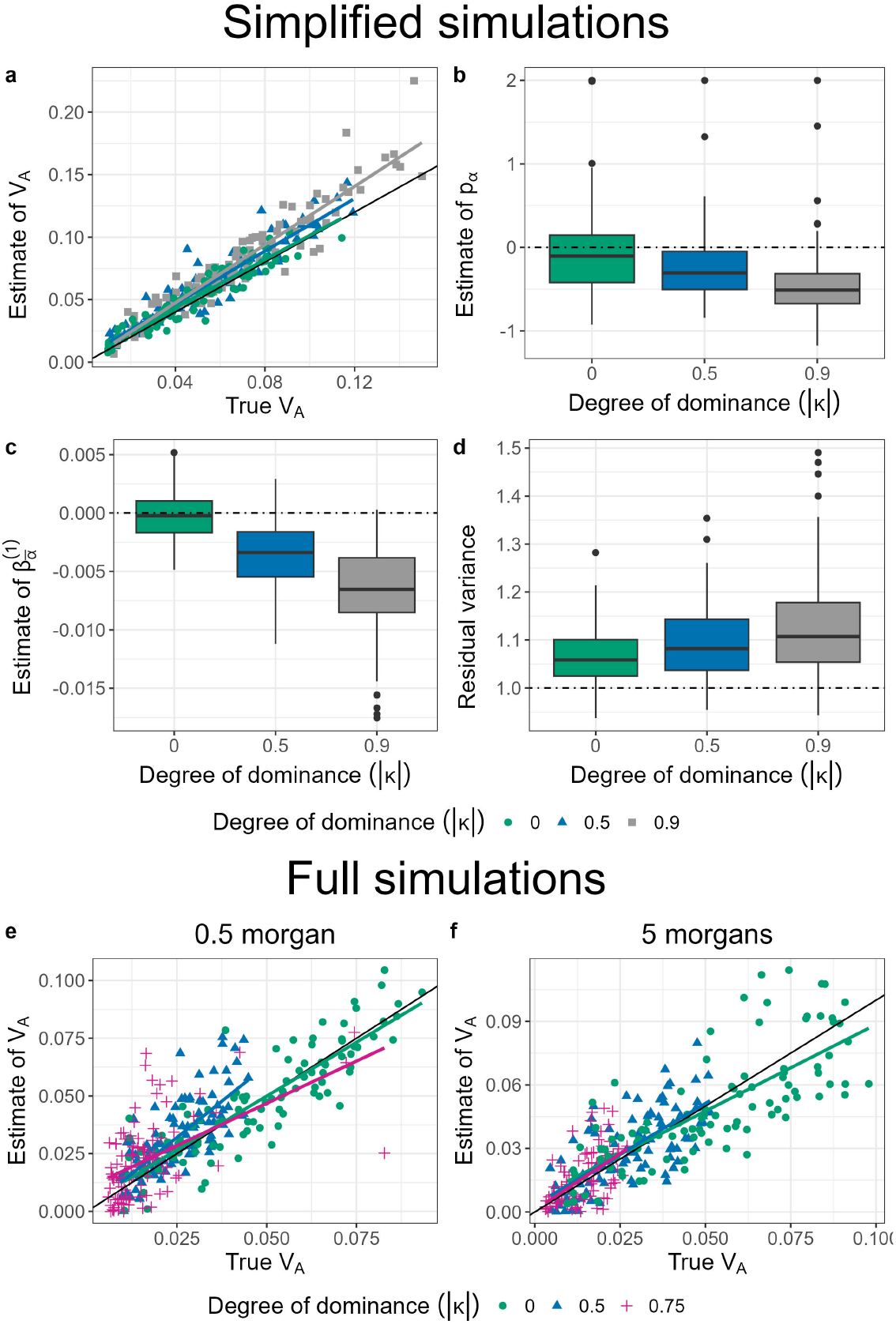
Results of simulations with map lengths in the experiment phase = 2 morgans, number of replicate populations = 10, population size = 1,000, number of generations = 3, and different degrees of dominance: *κ* = 0 (green), *κ* = 0.5 (blue), *κ* = 0.75 (magenta) and *κ* = 0.9 (grey). (a-d) are the results from the simplified simulations with the map length in the history phase = 0.5 morgan and the mean of the gamma distribution from which effect sizes for log absolute fitness are sampled for non-neutral mutations (*E*[|*η*|]) = 0.02. (a) A scatter plot of estimates of *V*_*A*_ vs true values of *V*_*A*_. The solid black line indicates the 1:1 line. The coloured lines represent regression lines for estimates of *V*_*A*_ vs true values of *V*_*A*_. (b-d) Estimates of model parameters 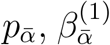, and the residual variance, respectively. (e-f) Scatter plot of estimates of *V*_*A*_ vs true values from full simulations (with *E*[|*η*|] = 0.03) when the map length in the history phase was either 0.5 morgan (e) or 5 morgans (f).

**Figure 4:**
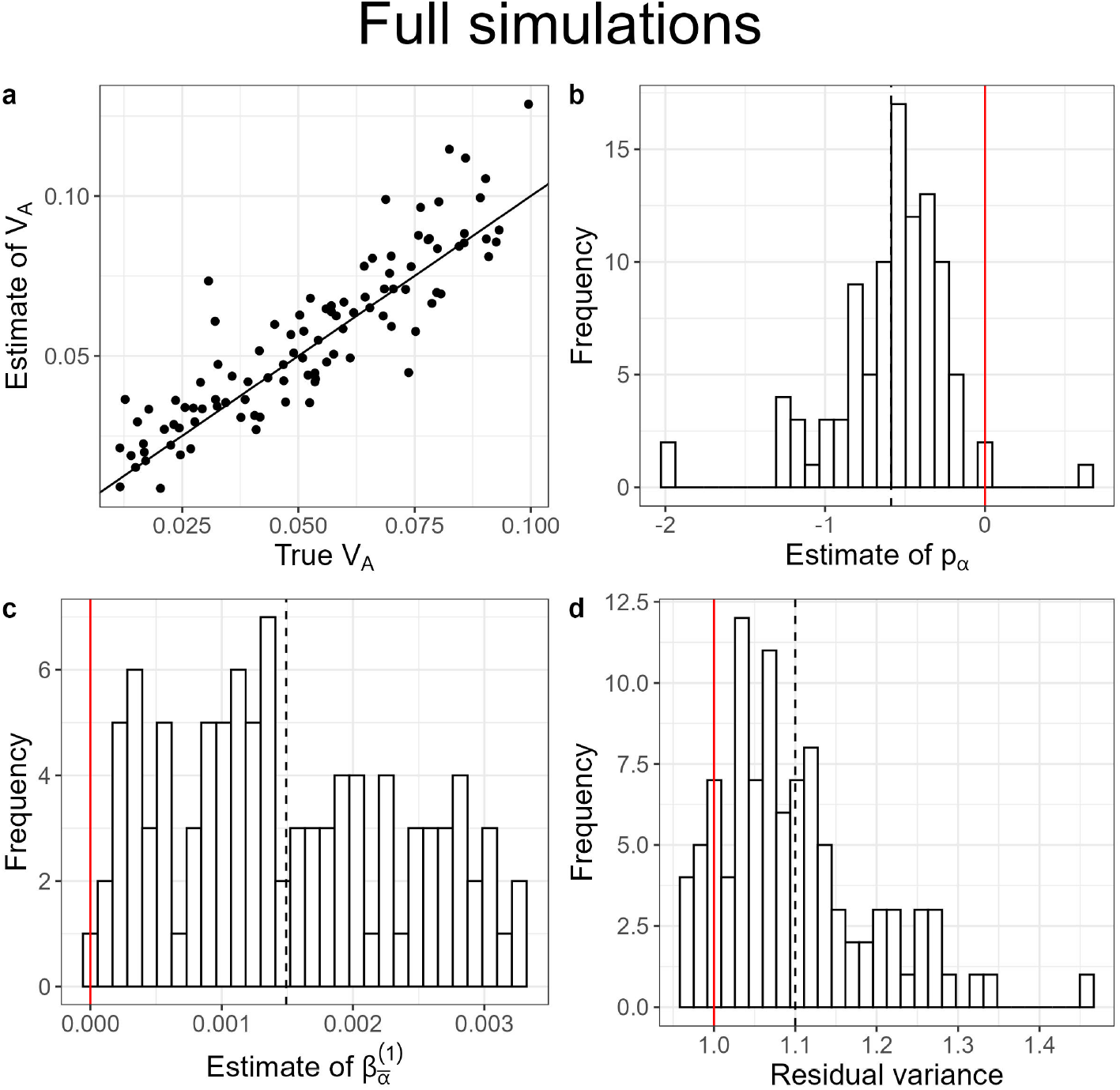
Results of full simulations with a burn-in phase of 25,000 generations (map length in the history phase = 0.5 morgan, map length in the experiment phase = 2 morgans, number of replicate populations = 10, population size = 1,000, number of generations = 3, the mean of the gamma distribution from which effect sizes for log absolute fitness were sampled for non-neutral mutations (*E*[|*η*|]) = 0.02, and no dominance (i.e. *κ* = 0)). (a) A scatter plot of estimates of *V*_*A*_ vs true values of *V*_*A*_. The solid black line indicates the 1:1 line. The inference of *V*_*A*_ was obtained by modelling the mean and the (co)variance of the average effects for relative fitness as 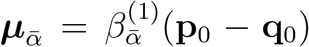 and 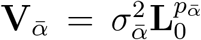, respectively. (b)-(d) Histograms of the estimates of 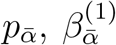, and the residual variance, respectively. The vertical red lines indicate null expectations (0 for 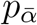 and 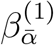, and 1 for the residual variance). The black dashed lines indicate the means of the respective distributions.

Finally, we re-analysed our standard set of full simulations in two different ways (Figure 5e-f). First, to accommodate cases in which phased genomes, and therefore 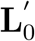, are unavailable, we assumed that 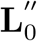 was 0 and set 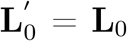. In other words, we assumed that 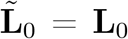. Our analyses suggest that this assumption leads to a slight downward bias in our estimates of *V*_*A*_ (Figure 5e). Second, to account for instances where *N*_*E*_ in the experiment phase is unknown, we replaced our estimate of *N*_*E*_ in the experiment phase by the number of individuals. Our analyses suggest that this does not affect our estimates of *V*_*A*_ in any way (Figure 5f), with the faster than expected drift being absorbed by an increased residual variance (Supplementary Figure 3): estimates of *N*_*E*_ can be obtained by dividing the assumed value of *N*_*E*_ by the estimated residual variance.

**Figure 5:**
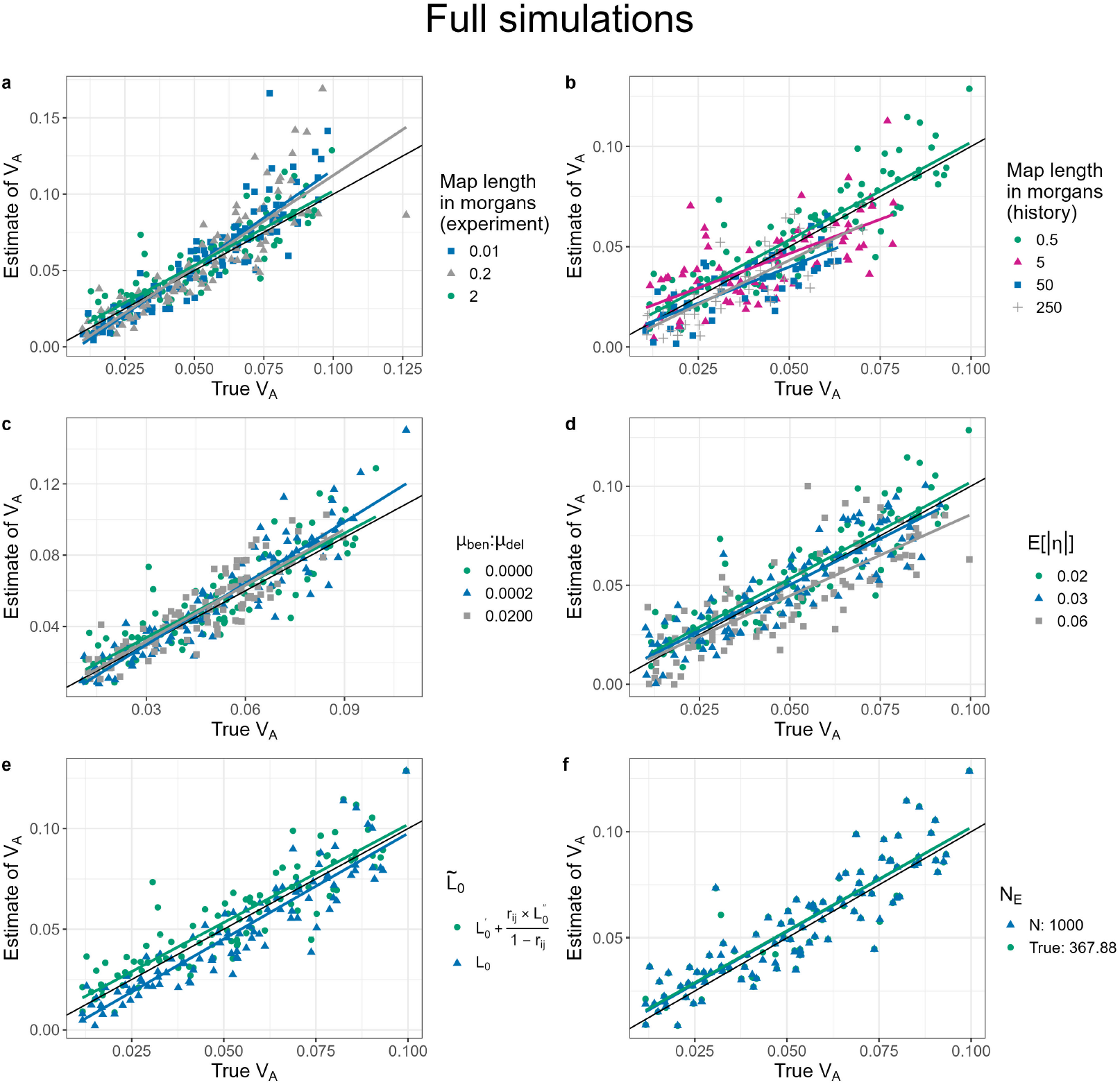
Scatter plots of estimates of *V*_*A*_ vs true values of *V*_*A*_ for full simulations with a burn-in phase of 25,000 generations at different values of (a) map length in the experiment phase, (b) map length during the history phase, (c) the ratio of rates of beneficial and deleterious mutations in the history phase, and (d) the mean of the gamma distribution from which effect sizes for log absolute fitness were sampled for non-neutral mutations (*E*[| *η*|]). In each case, other than the parameter to be varied, the other parameters were fixed at their default values (map length in the history phase = 0.5 morgan, map length in the experiment phase = 2 morgans, number of replicate populations = 10, population size = 1,000, number of generations = 3, *E*[|*η*|] = 0.02, and no dominance (i.e. *κ* = 0)). The solid black line indicates the 1:1 line. The coloured lines represent regression lines for estimates of *V*_*A*_ vs true values of *V*_*A*_. The effect of analysing the standard set of simulations (Figure 4) (e) without partitioning 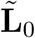 into its gametic 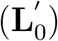 and non-gametic phase 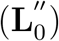 components, and (f) using *N*_*E*_ = *N* = 1, 000. Note that in (F), a large number of green points are eclipsed by the blue points.

### Simulating pool-seq

When we incorporated the expected covariance structure due to pool-seq sampling in our models, the performance of our method was practically unaffected by obtaining allele frequencies in the experiment phase using simulated pool-seq in the simplified simulations (Figure 6A), although a modest negative bias was observed at low (100x) coverage in the full simulations (Supplementary Figure 6). Additionally, both in the simplified and the full simulations, the degree of overdispersion in the number of reads mapping to an individual did not adversely affect our estimates of *V*_*A*_. Along expected lines (see Supplementary information S6), omitting the covariance structure of pool-seq sampling from our models led to estimates of *V*_*A*_ that were significantly downwardly biased, with virtually no *V*_*A*_ detected at 100x coverage (Figure 6B).

**Figure 6:**
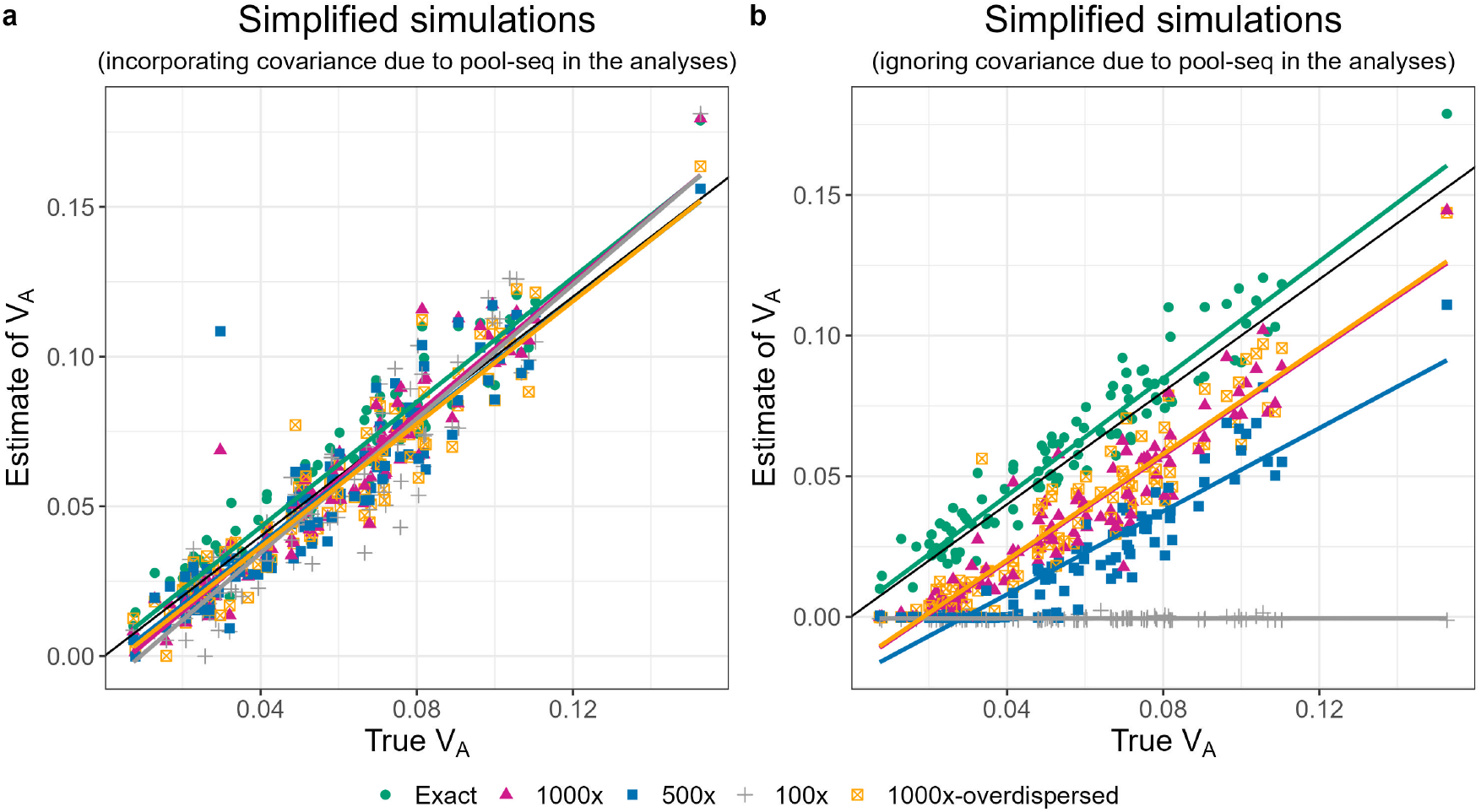
Scatter plots of estimates of *V*_*A*_ vs true values of *V*_*A*_ for simplified simulations (map length in the history phase = 0.5 morgan, map length in the experiment phase = 2 morgans, number of replicate populations = 10, population size = 1,000, number of generations = 3, the mean of the gamma distribution from which effect sizes for log absolute fitness were sampled for non-neutral mutations (*E*[|*η* |]) = 0.02, and no dominance (i.e. *κ* = 0)) using either exact allele frequencies in the experiment phase (green), or allele frequencies in the experiment phase obtained via simulated pool-seq implemented using three different levels of coverage (expected number of reads mapping a segregating site: 1000x (magenta), 500x (blue), and 100x (grey)) without any overdispersion in the number of reads mapping to an individual, as well as estimates obtained from simulated pool-seq implemented at 1000x coverage with overdispersion in the number of reads mapping to an individual (orange). Reads were modelled to be 800 base-pairs long. The solid black line indicates the 1:1 line. The coloured lines represent regression lines for estimates of *V*_*A*_ vs true values of *V*_*A*_. Estimates of *V*_*A*_ were obtained by either incorporating (a) or omitting (b) the expected covariance structure due to pool-seq sampling in the models.

### Comparison with B&C

In both simplified and full simulations, all implementations of B&C’s method resulted in a strong upward bias at low recombination rates in the history phase (Figure 7 and Figure 8). This was likely a consequence of Assumption G being violated; the correlation between the contribution of a selected site *j* to *V*_*a*_ and the persistent association of this site with neutral alleles 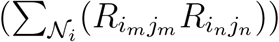 was considerably higher than 0 when the map length in the history phase was low (panel h of Figure 7 and Figure 8). In contrast, the estimates of *V*_*A*_ provided by our method were unbiased and had far greater precision compared to any of the six implementations of B&C’s method in the simplified simulations (Figure 7g), although there was a small upward bias at low recombination rates in the full simulations (Figure 8g). Due to the difficulty of distinguishing selected from unselected sites, B&C suggested that the average LD between all sites could be used instead of the LD between selected and unselected sites (Assumption Ib) and indeed this assumption seems to be well justified (panels d & f in Figure 7 and Figure 8 are very similar). However, an inability to distinguish selected from unselected sites would likely mean that B&C’s method is applied to allele frequency change data from *all* segregating sites, contravening Assumption B. This results in considerable overestimation of *V*_*A*_ in simplified simulations (Figure 7a-b) but not in full simulations (Figure 8a-b). Under all scenarios the bias in the method of B&C can be reduced (but not eliminated) by relaxing Assumption N (compare panels b, d, and f in Figure 7-Figure 8 to panels a, c and e). Under assumption N, it is assumed that the (co)variance in allele frequency change weighted by the square root of twice the genetic diversities can be approximated by the (co)variances in allele frequency change divided through by twice the average diversities. However, this approximation is not required.

**Figure 7:**
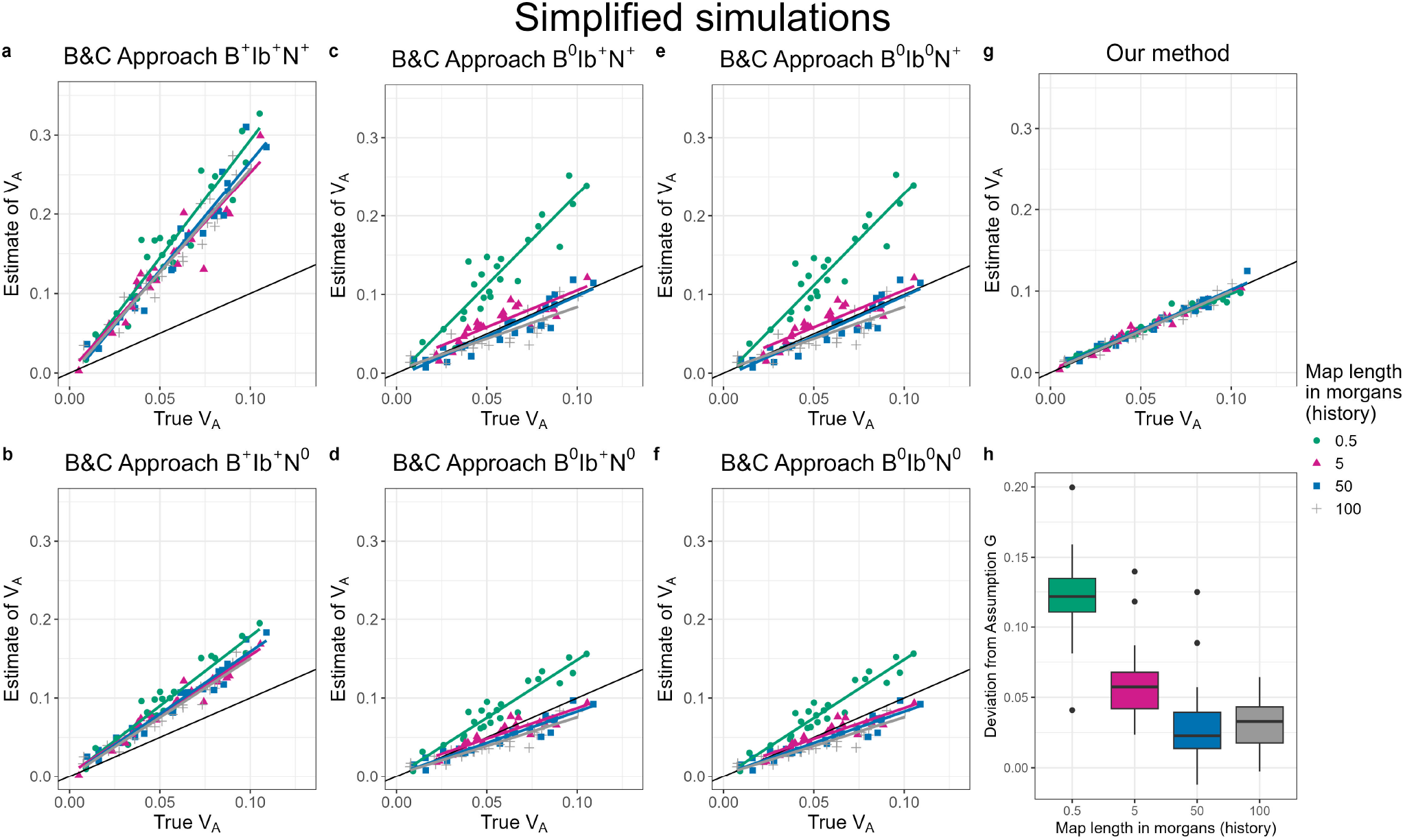
Estimates of *V*_*A*_ obtained using our method (g) and various applications of the B&C Approach (a–f) from simplified simulations with varying map lengths in the history phase: 0.5 morgan (green), 5 morgans (magenta), 50 morgans (blue), and 100 morgans (grey). The solid black line indicates the 1:1 line. Each application of B&C is defined in terms of which three assumptions (B, Ib, N) are required or not (designated with a ‘+’ or ‘0’ superscript, respectively). Assumption B is that allele frequencies are tracked at unselected loci only, Assumption Ib is that the average LD between all sites is equal to the average LD between selected and unselected sites, and Assumption N is that the covariance between projected allele frequency changes is equal to the covariance between allele frequencies changes scaled by the average projection (see Table 2 for details). Panel h is the correlation between the contribution of a selected sites to *V*_*a*_ and their persistent associations with neutral alleles (i.e. deviations from Assumption G in B&C)

**Figure 8:**
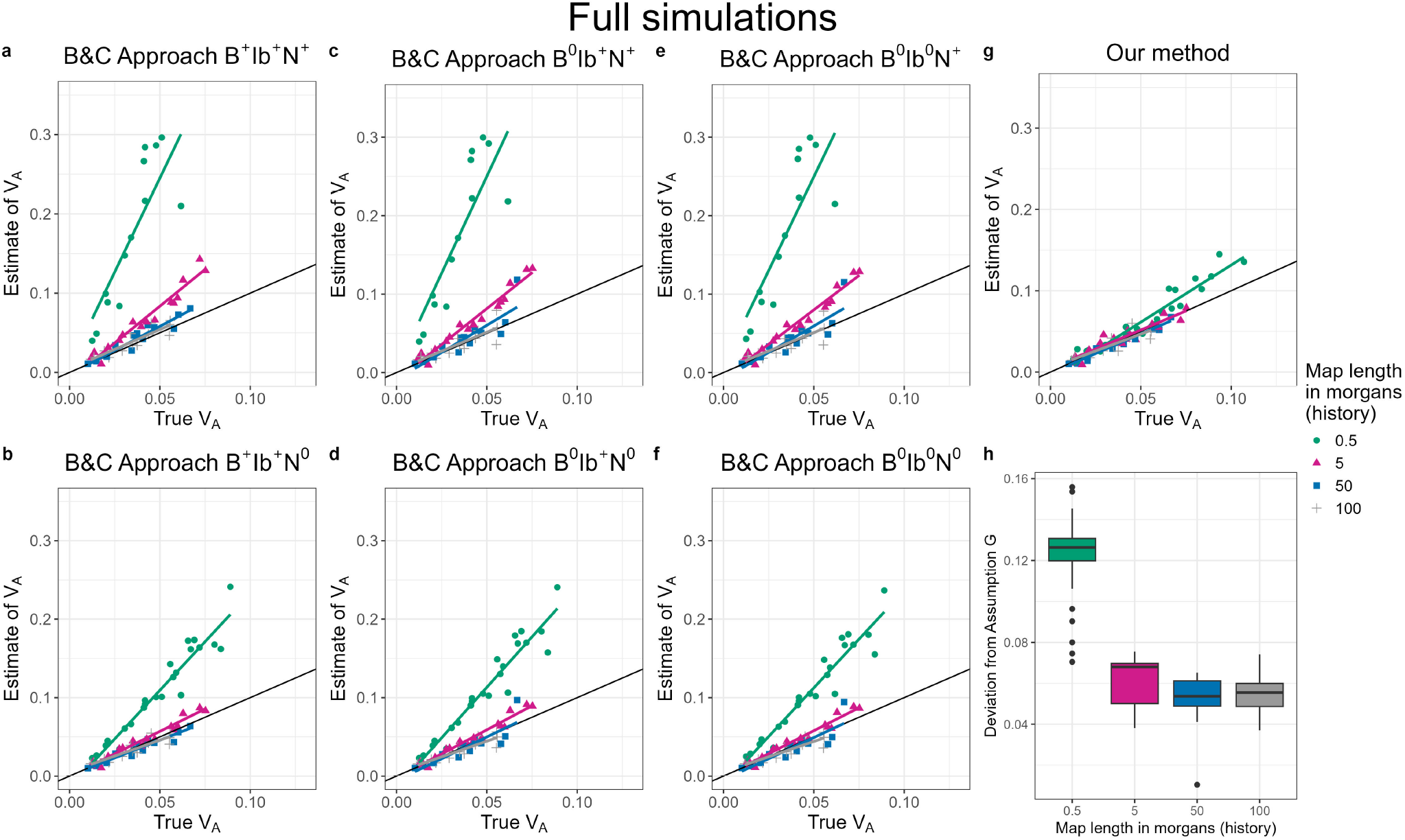
Estimates of *V*_*A*_ obtained using our method (g) and various applications of the B&C Approach (a–f) from full simulations with varying map lengths in the history phase: 0.5 morgan (green), 5 morgans (magenta), 50 morgans (blue), and 100 morgans (grey). The solid black line indicates the 1:1 line. Each application of B&C is defined in terms of which three assumptions (B, Ib, N) are required or not (designated with a ‘+’ or ‘0’ superscript, respectively). Assumption B is that allele frequencies are tracked at unselected loci only, Assumption Ib is that the average LD between all sites is equal to the average LD between selected and unselected sites, and Assumption N is that the covariance between projected allele frequency changes is equal to the covariance between allele frequencies changes scaled by the average projection (see Table 2 for details). Panel h is the correlation between the contribution of a selected sites to *V*_*a*_ and their persistent associations with neutral alleles (i.e. deviations from Assumption G in B&C). For clarity of presentation, in (a), (c), and (e), we have restricted the range of the y axis between 0 and 0.35. As a consequence, points having an estimate of *V*_*A*_ above 0.35 have been excluded from the plot for 0.5 morgan (green circles).

## Discussion

In this paper we estimate the additive genetic variance for relative fitness (*V*_*A*_) directly from the change in the genetic composition of a population caused by selection. Assuming only the absence of meiotic drive and ignoring mutation, we show that *V*_*A*_ can be conveniently expressed as a function of the genome-wide genetic diversity matrix **L**, and the vector of genome-wide expected allele frequency change due to selection, *E*[Δ**p**]. In our inference approach, we describe how a linear mixed model can be employed to estimate *E*[Δ**p**] via independent evolutionary replicates derived from the same base population with a known **L** – a common feature of evolve-and-resequence studies. Unlike alternative methods (Buffalo and Coop, 2019), this allows us to obtain estimates of *V*_*A*_ that are largely robust to the underlying genetic properties of the population and the genetic architecture of fitness. Moreover, the underlying modelling framework not only allows *V*_*A*_ to be estimated, but allows inferences about the relationship between effect sizes and allele frequency.

Although *E*[Δ*p*]’s at individual loci cannot be usefully estimated, for our purposes it is sufficient to estimate their distribution as parameterised through the mean vector 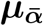 and the (co)variance matrix 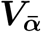 for the distribution of the average effects for fitness, the *α*’s. Our inference approach uses a low-dimensional (3-parameter), but biologically sensible, model for the means and (co)variances. It is superficially surprising that such a simple model for an *n*_*L*_ (the number of loci) dimensional distribution can produce accurate results. However, in a typical dataset one expects the number of individuals, *N*, to be far smaller than the number of loci, *n*_*L*_. Therefore, the non-null subspace of **L** has only *N* dimensions, and we can work with allele frequency changes projected into this reduced space. This provides a route to understanding how the distribution of *α*’s can be estimated from selected changes in allele frequency that must be negligible compared to the impact of drift: the projection defines ‘*chunks*’ of genome, and it is the frequency changes in these chunks, rather than individual alleles, that are tracked. Since the aggregate fitness effects of alleles across chunks will be more substantial, they can be more easily detected. Moreover, since these aggregate effects may involve a large number of loci, they will, thanks to the central limit theorem, tend to normality and, conditional on the projection, converge in distribution. Since the projection is defined by **L**, a model of the *α*’s that conditions on aspects of **L** (*p − q* and *pq* in our case) is expected to be sufficient. The results of our simulations demonstrate that our approach provides usefully precise and consistent estimates of *V*_*A*_ over a wide range of parameter combinations and experimental designs. In the simplified simulations, in which the relationship between allele frequencies and *α*’s was expected to be absent, the model requires only a single parameter: the variance in average effects, 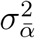. If such a model is assumed, this is a relatively straightforward inference problem and the method of Buffalo and Coop (2019) can give accurate estimates if selected and neutral sites can be distinguished, and certain patterns of recombination and linkage-disequilibrium hold. Although our approach does not assume such a model, it can accurately infer *V*_*A*_ and also the lack of relationship between allele frequencies and *α*’s, with the two parameters that determine the relationship between allele frequency and *α* all centered on their null expectations (Figure 1). However, in reality, independence between allele frequency and *α* seems implausible at mutation-selection-drift balance, and the elevated contribution of high-diversity loci to *V*_*A*_ under the simple, but unrealistic, scenario may result in greater power to detect *V*_*A*_.

At mutation-selection-drift balance a negative relationship is expected between *α*’s and allele frequencies (Charlesworth and Charlesworth, 2010). Although it could be argued that the strength of this relationship will be reduced in experimental evolution studies, which generally expose populations to novel environmental stressors (such as a laboratory environment, pathogens (Basu *et al*., 2024), extreme population densities (Joshi and Mueller, 1996) and temperatures (Singh *et al*., 2015; Hsu *et al*., 2024), desiccation (Gibbs *et al*., 1997), malnutrition (Kawecki *et al*., 2021), or toxic substances (Godinho *et al*., 2024; Xiao *et al*., 2019)) it seems unlikely that no relationship would persist. Even under the more realistic scenario of our full simulations, in which the base population had evolved under selection for 25,000 generations, our approach provided reliable estimates of *V*_*A*_ that were remarkably robust to the details of the distribution of fitness effects, such as the relative frequency of beneficial mutations, or properties of the population such as the recombination rate. Furthermore, it did so by correctly inferring a negative relationship between fitness effects of alleles and genetic diversity, and a positive relationship between fitness effects and *p* − *q*, as expected under mutation-selection balance.

An important assumption of our method is that any consistent change in the *α*’s is vanishingly small in the time-frame over which allele frequency changes are recorded. However, when fitness-causing alleles act non-additively, *α*’s depend on allele frequency and so are expected to change as allele frequencies change. In spite of this, in the simplified simulations where allele frequency change was expected to be large, the performance of the method was minimally affected when allowing dominance at all loci. Moreover, given the form *α*_*i*_ *≈ η*_*i*_[1 − *κ*_*i*_(*p*_*i*_ − *q*_*i*_)] (where the product *η*_*i*_*κ*_*i*_ is always positive in our simulations), the method correctly inferred that the two parameters describing the relationship between the *α*’s and allele frequencies would become increasingly negative with the degree of dominance (Figure 3). Surprisingly, when we allowed dominance effects in the history phase and so allele frequencies and effects had come to an equilibrium we found that our method overestimated *V*_*A*_ to varying degrees depending on how recessive the deleterious allele was. However, this was not because of a failure of the infinitesimal approximation in the experimental phase, but because substantial negative LD had built up during the history phase and our model of 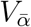 failed to capture this: the additive genetic variance was roughly half the additive genic variance. It seems that our simulations were resulting in substantial pseudo-overdominance (Abu-Awad and Waller, 2023) a phenomenon characteristic of regions of very low recombination (Salson *et al*., 2025). When we performed identical simulations at higher recombination rates in the history phase, this problem was not observed.

Another key assumption of our method is that any consistent, selection-induced changes in **L** are negligible, which then allows us to model the expected evolution of **L** due to the action of drift and recombination. Selection-induced changes in **L** may be safely ignored if the genetic architecture of fitness is sufficiently polygenic such that selection coefficients associated with each locus are vanishingly small. It is important to note that this assumption was fully relaxed in all of our simulations, in which selection was free to influence the evolution of **L** along with drift and recombination. Despite this, the performance of our method was virtually unaffected, even in the simplified simulations where selection-induced changes in **L** were expected to be relatively large.

While our method generally performs well, some biases were observed. In terms of experimental design, small upward biases are evident when power is low either because there are few replicates and/or allele frequency change is only calculated over a single generation. In terms of genetic architecture, our approach marginally underestimates *V*_*A*_ when the fitness effects of new non-neutral mutations are large. This downward bias seems to arise because there is a large contribution of rare highly deleterious variants to *V*_*A*_ and these get lost during the experimental phase. Nevertheless, even when 25% of *V*_*A*_ was lost during the course of the experiment the impact on estimates was rather minor. Similarly, modest downward biases were observed when the recombination rate in the history phase was increased, but the source of this bias has been harder to diagnose. Increasing the recombination rate resulted in an increase in the number of segregating selected sites and their genetic diversity, and a steeper relationship between *α* and genetic diversity, consistent with a reduction in the effects of background selection (Charlesworth *et al*., 1993) on weakly selected sites (Stephan *et al*., 1999). While it is not clear why this causes downward bias in the estimates, it is also unclear whether real populations would ever have such extreme genetic architecture where the bulk of the additive genetic variance for fitness is caused by highly deleterious variants segregating at very low frequencies. In order to make the forward simulations manageable we were working with considerably smaller population and genome sizes than are typical of real populations. It is not clear, however, whether the standard rescaling of mutations rates, recombination rates and selection coefficients to accommodate this downsizing results in genetic architectures that would be typical of larger populations and larger genomes (Dabi and Schrider, 2025).

While our method performs well when applied to simulated data, application to real-world data would involve overcoming a number of challenges. First, we require genome-wide allele frequency change data from multiple independent evolutionary replicates – although this should be readily available for most evolve-and-resequence experiments using approaches such as pool-seq. An important point of consideration, therefore, is the minimum pool-seq coverage that the method requires. For our simplified simulations, there was little deterioration in model performance as coverage dropped or overdispersion increased, suggesting modest coverage (100x) should be sufficient if the appropriate covariance structure is used. While this might be surprising, the relevant parameter is probably not coverage *per se*, but the number of reads overlapping at least one segregating site per unit map length, as this determines how accurately the frequency of a ‘chunk’ of genome can be measured. Using a similar argument, Tilk *et al*. (2019) demonstrated that supplementing pool-seq sampling with haplotype inference tools can result in nearly 500x ‘effective’ coverage at 10x empirical coverage. Given the ancestral haplotype structure is a core requirement of our method, it may be possible that even greater precision could be achieved by also using such tools. However, this could come with the risk that estimation errors are correlated across replicates and be mistaken for patterns of selection. For our full simulations, a moderate downward bias was observed at 100x coverage. While this may suggest that our method demands relatively high coverage (≥ 500x), it is worth noting that we only implemented an approximate (i.e. diagonal) covariance structure for pool-seq sampling while analysing our simulations due to computational limitations. Incorporating the full covariance structure should lead to improved estimates.

Second, we require the genetic diversity matrix **L** in the base population from which the replicates are derived. This is not always the case for evolve-and-resequence studies, in which base populations are often split into replicate baseline populations long before the experiment, or when newer selection regimes are derived midexperiment (Burke *et al*., 2010; Singh *et al*., 2015; Gupta *et al*., 2016; Robinson *et al*., 2023). Furthermore, our method requires that sufficient individuals from the base population are individually sequenced to estimate **L**, or – even better – its gametic phase (**L**^*′*^) and non-gametic phase (**L**^*′′*^) components, although the required phasing should become more readily available with long-read sequencing. Third, to predict how **L** evolves, we require an estimate of the recombination probability between all pairs of segregating sites, such as a recombination map for the population. In reality, recombination maps are likely to have been derived from other populations, in which recombination patterns may differ (Johnston, 2024). Although recombination maps can be approximated, for example by using Haldane’s mapping function, it is not clear how sensitive the method is to errors in the recombination map. Fourth, our method requires the mean number of generations over which allele frequency changes are calculated, which may be hard to infer with overlapping generations. Since allelefrequency change will be roughly proportional to the number of generations that have elapsed (*n*_*g*_), and *V*_*A*_ is quadratic in allele-frequency change, estimates of *V*_*A*_ might be out by a factor 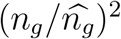, where 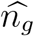 is the assumed number of generations. Fifth, our method uses effective population size to predict how **L** evolves (*N*_*e*_) and to derive expressions for the drift (co)variance (*N*_*E*_). Although reliable estimates of both effective population sizes may be hard to obtain, this is unlikely to be a major issue because: (i) **L**^*′*^ decays with a rate roughly proportional to 1 − 1*/N*_*e*_ (likely making it insensitive to errors in estimating *N*_*e*_) and (ii) our simulations suggest that using the wrong *N*_*E*_ does not adversely affect our estimates of *V*_*A*_ because the misspecification is absorbed by the residual variance of the model. Sixth, our simulations minimise the unpredictable response to selection by modelling a constant environment. However, with selection coefficients that vary in time or across replicates, the unpredictable response to selection will be greater. Indeed in outdoor mesocosms of *D. melanogaster*, Bitter *et al*. (2024) report that allele frequency changes can switch signs over time-points separated by a matter of weeks in spite of exhibiting highly concordant evolution between replicates. It is not clear to what degree this will affect inferences. Finally, selection can often act in different ways in different contexts such as space (Whitlock, 2015; Delph, 2018), time, and between the two sexes (Schenkel *et al*., 2018). Our approach captures the effects of selection averaged over all these different contexts. Specifically, if loci have different fitness effects in males and females, we effectively estimate (*V*_*A,f*_ + *V*_*A,m*_ + 2*COV*_*A,mf*_)*/*4, where *V*_*A,f*_ and *V*_*A,m*_ are the additive genetic variances for relative fitness in females and males respectively, and *COV*_*A,mf*_ is the intersexual additive genetic covariance for relative fitness. For these reasons, when applied to replicate populations, our estimates of *V*_*A*_ are perhaps best thought of as the additive genetic *covariance* in fitness between replicates. Although rarely made explicit, estimates from wild systems should be interpreted in the same way: the additive genetic covariance between the environments in which relatives live (Vehviläinen *et al*., 2008).

Analysing data from wild populations is likely to entail further challenges. For example, in natural populations, allele frequency change from immigration may be consequential, and without some modification is likely to be mistaken for allele frequency change caused by natural selection (Simon and Coop, 2024). Furthermore, replicate populations are unlikely to be available for natural systems – with some exceptions such as Trinidadian guppies (Reznick and Bryga, 1996) – although with some modifications our method could be adapted to use allele frequency change data from multiple time points. However, a lack of individual-level sequences in the base population would mean that the estimates of **L** will likely be considerably noisier in natural populations.

While we accept that the data requirements for our method are steep, and the list of caveats appears long, we do think that information about *V*_*A*_ can be successfully leveraged from current evolve and resequence studies. Going forward, we hope this work will inform future evolve and resequence study design, and that estimates of *V*_*A*_ from a wide range of organisms and environments become available. Partitioning *V*_*A*_ into genomic features is an obvious next step, and in the future we hope to go beyond simply knowing the magnitude of *V*_*A*_ and start to understand its underlying causes.

## Data availability statement

This study does not use any data. The code used for simulations and analyses is available in the following GitHub repository: https://github.com/manas-ga/Va_simulations.

## Acknowledgments

The authors would like to thank Bill Hill, Brian Charlesworth, Peter Keightley, Konrad Lohse, Bruce Walsh, Ben Longdon and Vince Buffalo for their insightful comments, and The Argyle for hosting us at the early stages of this project. Computer simulations and analyses described here were performed on the AC3 computing cluster based at Ashworth Laboratories, and on Eddie, the University of Edinburgh’s main high performance computing facility.

## Supplementary information

## S1 The dynamics of L under drift and recombination

As outlined in the section ‘Extending our approach to practical situations’ our goal is to infer the additive genetic variance for relative fitness in the base population (*V*_*Ā*_(0)) using genome-wide allele frequency changes between generations *t* and *τ* . In order to derive the expected allele frequency change due to selection over this time period, we first derive the expected dynamics of **L** under drift and recombination.

The matrix **L** is a covariance matrix whose elements are proportional to the genotypic linkage-disequilbria (off-diagonals) or variance in genotypic allele frequencies (diagonals) at the start of a generation, before selection has acted. We can decompose **L** into **L**^*′*^ and **L**^*′′*^ following the notation of Buffalo and Coop (2019) where **L**^*′*^ represents the (co)variances that arise due to alleles in the same gamete and **L**^*′′*^ represents the (co)variances that arise due to alleles in the different gametic contributions of a genotype. Note that in Santiago and Caballero (1998) the primes have a subtly different meaning after the initial generation, as the double prime in following generations designates gametic phase disequilibrium due to recombination and nongametic phase disequilibrium in the initial generation. The elements of **L**^*′*^ and **L**^*′′*^ have direct correspondences with genetic diversities and additive measures of disequilibria. Under the notation of Weir and Cockerham (1989) (see also Bulmer (1980), Chapter 12), the diagonal elements of **L**^*′*^ are half the gametic genetic diversities (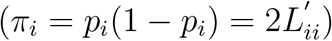, and the off-diagonals are half the gametic-phase disequilibria 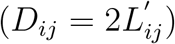. Note that Weir and Cockerham (1989) uses *π* to denote *p*_*i*_(1 − *p*_*i*_) rather than the more usual (and less natural) 2*p*_*i*_(1 − *p*_*i*_). The diagonal elements of **L**^*′′*^ are half the additive coefficients of Hardy Weinberg disequilibria 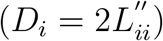, and the off-diagonals are half the nongametic-phase disequilibria 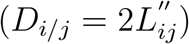.

To see these correspondences, imagine two bi-allelic loci, *i* and *j*, with reference/alternate alleles A/a and B/b, respectively. There are four possible gametic haplotypes, and we can denote the frequency of haplotype *AB* as *p*_*AB*_ in the gametes and the frequency of allele *A* as *p*_*A*_. The proportion of copies of the reference allele at locus *i* in a randomly chosen individual, *c*_*i*_, can be decomposed into into the sum of maternal and paternal contribution: *c*_*i*_ = *m*_*i*_ + *f*_*i*_ where *m*_*i*_ (or *f*_*i*_) takes the value 1/2 if the mother (or the father) contributed a reference allele and 0 if not. Then,

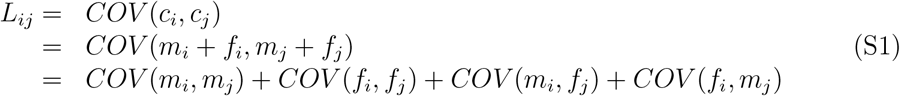

Assuming haplotype frequencies are identical in male and female gametes we get

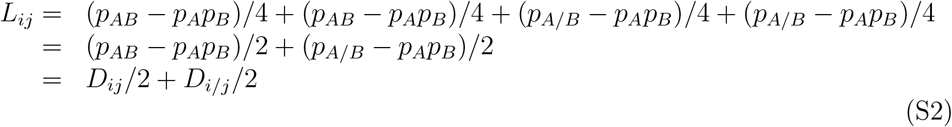

where *p*_*A/B*_ is the frequency of zygotes that have an A from their mother and a B from their father, or vice versa. When *i* = *j*,

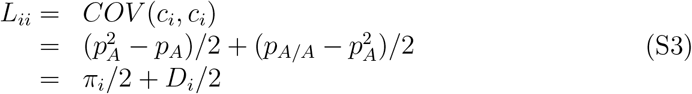

where *p*_*A/A*_ is the frequency of *A* homozygotes. The term *D*_*i*_ is an alternative, additive, measure of deviation from Hardy-Weinberg Equilibrium than the more commonly used inbreeding coefficient, *F* .

To derive expressions for the dynamics of **L**^*′*^ and **L**^*′′*^, note that **L**^*′′*^ is generated anew each generation and under random mating has zero expectation such that

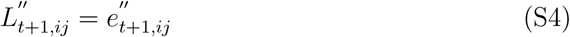

where 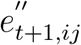 is a stochastic term with mean zero and variance given in Weir (1996). The elements of **L**^*′*^ under recombination and drift are (Hill and Robertson, 1968; Santiago and Caballero, 1998)

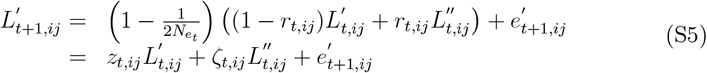

where 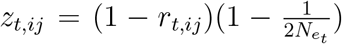 and 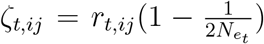, with *r*_*t,ij*_ being the recombination rate between locus *i* and *j* in generation *t*, and 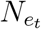 the effective population size in generation *t*. In the absence of selection, the stochastic terms, 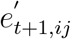, have zero mean and are uncorrelated over time, with variances given by Ohta and Kimura (1969, Eq 25, although those variances must be divided by 4 here since we are working with the frequency of alleles in individuals rather than gametes). The covariances between, for example, 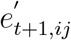 and 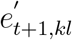 are not given in Ohta and Kimura (1969), and may be unknown. Using the above results we can derive a recursion for *L*^*′*^ at some arbitrary time *t*:

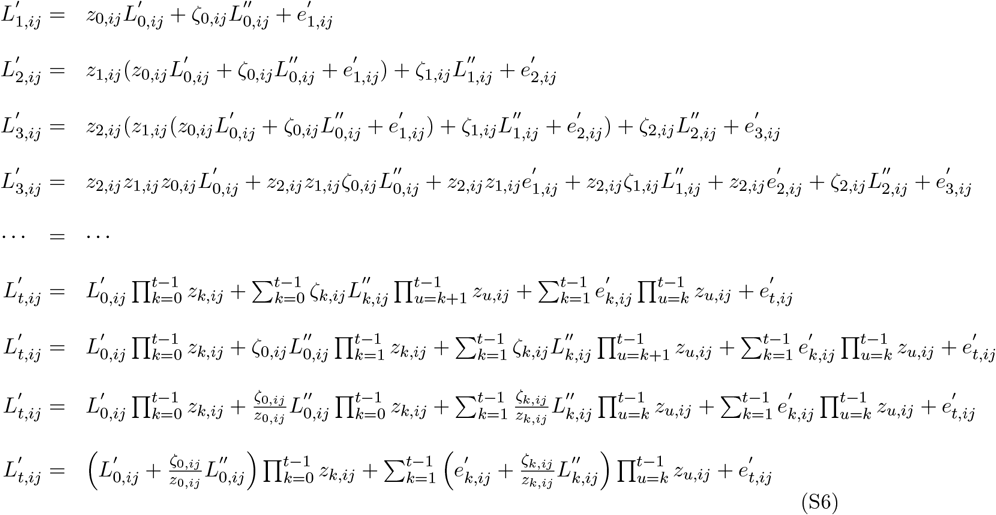

Having the matrix **N**_*t*_ with the *ij*^*th*^ element 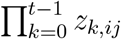, we have for *t >* 0

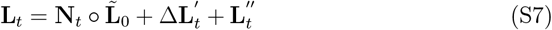

where is the Hadamard (i.e. lement-wise) product. The first term is the expected **L** (and **L**) in generation *t*, conditional on the genotypic composition of the population in generation 0, where 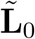 is the sum of 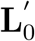 and 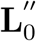 with the elements of the latter weighted by *ζ*_0,*ij*_*/z*_0,*ij*_ = *r*_0,*ij*_*/*(1 − *r*_0,*ij*_). Δ**L**^*′*^ is a matrix with elements equal to the sum of the stochastic terms involving *e* in Equation S6 and represents stochastic changes in **L**^*′*^ from generation 0 to generation *t*. 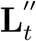 are the new nongametic-phase disequilibria that arise in generation *t*. Note that if replicate populations are initiated from the offspring of Generation 0, then the stochastic terms, 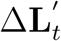 and 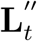, will be unique in each replicate although the deterministic part is shared. If recombination rates are constant in time *r*_*t,ij*_ = *r*_*ij*_ then *ζ*_0,*ij*_*/z*_0,*ij*_ = *r*_*ij*_*/*(1 − *r*_*ij*_) and 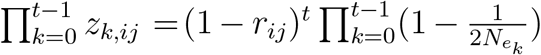. When population sizes are also constant then 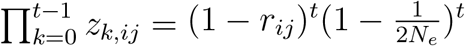.

## S2 The mean and (co)variance of allele frequency changes

Given our model for the dynamics of **L** (See Supplementary information S1) we can derive the mean and (co)variance of the allele-frequency changes within a replicate. Here expectations and variances are taken with respect to the evolutionary process, which not only includes the effects of drift 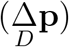 but also stochastic changes in **L** and ***α***. The change in allele frequency in replicate *m* is

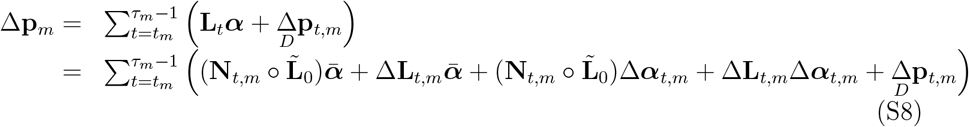

where 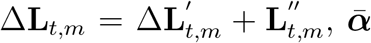 is a vector of mean average effects and Δ***α***_*t,m*_ a vector of deviations specific to time *t* and replicate *m*. By definition, drift is nondirectional and so 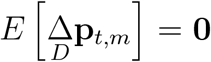. We also assume that the unpredictable response to selection has zero expectation which requires that deviations in **L** and ***α*** from their predicted means are zero on average (*E*[Δ**L**] = **0** and *E*[Δ***α***] = **0**) and there is no correlation between them (*COV* (Δ**L**, Δ***α***) = 0). If the genetic architecture of fitness is sufficiently polygenic, selection-induced changes in **L** may be negligible. Similarly, since the expected change in **L** is slow, the induced change in ***α*** in the presence of non-additive gene action should be minor. Making these assumptions we have:

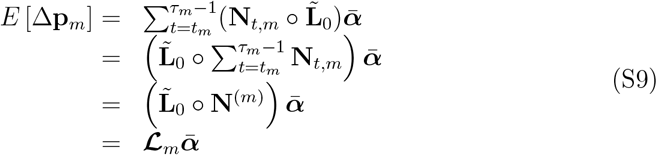

where 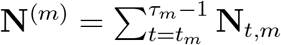 and 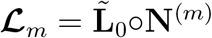. The within-replicate (co)variances are more challenging to derive. The variance due to the unpredictable selection at time *t* in replicate *m* is

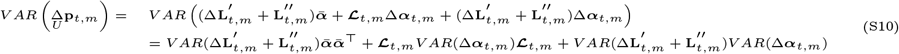

In addition, covariances between the unpredictable response at different time points within a replicate will be non-zero because 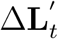 is the sum of stochastic changes from generation 1 to *t* and so 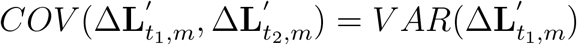 when *t*_1_ ≤ *t*_2_, although 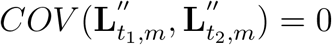. This gives

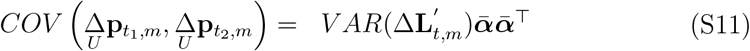

when *t*_1_ ≤ *t*_2_. Consequently, the total variance in the unpredictable response to selection will be

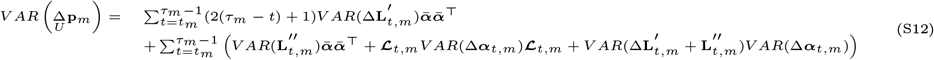

and cannot be simplified we simply refer to it as **𝒰** _*m*_.

For the drift (co)variances, note that under random mating, haplotypes are drawn from a multinomial with 2*N* trials. Using AB, Ab, AB and aB to denote the *number* of each haplotypes in the gamete pool, the number of *mn* haplotypes has variance 2*Np*_*mn*_(1 − *p*_*mn*_) and the covariance in the numbers of *mn* and *op* haplotypes is −2*Np*_*mn*_*p*_*op*_ where *p*_*mn*_ is the frequency of the *mn* halpotype in the gamete pool (i.e the parental haplotype frequencies modified by recombination). The covariance in the number of A and B alleles sampled, is therefore

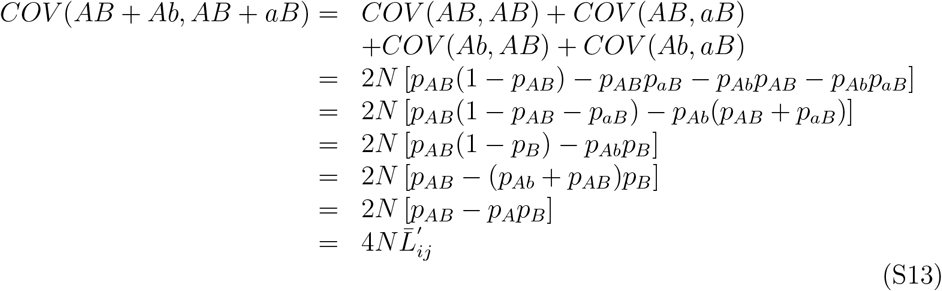

where 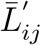 is the gametic-phase linkage disequilibria that would be achieved in an infinite population. We can divide through by (1*/*2*N*)^2^ to obtain the drift covariance in frequency (rather than counts) as 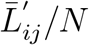 (i.e the drift (co)variances in a allele fre-quency from generation *t* to *t* + 1 are proportional to the gametic-phase disequilibria in generation *t* + 1 that would be achieved in an infinite population conditional on the genotypic composition of the population in generation *t*). This recovers the well known result for the drift variance in allele frequency: 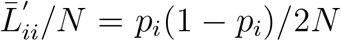. We can replace the census population size, *N*, with the effective population size *N*_*E*_, since this approximates the sampling of genotypes in non-idealised populations well (Ethier and Nagylaki, 1980). However, note that *N*_*E*_ differs from *N*_*e*_ in that it does include the impact of linked selection since this is conditioned on in the expectation, *E*[Δ**p**]. In matrix terms (since 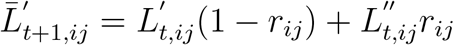):

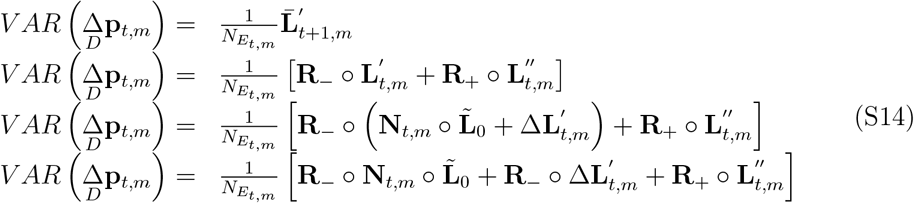

where **R**_+_ and **R**_ are matrices with the *ij*^*th*^ elements being *r*_*ij*_ and 1 − *r*_*ij*_ respectively. The expected drift terms (averaged over the evolutionary stochasticity in **L**) is therefore:

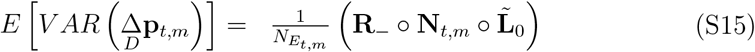

We define a new matrix **M**_*t,m*_ with the *ij*^*th*^ element being 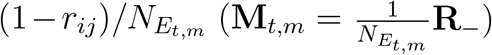 to give

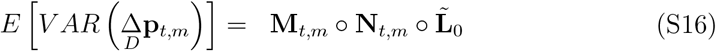

Since the drift terms are independent, this gives:

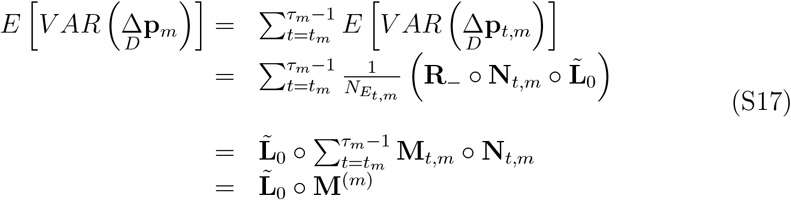

where 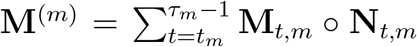 and is null when *t*_m_ = 0 and *τ* = 1. Note the *ij*^*th*^ element of 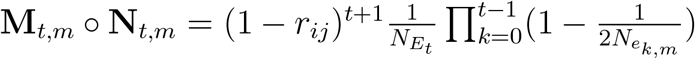 which reduces to 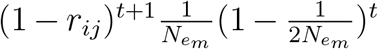 with constant population size and 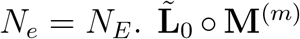 is denoted as **𝒟**_*m*_ in the main text. It is not clear how large the evolutionary variance in 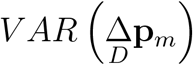 is, and whether this would need to be accommodated.

When the vector of average effects, ***α***, is constant in time and across replicates, 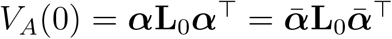. However, when they do vary, this equality does not hold and so we use the notation *V*_*Ā*_(0) to distinguish 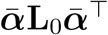 from *V*_*A*_(0). We claim that *V*_*Ā*_(0) can be interpreted as the genetic covariance in fitness between time-replicate combinations for a population with a genotypic composition identical to the base population: it is the expected covariance in breeding value of an individual taken from the base population and (hypothetically) raised in two different replicates at two different times.

To see this, we switch from our original definition of ***α***_*t,m*_ as being equal to 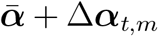 and with some abuse of notation use the form 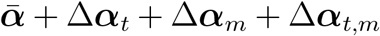 where Δ***α***_*t*_ is the deviation of the average effects (averaged over replicates) at time *t* and Δ***α***_*m*_ is the deviation of the average effects (averaged over time) in replicate *m*, leaving Δ***α***_*t,m*_ as the ‘residual’ deviation in ***α***. By definition, the terms Δ***α***_*t*_, Δ***α***_*m*_ and Δ***α***_*t,m*_ have expectation zero, where the expectations are taken over time, replicates and all time-replicate combinations respectively. 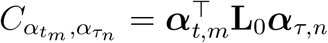 is the additive genetic covariance between replicate *m* at time *t* and replicate *n* at time *τ* given the base population structure.

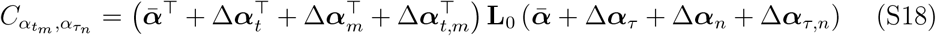

If we take the expectation of this over pairs of replicates we get:

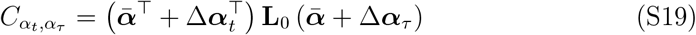

since the Δ terms, and the product of pairs of Δ terms when *n* ≠ *m*, have zero expectation. Here, 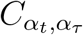 is the between-replicate covariance at time *t* and *τ*, and has expectation (with respect to time) 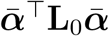 when *t* ≠ *τ* and 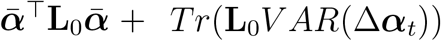 when *t* = *τ* where *V AR*(Δ***α***_*t*_) are the (co)variances of ***α*** over time.

## S3 Treating *α* as random

In standard quantitative genetic theory, ***α*** is considered fixed and the genotypes of individuals in the population are considered random, leading to a distribution of breeding values with variance *V*_*A*_. More recently, however, dense molecular markers have been used as covariates for predicting genetic values, and the associated coefficients have been treated as random (Meuwissen *et al*., 2001). As noted by Gianola *et al*. (2009), this raises interpretational problems when trying to relate the variance parameters associated with the coefficients to the concept of *V*_*A*_.

In our inference section we show that

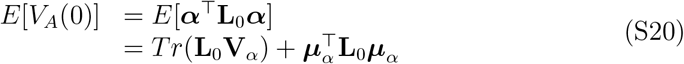

where here we assume the average effects are constant across time/replicates for ease and ***µ***_*α*_ and **V**_*α*_ are the mean vector and covariance matrix for the average effects (see the Appendix in Gianola *et al*. (2009) also). Critically, we interpret this expectation as an average over the epistemic uncertainty in ***α***: it is a posterior mean after marginalising the distribution of ***α*** (but conditioning on ***µ***_*α*_ and **V**_*α*_):

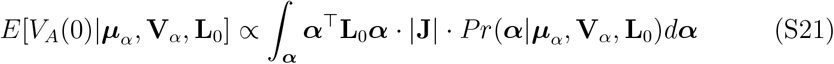

where **J** is the Jacobian of the transform from ***α*** to *V*_*A*_ and depends on **L**_0_.

For an inferential method we see no contradiction between treating ***α*** as fixed when *defining V*_*A*_ but treating it as random when *estimating V*_*A*_. However, when treating 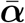 as random it is important to ensure that the underlying model is invariant with respect to which allele is chosen as the reference. Quantities such as *V*_*A*_ and genomic best linear unbiased predictors (gBLUP) are insensitive to which allele at a locus is chosen as the reference allele and which allele is chosen as the alternate allele. To make this explicit, consider the diagonal ‘assignment’ matrix **A** for which the diagonal elements are either 1 (the fittest allele is the reference allele) or -1 (the fittest allele is the alternate allele). Under a particular assignment, ***α*** = **A*α***_+_ and **L** = **AL**_+_**A**, where the subscript + indicates the quantity had the fitter of the two alleles been the reference allele at all loci. If we consider *V*_*A*_ conditional on a particular assignment we get (since **AA** = **I**):

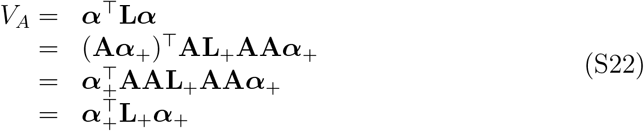

showing we get the same value of *V*_*A*_ irrespective of the assignment we choose. In contrast, quantities such as *E*[**L*α***] are sensitive to the assignment, since they undergo a sign reversal under a different choice:

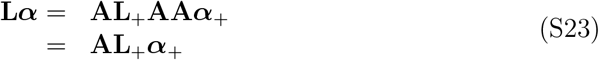

It is tempting to use the argument that *E*[**L*α***] = **0** when the reference allele is chosen at random, since *α* is equally likely to be positive as negative. The logic behind this argument can be expressed mathematically as *E*[***α***] = *E*[**A**]*E*[***α***_+_] since the reference allele is chosen at random and so **A** must be independent of ***α***_+_. Under this same assumption *E*[**A**] = **0**, since any diagonal element has an equal chance of being -1 or 1, such that *E*[**L*α***] = 0. However, this logic is incorrect. The argument envisages **A** as random, yet for any particular analysis **A** is no longer a random variable but fixed - a choice has been made as to which allele is the reference allele - even if there remains epistemic uncertainty as to whether the reference allele is the fitter of the two alleles.

When the reference allele is chosen arbitrarily, any sensible distribution for ***α*** must induce the same distribution on ***α***_+_ regardless of the assignment. If 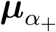 and 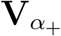 are the means and (co)variances of the average effects had all reference alleles been the fitter allele, then the distribution for a particular assignment becomes 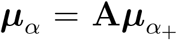 and 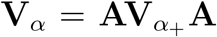. Given **A**^−^ = **A** this implies 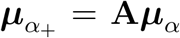 and 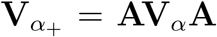. For, ***µ***_*α*_ this implies that suitable models should be (weighted) sums of differences between invariant properties of the alleles such that the difference reverses sign when the reference and alternate allele are switched. This might be their (log) frequency, such that the model is *β*(*p* − *q*), with *β* a parameter, or it might be the difference in derived vs ancestral coded as 1 vs -1, such that the model is 2*β* or −2*β* depending on whether the reference allele is derived or ancestral, respectively.

If 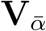 is assumed to be diagonal, all models are permissible since the square removes any sign. Since the multiplication of diagonal matrices is not affected by order we can see this directly:

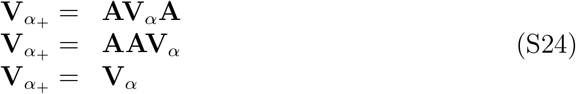

However, for non-diagonal matrices a suitable distribution must result in a sign reversal of all covariances at a locus when the reference and alternate alleles are switched. The most obvious way to achieve this is to allow **V**_*α*_ to be proportional to **L**^*p*^ since

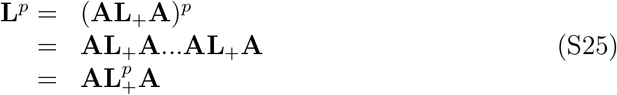

where **L**_+_ is the linkage-disequilibrium matrix had the fittest allele been the reference allele at all loci. Under this assumption

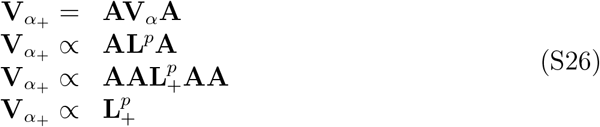

A similar model was proposed by Zeng *et al*. (2018), although there, **L** was treated as diagonal such that the variance of the average effects are assumed to be proportional to the genetic diversities to some power.

## S4 Projection matrices

Rather than working with original allele frequencies, we project them into a new reduced subspace defined by **L**_0_ and claim that *V*_*A*_(0) does not depend on allele frequency changes outside of this space. de Los Campos *et al*. (2015) (in Supplementary Methods I) show that additive genetic variances are rotationally invariant: they are invariant to arbitrary linear transformations of the genotypes. Have **U** be *all* eigenvectors of **L**_0_ and **D** a diagonal matrix with the eigenvalues of **L**_0_ square-rooted along the diagonal. If we use the transformation **U**^⊤^, de Los Campos *et al*. (2015) show that the average effects in the projected space are **U**^−⊤^***α*** and the covariance of the projected genotypes is **U**^⊤^**L**_0_**U**, and:

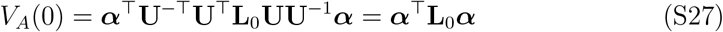

Since **L**_0_ = **UDDU**^⊤^,

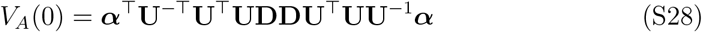

which reduces to

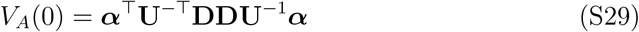

since eigenvectors are unitary **U**^−1^ = **U**^⊤^ and so **UU**^⊤^ = **I**. The maximum rank of **L**_0_ will generally be the number of individuals in the base population and will be substantially less then the number of loci. Consequently some diagonal elements of **D** will be zero which will set all elements in the corresponding column of **U**^−⊤^**D** to zero. Consequently we can simply define the projection matrix **U**_**L**_ with all columns associated with zero eigenvalues dropped without loss of information.

To show that our chosen projection 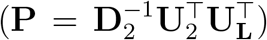 results in identical and independently distributed residuals, we need to show that the drift covariance matrix is an identity matrix under this projection. If we write down the eigendecomposition of 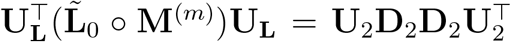, and use the unitary property of **U**_**L**_ to note that 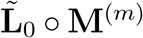 can be expressed as 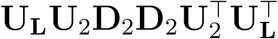. The drift covariance matrix under the projection is then:

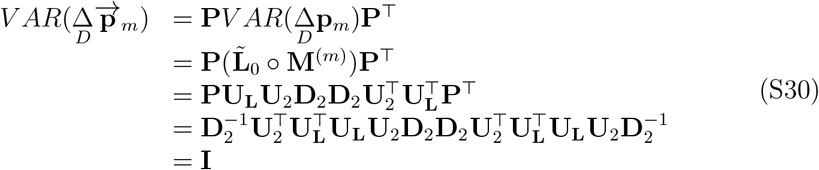

## S5 Bias correction for 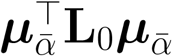

The expression for *E*[*V*_*Ā*_(0)] involves a quadratic in the vector of expected average effects, 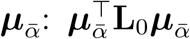. Replacing 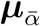 with an estimate of it, 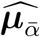, will generally result in the estimate of *V*_*Ā*_(0) being upwardly biased even if 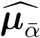 is an unbiased estimate of 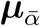. To understand this, and correct for it, have the linear model

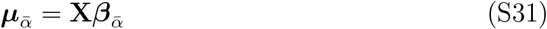

where **X** is the fixed-effect design matrix and 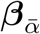 an associated parameter vector.

In our analyses of simulated data, **X** only has one column, **p**_0_ − **q**_0_, and 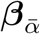 scalar: 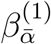. We can have

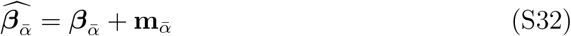

where 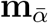 is the vector of deviations of the estimates from their true value. The expected estimate of 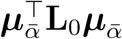 is then

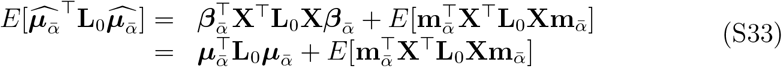

Here, the expectation is taken over the distribution of estimates, and it is assumed the estimates are unbiased (since then, 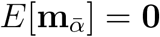, and so 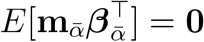). This same assumption implies

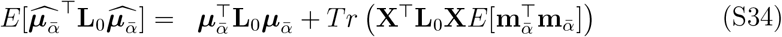

Since 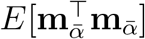 is a matrix of sampling (co)variances for the parameters, to get an improved estimate we can use the inverse Hessian to get an approximate 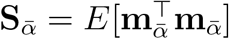:

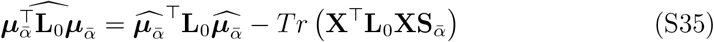

In addition, 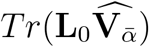 is most likely an upwardly biased estimator of 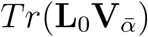. However, we found no easy way to determine or correct for the degree of bias, and the simulation results suggest that the bias is likely to be small, at least when the number of replicates or generations is moderately large.

## S6 Analysing allele frequency data from pool-seq

The main derivation of our method assumes allele frequency changes in each replicate are known without error, and the majority of our simulations were analysed with this being true. However, this assumption is unlikely to be met in reality and here we develop a method for incorporating uncertainty in allele frequencies estimated using pool-seq.

Assuming that all individuals have the same probability of being sampled, the covariance in reference allele number at locus *i* and *j* is equal to the number of reads that span both sites, *O*_*ij*_, multiplied by the covariance in allele number across haplotypes 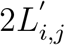 . To get the covariance in reference allele frequency we must divide this by the number of reads spanning site *i* (*O*_*ii*_) and the number of reads spanning site *j* to get 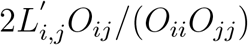. When *i* = *j* this reduces to the familiar binomial variance for allele frequency at a single site 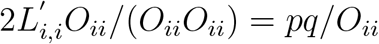. When analysing binomial data the variance often exceeds its expectation - a phenomenon known as overdisperion. In the context of pool-seq this will be mostly driven by individuals, rather than gametic contributions, varying in their probability of being sampled. In this instance, the exact form of the sampling (co)variances will depend in a complicated way on both **L**^*′*^ and **L**^*′′*^, but here we assume that for sites close enough to be spanned by the same read **L**^*′*^ dominates over **L**^*′′*^ and at a site *pq* dominates over **L**^*′′*^ . We then simply accommodate any overdispersion using a parameter that scales the sampling (co)variances, 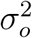, which is constant over loci. This approach is widely used when analysing binomial data (the qausibinomial approach - (McCullagh and Nelder, 1989)) and values greater than one indicate overdispersion. In what follows we also assume 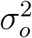 is constant across replicate/time-points although this could be relaxed.

Having the matrix **Q** with the *ij*^*th*^ element equal to *O*_*ij*_*/*(*O*_*ii*_*O*_*jj*_) the sampling covariance in allele frequency in replicate *m* at time *t*_*m*_ is equal to 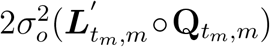. Since errors in allele frequency estimates are independent at different time points the expected sampling covariance in allele frequency change is proportional to:

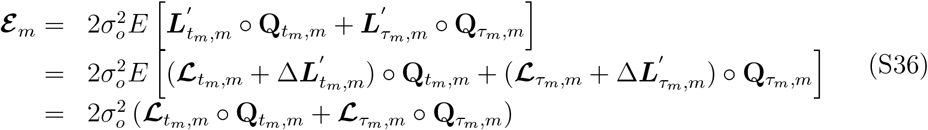

where the expectation is taken over evolutionary realisations of **L**^*′*^ and depends on 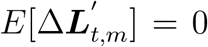. The expected sampling covariance in projected allele frequency change is therefore:

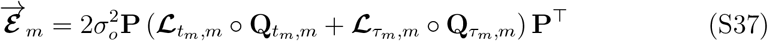

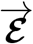 is a square dense matrix equal in dimension to the number of projected loci. Moreover, it will vary over replicates such that the complete covariance structure across all replicates is a large block-diagonal matrix with large blocks. Given the computational burden this entails we fitted approximate models to the simulated data where 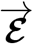 was diagonalised. For the same reason, we also used a diagonalised **Q** matrix which only contains information on coverage rather than complete information on the number of reads overlapping pairs of sites. In addition we also assumed 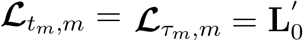 for all *m*. For real data we advocate not using these approximations.

Estimation error in allele frequency at a single-site is comparable to drift, with coverage *O*_*ii*_, or effective coverage 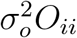, equivalent to *N*_*E*_. Because of this it might be tempting to think that estimates of *V*_*Ā*_(0) would remain unbiased if estimation error was ignored since the residual variance will soak up the excess variance when model fitting. The only advantage, then, of including 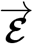 is to increase precision when there is heterogeneity in coverage across sites. However, this is not the case - estimates of *V*_*Ā*_(0) will be downwardly biased if estimation error is ignored and this bias can be substantial unless reads are very long and/or coverage is high. The reason for this is that a major source of information on *V*_*Ā*_(0) comes from correlated changes in allele frequency at loci that are in LD, and in particular correlations that are tighter than expected under drift. If reads only span single sites then the error covariance in allele frequency is zero since *O*_*ij*_ = 0 resulting in correlations that are weaker than under drift which partly destroys the signal of selection. If reads are longer, then the error covariance in allele frequency will come to resemble that caused by drift and any bias when ignoring the errors should be reduced. However, at very long read lengths the bias may even become positive as the sampling correlations between distant sites on the same chromosome will be higher than that under drift where it will be broken down by a round of recombination (Equation S14).

## S7 Comparison with the method of Buffalo and Coop (2019)

To make clearer the distinction between the approach of B&C and our approach, here we explicitly express the expectations and covariances appearing in B&C as conditional on **B** and ***α***. In the sections dealing with our theory and inference, the conditioning (on **L**_0_) is left implicit. B&C work with the quantity

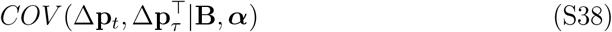

where *τ* is some generation after *t* (B&C actually only work with the diagonal elements of this matrix, but we retain the full multi-locus model here for generality). Importantly, when B&C estimate the additive genetic variance in fitness they assume that both Δ**p**_*t*_ and Δ**p**_*τ*_ both represent allele frequency change over a single generation (Assumption A). As the change due to drift will be independent in different generations, we can rewrite B&C’s covariance:

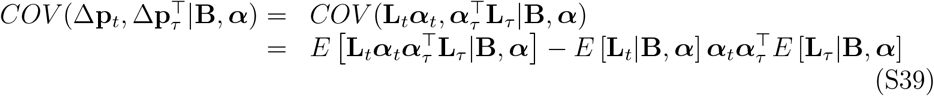

Since the diagonal elements of **L** have to be positive, a sufficient, but not necessary, condition for the final term to be zero is that there is no direct selection on the loci (i.e. ***α*** = **0**). It is not a necessary condition because the change caused by direct selection at all loci could be exactly balanced by the change caused by indirect selection at other loci, although we ignore this unlikely scenario. The assumption that ***α*** = **0** is achieved in B&C by assuming that sites can be partitioned into neutral and selected sites and that allele frequency change is only tracked at the neutral sites (Assumption B). To understand the consequences of this assumption we consider all loci are being followed, both a selected set (𝒮) and a neutral set (𝒩). Consequently,

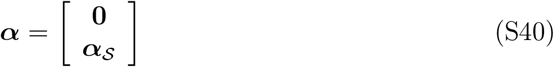

and so

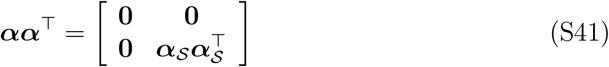

We can also partition **L**

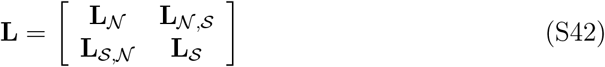

and writing **L**_*N*,*S*_ = **B**_*N*_ **R**_*N*,*S*_ **B**_*S*_, Equation S39 for neutral sites becomes

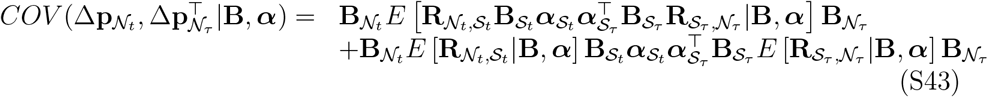

Under the conditioning of B&C, 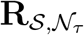 is a random variable. If we assume that the linkage-disequilibrium between the neutral alleles and the selected alleles has arbitrary sign, then 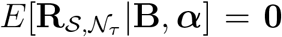, which will be met if the reference allele is chosen arbitrarily (e.g. not based on minor allele frequency (Good, 2022)). Under this assumption (Assumption C) the final term in Equation S43 disappears to give:

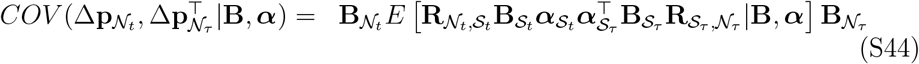

As in our inference section, the vector of allele frequency changes could be transformed using matrix 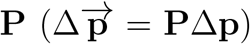 and B&C use the projection 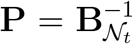 (they actually multiply this by 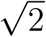 - see Equation S54) which results in

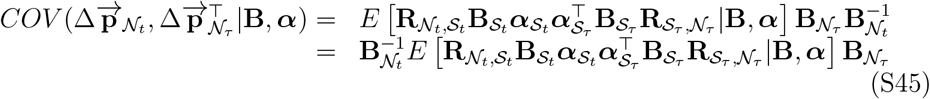

Further derivation in B&C considers the expected value of a diagonal element of this matrix: the average (over neutral loci) covariance in projected allele frequency change. However, it is perhaps easier to note that the trace of this matrix is equal to this average multiplied by the number of neutral loci, 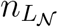 (see below). Since the trace of an outer product is equal to the inner product we get:

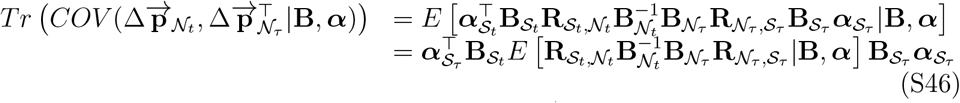

The diagonal element *j* of 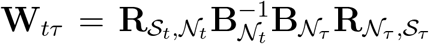 is equal to the sum of selected locus ‘s *R*_*t,ji*_*R*_*τ,ji*_(*b*_*τ,i*_*/b*_*t,i*_) across all neutral loci *i* . The *jk* ^*th*^ off-diagonal element is the sum of *R*_*t,ji*_*R*_*τ,ki*_(*b*_*τ,i*_*/b*_*t,i*_) for selected loci *j* and *k* across all neutral loci *i*. If we write **W**_*tτ*_ = **H**_*tτ*_ + (**W**_*tτ*_ − **H**_*tτ*_) where **H**_*tτ*_ and **W**_*tτ*_ − **H**_*tτ*_ are zero but for the diagonal and off-diagonal elements respectively, then Equation S46 becomes

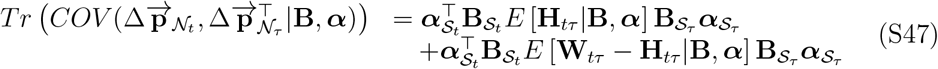

If we focus on a system with two selected loci, *j* and *k*, then the final term in Equation S47 is equal to:

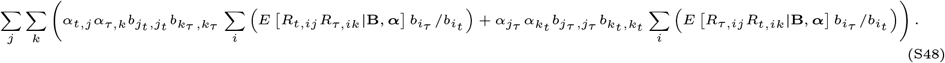

Under Hill-Robertson interference, if *α*_*j*_ and *α*_*k*_ have the same sign we expect them to be in negative LD with each other, and as a consequence have opposing patterns of LD with the neutral loci (i.e. if *α*_*j*_*α*_*k*_ *>* 0 then we expect *E*[*R*_*i,j*_*R*_*i,k*_] *<* 0 and vice versa). Although the terms of these products are evaluated at different generations in Equation S48 (*t* and *τ*) we expect the terms to share sign in the same way, generating a negative expectation for the second term in Equation S47. However, assuming an absence of Hill-Robertson interference, or signed linkage-disequilibrium more generally (Assumption D), then the final term in Equation S47 can be dropped:

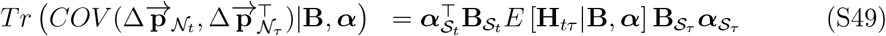

Based on a deterministic model for changes in linkage-disequilibrium (Assumption E) and assuming nongametic-phase linkage-disequilibrium is absent (Assumption F), Equations 40-44 in B&C derive an expression from which *E* [**H**_*tτ*_ | **B, *α***] can be computed. Under the assumption that drift (or selection) does not alter the dynamics of **R** then (Equation 42 in B&C):

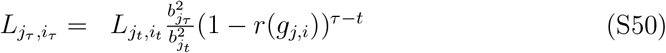

which implies

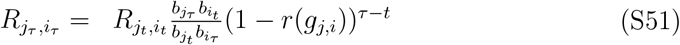

and so

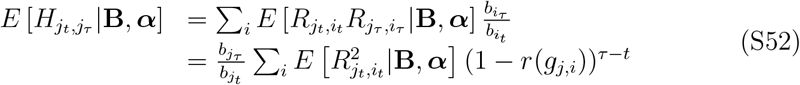

where *r*(*g*_*j,i*_) is the recombination rate as a function of the distance *g*_*i,j*_ between the two loci. Writing **F**_*tτ*_ a diagonal matrix with the *j*^*th*^ element equal to 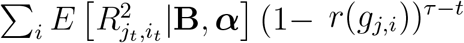, then 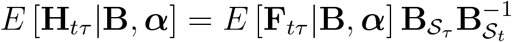 and so

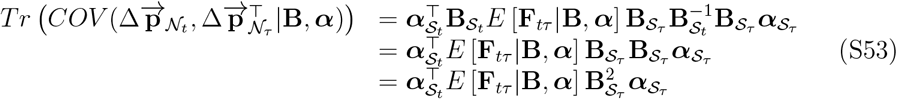

where (under random mating) 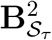 is a diagonal matrix with elements proportional to the genetic diversities in generation *τ* . However, this is still hard to evaluate since *E* [**F**_*tτ*_ | **B, *α***] will vary over loci in a way that may depend on **B** and hence 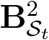. The method of B&C assumes that the sample correlation between the diagonal elements of *E* [**F**_*tτ*_ | **B, *α***] and the elements 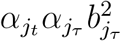 is zero (Assumption G). Under this assumption:

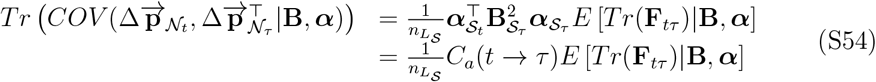

where 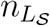 is the number of selected loci (note that in B&C the leading term is 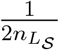 not 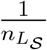 and this is because our projection matrices are only proportional by a factor 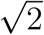 - see Equation S45). The method of B&C further assumes (Assumption H) that the average effects are constant in time such that ***α***_*t*_ = ***α***_*τ*_ then this reduces to

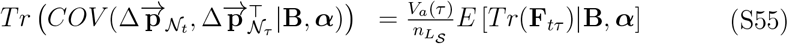

Under Assumption H, Assumption G implies that the additive genic variance contributed by a selected locus is independent of its associations with neutral loci as measured by 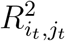. However, allele frequencies at selected loci enter the expression for the additive genic variance, and they also dictate the range of 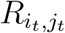 and there-fore the magnitude of 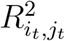: when selected allele frequencies are small compared to frequencies at neutral alleles, 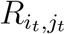 cannot cover the full range of -1 to 1 (Sved and Hill, 2018).

B&C approximate **R**_*t*_, and therefore **F**_*tτ*_ under the assumption of mutation-driftrecombination equilibrium (Assumption I). Ohta and Kimura (1971) derived the expectation of 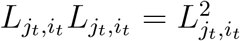 under this assumption, although the expectation of 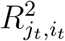 can only be approximated as the expectation of 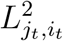 divided by the expectation

of the genetic diversities at the two loci, and is only accurate when the minor allele frequencies are greater than 10% (McVean, 2002) – and can be out by orders of magnitude when allele frequencies are extreme (Song and Song, 2007), as can be expected at loci under selection. Moreover, Ohta and Kimura (1971) derived the expectation 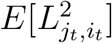 yet the B&C approach actually requires 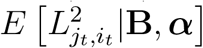 which is considerably more challenging to compute (Good, 2022). In the Appendix B&C also relax, to some extent, Assumption I, where *Tr*(**F**_*tτ*_) is calculated empirically using all loci, neutral and selected (Equation 55). In both cases - using the neutral expectation (Assumption I) or empirical LD (Assumption I-b) - the expected LD between selected and neutral loci will be overestimated since in reality selected alleles will be rarer than neutral alleles and so their LD, even measured as a correlation, will be reduced compared to that between neutral loci (Sved and Hill, 2018).

Equation S55 allows *V*_*a*_(*τ*) to be estimated, but B&C aim to estimate *V*_*a*_(*t*). Un-der Assumption H, and assuming that the proportional change in genetic diversity at selected loci between generation *t* and *τ* is constant across loci (Assumption J: 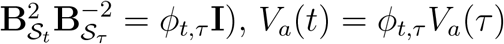.

The above derivation assumes that the sites can be partitioned into selected and neutral sites and that the map positions of all sites are known. In practice, this will often be infeasible and so a number of additional assumptions are required. In the absence of map positions, Haldane’s (1919) mapping function is assumed for *r*(*g*) (Assumption K) but, since the selected loci and their physical position, *g*, are assumed unknown, a model is also required for *g*. B&C assume that selected and neutral loci are distributed uniformly and independently such that *g*_*j,i*_ has a triangular distribution (Assumption L). Also, since the selected sites are unobserved, *ϕ*_*t,τ*_ cannot be computed and so it is assumed that that *ϕ*_*t,τ*_ is equal to the ratio of genetic diversity at generation *τ* to genetic diversity at generation *t* across all neutral loci (Assumption M).

In addition to the assumptions/approximations made when developing the theory, additional assumptions/approximations are made when making inferences from data. Rather than taking the average (over loci) covariance (over evolutionary replicates) the covariance over loci is taken. However, from the law of total covariance this is expected to yield the correct result under Assumptions B and C. In addition, rather than calculating the (co)variances in projected allele frequency change, it is assumed (Assumption N) that the (co)variances in actual allele frequency divided through by the average projection is a good approximation: 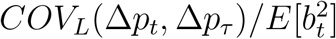 is a good approximation of *COV*_*L*_(Δ*p*_*t*_*/b*_*t*_, Δ*p*_*τ*_ */b*_*t*_) where *COV*_*L*_ designates a covari-ance over loci (Equation 16 B&C). The two quantities are only expected to agree under restrictive conditions (Bohrnstedt and Goldberger, 1969). However, it is not necessary to make Assumption N and relaxing it can reduce the bias in the estimator considerably. Finally, if there is measurement error in allele frequencies and the same allele frequency measurements are used to calculate change over adjacent intervals then this will generate downward bias in the estimated covariances. This is corrected for in B&C by assuming the sampling error in allele frequencies is binomial around the true value (Assumption O), although in practice non-binomial causes of overdispersion are likely.

To connect Equation S55 with the derivation in B&C more clearly, we can write

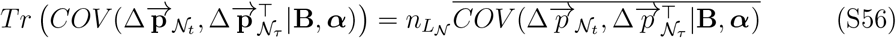

where the overline denotes the sample mean for the 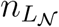 neutral loci. Similarly,

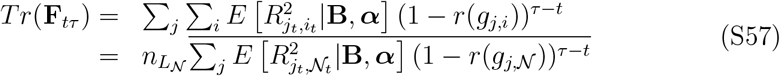

Equation S55 can then be written as

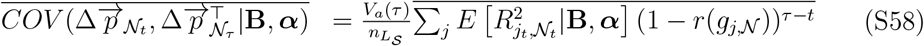

which is Equation 8 in B&C averaged over neutral loci and can be further simplified to

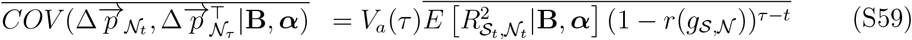

where the overline indicates the sample average of all the 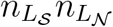 pairwise comparisons between selected loci and neutral loci.

Extending this theory to allele frequency change measured in independent replicates, rather than at different time-points in a single population is straightforward (Buffalo and Coop, 2020). If allele frequency change is measured in two populations, *m* and *n*, both of which have been derived independently from the base population *t* generation in the past, Equation S59 can be expressed as:

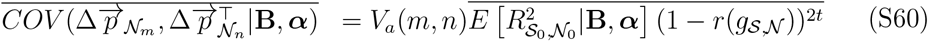

assuming the average effects have stayed constant.

## S8 *N*_*E*_ and average effects under a log-linear model

In our simulations, we model the logarithm of absolute fitness of an individual *k* using an additive model across loci:

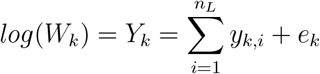

where *y*_*k,i*_ is the genotypic contribution of locus *i* to the log absolute fitness of individual *k*, and the environmental deviations, *e*, have zero mean and standard deviation *σ*_*e*_. We allow within-locus deviations from additivity using a fitness-scheme equivalent to the one employed by Lynch and Walsh (1998) (pp. 67) adapted for *proportions* of reference alleles as opposed to counts:

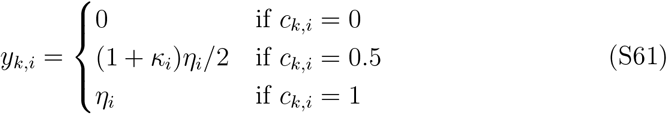

Note that in the parameterisation of Falconer and Mackay (1996), the genotypic value of the heterozygotes is (*η*_*i*_ + *d*_*i*_)*/*2 such that *d*_*i*_ = *κ*_*i*_*η*_*i*_.

We then draw 2*N*_*t*+1_ parents with replacement from a multinomial with the probability of *k* being a parent of an offspring proportional to its absolute fitness *W*_*k*_ = *exp*(*Y*_*k*_). Following Kojima (1959) we can define Fisher’s average effect for relative fitness in terms of the partial derivative of mean fitness with respect to each *E*[*c*] = *p*_0_ and then rescale by mean fitness:

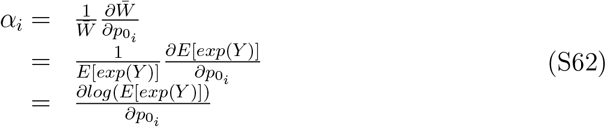

Assuming *Y* to be normally distributed with mean *µ*_*Y*_ and variance *V*_*Y*_ allows us to use the expression for the expectation of a log-normal distribution to write:

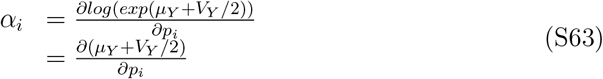

It is worth highlighting that, so far, we have made no assumptions about the distribution of the contributions (*y*_*i*_) made by individual loci to *Y* .

Next, we assume Hardy-Weinberg genotypic frequencies to write:

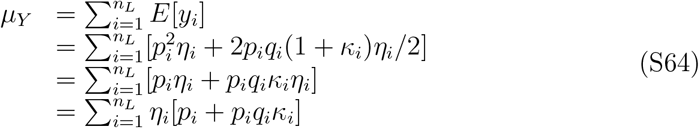

Differentiating with respect to *p*_*i*_ yields:

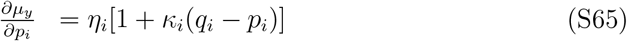

Assuming that all additive genetic variance is genic:

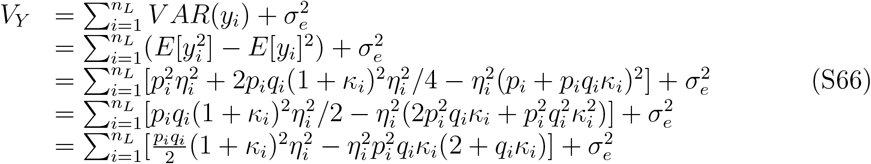

Differentiating with respect to *p*_*i*_:

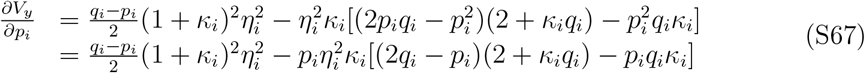

Substituting expressions for 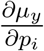 and 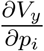 yields:

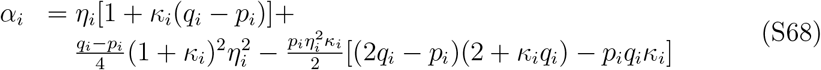

In the parameterisation of Falconer and Mackay (1996), this translates to:

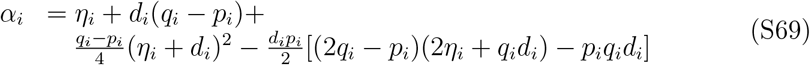

Note that as a first approximation Equations S69 and S68 reduce to the classical quantitative genetic expressions for the average effect of gene substitution (Equation 7.5 in Falconer and Mackay (1996) and Equation 4.10b in Lynch and Walsh (1998)).

In order to calculate *N*_*E*_ under this multinomial log-linear model we need to know the variance in offspring number *V*_*o*_ caused by any environmental or non-additive genetic variance. Then *N*_*E*_ = 4*N/*(2 + *V*_*o*_) where *N* is the census population size (Wright, 1938). The expected number of offspring for parent *k* is 2*N*_*t*+1_*p*_*k*_, as given above, and the variance in the number offspring is 2*N*_*t*+1_*p*_*k*_(1 − *p*_*k*_). From the the

law of total variance 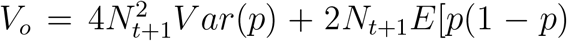. Since *E*[*p*(1 − *p*)] = *E*[*p*] *E*[*p*^2^] and *V ar*(*p*) = *E*[*p*^2^] − *E*[*p*]^2^ then *E*[*p*(1 − *p*)] = *E*[*p*] − *V ar*(*p*) + *E*[*p*]^2^. Since *E*[*p*] = 1*/N*_*t*_ by definition,

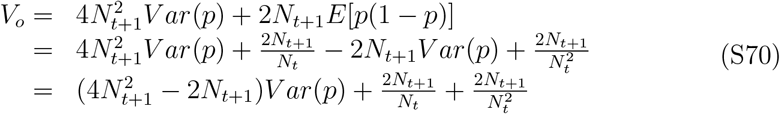

From the properties of the log-normal (with zero mean) we know 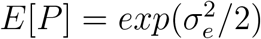 and 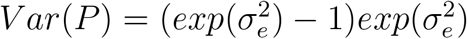. Since *p* = *P/*(*N*_*t*_*E*[*P*])

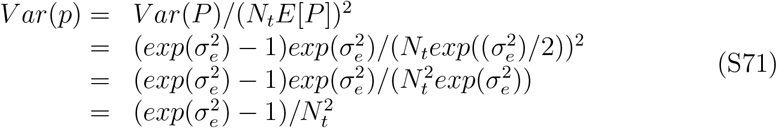

If *N*_*t*+1_ and *N*_*t*_ are large then Equation S70 simplifies to

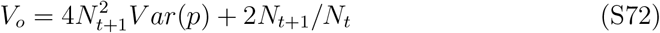

such that

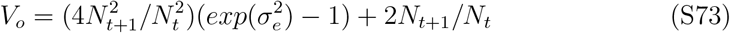

If the population size is constant then this simplifies to 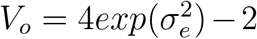 and the variance effective population size, 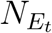, is 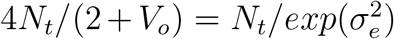. Note however, that in these expressions using 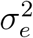 only assumes the non-additive genetic variance is zero. When non-additive genetic variance is present (which there will be to a small degree even with additivity on the log-scale) *V*_*o*_ will be greater than predicted here and *N*_*E*_ will be consequently smaller.

## S9 Workflow for implementing the method

## Box 1. General workflow of the method

We use genome-wide allele frequency change data (Δ**p**) from independent evolutionary replicates derived from the same ancestral base population with a known linkage structure (**L**_0_) to infer the mean 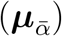 and the variance 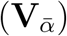 of the distribution of average effects for fitness. Our goal is to then estimate an expectation for the additive genetic variance for relative fitness by averaging over the distribution of average effects; i.e. 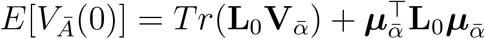. We use simple, yet biologically sensi-ble models, 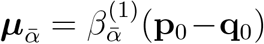 and 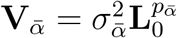, which essentially allows the average effect for fitness at a site to depend on allele frequency and genetic diversity. This model can be readily implemented as a linear mixed model for suitably projected allele frequency change treating locus effects as random (see below). This, effectively, allows us to partition the total allele frequency change into a component caused by predictable responses to direct and indirect selection (modelled using the random effects of locus) and a component caused by unpredictable responses to selection as well as genetic drift (absorbed by the model residuals).

### Experimental design and data

A typical experimental design permitting the implementation of our method would involve a base populations of *N*_0_ individuals which are sequenced individually. A number of independent evolutionary replicate populations derived from the base populations are then allowed to evolve between generations *t* and *τ* after the base population. If the replicate populations are derived from the base population after just a single round of reproduction *t* would be 1.

The following data are compiled from the experiment:

**C**_**g**,**0**_: A matrix with 2*N*_0_ rows and *n*_*L*_ columns indicating the presence or absence (1 or 0) at each of the *n*_*L*_ segregating sites in each of the 2*N*_0_ gametic contributions in the base population. If phased genomes are unavailable, the *N*_0_ *× n*_*L*_ matrix of the proportions of the copies of the (arbitrarily chosen) reference allele at each locus in each individual (**C**_**0**_) may be used.

Δ**p**_*m*_: For each replicate *m*, a vector representing allele frequency change at each segregating locus between generations *t* and *τ* after the base populations. *t* and *τ* must be known.

**R**_−_: An *n*_*L*_ *× n*_*L*_ matrix with 1’s on the diagonal and the probabilities of non-recombination between pairs of sites as off-diagonal elements. A recombination map if available, may be used to construct this matrix.

#### Step 1: Compute 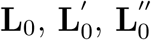, and 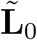

First, we compute the covariance of **C**_**0**_ over individuals to obtain **L**_0_, and the covariance of **C**_**g**,**0**_ over gametic contributions to obtain the matrix of gametic phase disequilibria 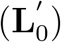. The matrix of the non-gametic phase disequilibria 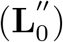 is then given by 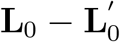 allowing us to compute 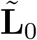 as a weighted sum of 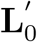 and 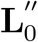, with the the *ij*^*th*^ element of the latter being weighted by *r*_*ij*_*/*(1 − *r*_*ij*_), where *r*_*ij*_ is the probability of recombination between the two loci. If phased genomes are unavailable, 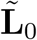 may be set to **L**_0_ which assumes that the non-gametic phase disequilibria are negligible.

#### Step 2: Compute the predicted L in generation *t* in replicate *m* (**ℒ**_*t,m*_)

Given 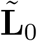 in the base population, the expected **L** in generation *t* in replicate *m* under the action of drift and recombination is given by 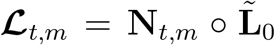, where the *ij*^*th*^ element of **N**_*t,m*_ is 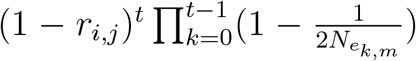, and *∘* indicates Hadamard (i.e. element-wise) product. We then sum the **ℒ**_*t,m*_’s for all the generations over which Δ**p**_*m*_ is measured to obtain 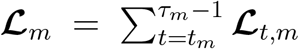. Note that the expected predicable change due to selection between generations *t* and *τ* is 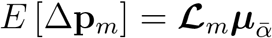 and the among-replicate covariance in allele frequency change is 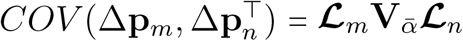.

#### Step 3: Compute drift covariance and the projection matrix

Instead of working with raw allele frequency changes, we work with *projected* allele frequency changes. The projection matrix **P** is chosen to be such that (1) the number of dimensions of the dataset are reduced to the minimum of *N*_0_ and *n*_*L*_ (typically *N*_0_ *<< n*_*L*_), and (2) the allele frequency changes due to drift are independent and identically distributed. First, we perform eigen-decomposition of **L**_0_ (or, equivalently, singular-value decomposition of **C**_**0**_) and store the eigenvectors corresponding to non-zero eigenvalues (**U**_**L**_). The (co)variance in allele frequency change due to drift in replicate *m* is 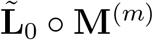 where 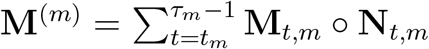 and the *ij*^*th*^ element of **M**_*t,m*_ is 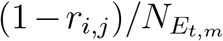 . We then perform eigen-decomposition of the drift covariance expressed using the eigen-vectors of **L**_0_, i.e. 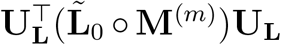. Let **U**_2_ be a matrix having the eigenvectors of this 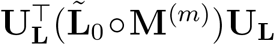 as columns, and let **D**_2_ be a diagonal matrix of square-rooted eigenvalues of this 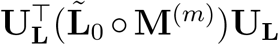. We can then calculate the projection matrix 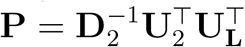, and obtain the projected allele frequency change vector for replicate *m* as 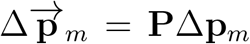. Note that the mean and covariance of the projected allele frequency changes due to predictable selection are 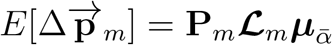 and 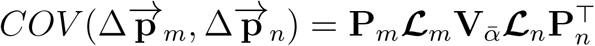.

#### Step 4: Preparing data for model

Next, using our models 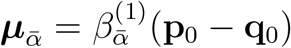 and 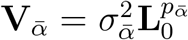 we calculate the fixed effect covariate P *ℒ* _*m*_(p_0_ − q_0_), and the covariance structure of the locus effects as 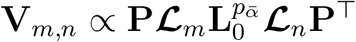.

#### Step 5: Fit linear mixed models

For a given 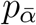, our goals is to fit a linear mixed model with the projected allele frequency change in all replicates 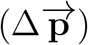 as the response variable and a fixed effect predictor given by **P *ℒ*** _*m*_(**p**_0_ − **q**_0_) with a coefficient *β*^(1)^ to be estimated. We treat locus effects to be random with a covariance structure assumed proportional to 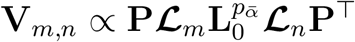, with a proportionality constant 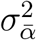 to be estimated. We use the R function *‘optim()’* to find 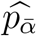 that maximises the conditional likelihood of the above model. We then use 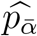, fit the above model, and obtain 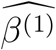 and 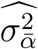.

#### Step 6: Estimate *E*[*V*_*Ā*_(0)]

The final step is to use the estimates obtained in the previous step (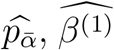, and 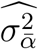) to obtain 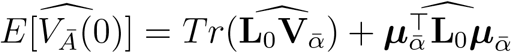. We obtain 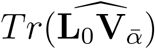 as 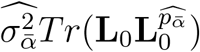, and bias corrected 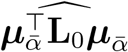 as 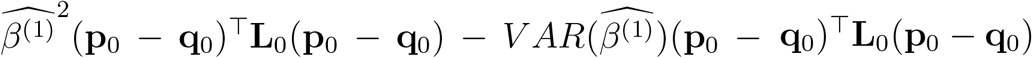.

## S10 Supplementary figures

**Supplementary Figure 1:**
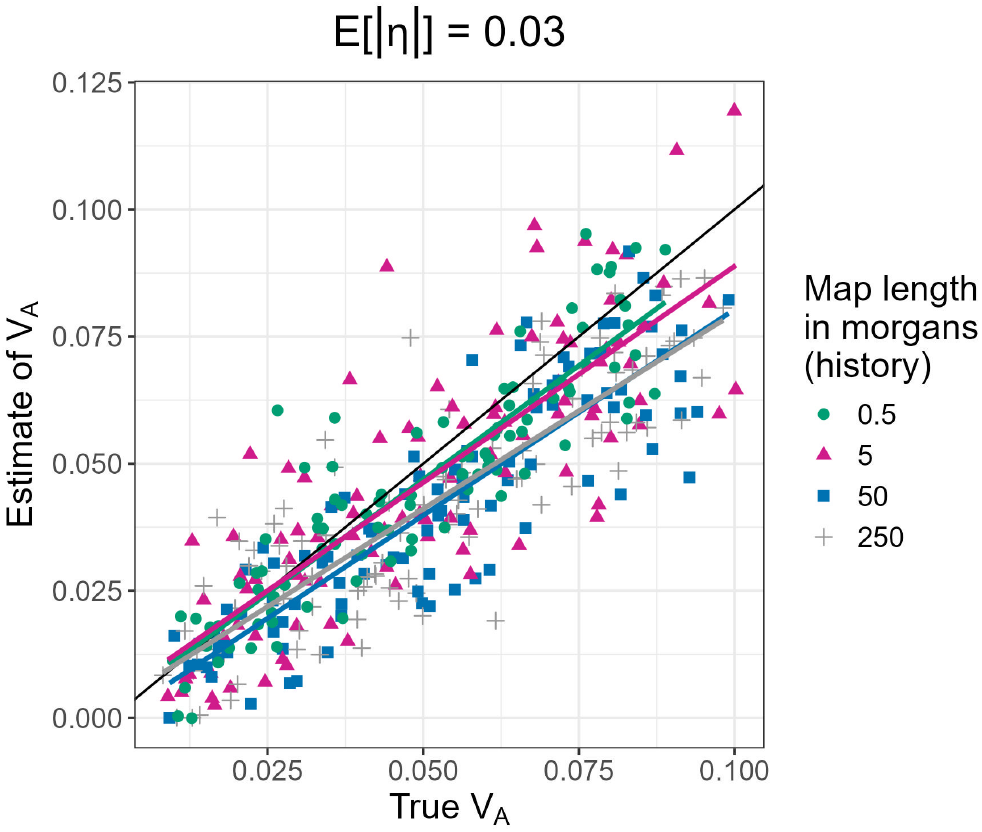
Results of full simulations with a burn-in phase of 25,000 generations (map length in the history phase = 2 morgans, number of replicate populations = 10, population size = 1,000, number of generations = 3, the mean of the gamma distribution from which effect sizes for log absolute fitness are sampled for non-neutral mutations (*E*[|*η*|]) = 0.03, at different levels of the map length in the history phase (0.5 morgan: green circles, 5 morgan: magenta triangles, 50 morgans: blue squares, and 250 morgans: grey plus symbols). The coloured solid lines represent regression lines for estimates of *V*_*A*_ vs true values of *V*_*A*_.

**Supplementary Figure 2:**
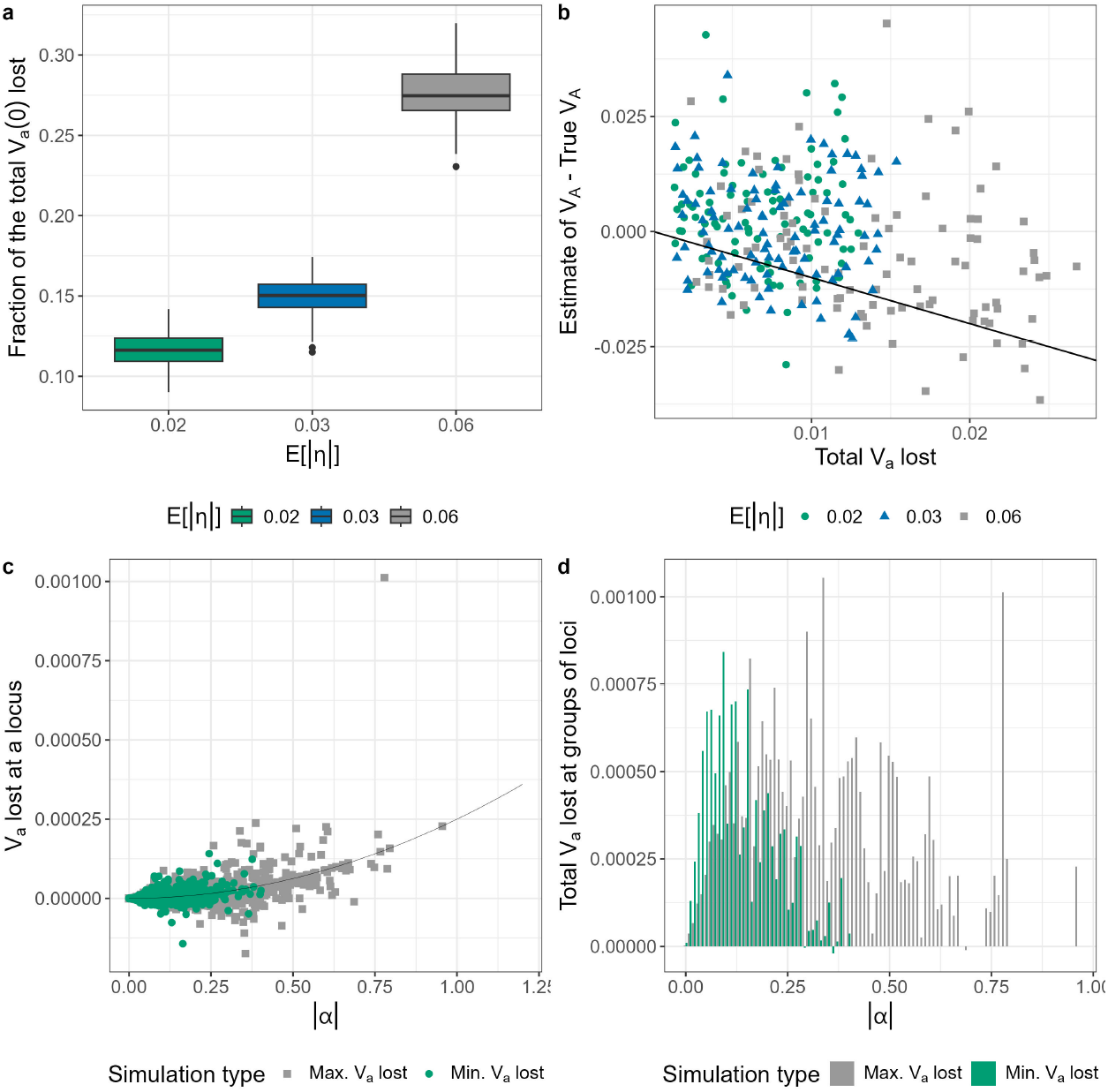
(a) Fraction of the total additive genic variance (*V*_*ā*_(0)) lost during the experiment (averaged over all the replicate populations) plotted as a function of the mean of the gamma distribution from which effect sizes for log absolute fitness are sampled for non-neutral mutations (*E*[| *η*|]). Each boxplot represents 100 independent simulations (for the corresponding level of *E*[|*η*|]) shown in <u>Figure 5</u>D. (b) The error in our estimates of *V*_*A*_ plotted versus the total additive genic vari- ance (*V*_*a*_) lost during the experiment (averaged over all the replicate populations) for simulations where the *E*[|*η*|] was either 0.02 (green circles), 0.03 (blue triangles), or 0.06 (grey squares). The solid line has an intercept of 0 and a slope of -1. (c) The contribution of each non-neutral locus (averaged over all the replicate populations) to *V*_*a*_ lost during the experiment plotted versus the absolute average effect for relative fitness (|*α*|) for that locus for a simulation with *E*[| *η*|] = 0.06 that had the maximum *V*_*a*_ loss (grey squares) and a simulation with *E*[| *η*|] = 0.02 that had a comparable true initial *V*_*A*_ but the minimum *V*_*a*_ loss (green circles) in the experiment. The solid curve represents the additive genic variance lost when a singleton goes extinct. (d) The total *V*_*a*_ lost at groups of loci classified by dividing |*α*| into intervals of 0.01 for the simulation with the maximum *V*_*a*_ loss (grey) and the simulation with the minimum *V*_*a*_ loss (green) in the experiment. In the simulation with the maximum *V*_*A*_ loss – in which the mean |*α*| for non-neutral segregating loci was 0.0410 – more than 50% of the total loss in *V*_*a*_ was driven by just 0.96% loci whose |*α*| was greater than 0.3.

**Supplementary Figure 3:**
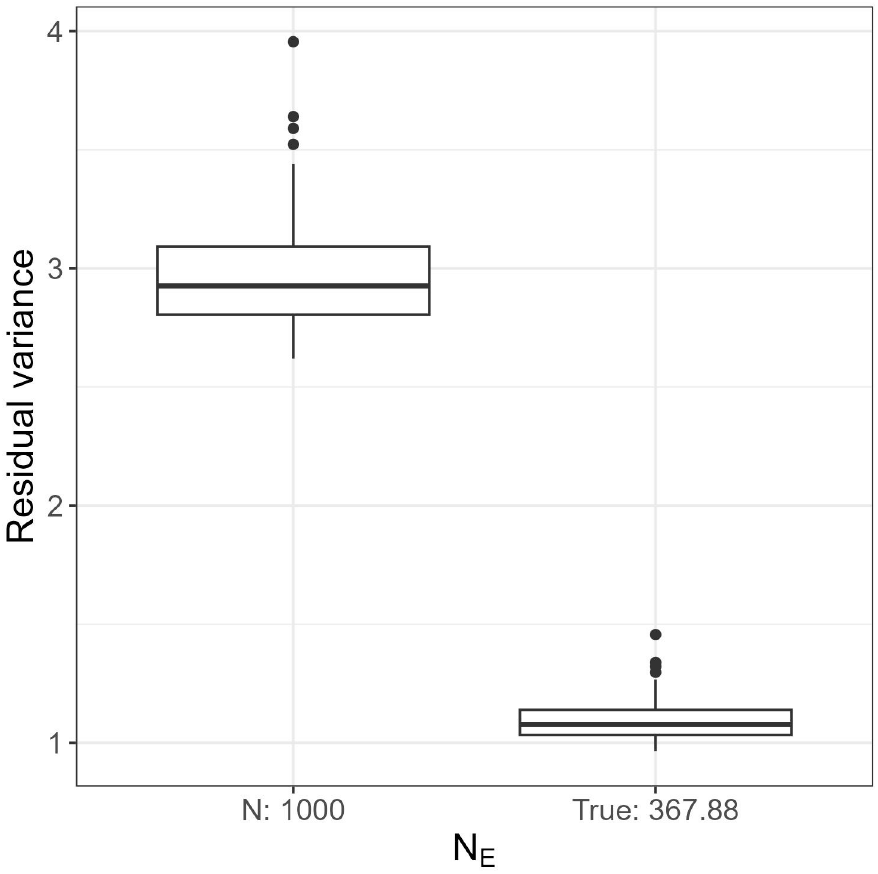
Residual variance in the models analysing the results of full simulations shown in <u>Figure 4</u> using either the number of individuals (1,000) instead of the *N*_*E*_ or the true *N*_*E*_.

**Supplementary Figure 4:**
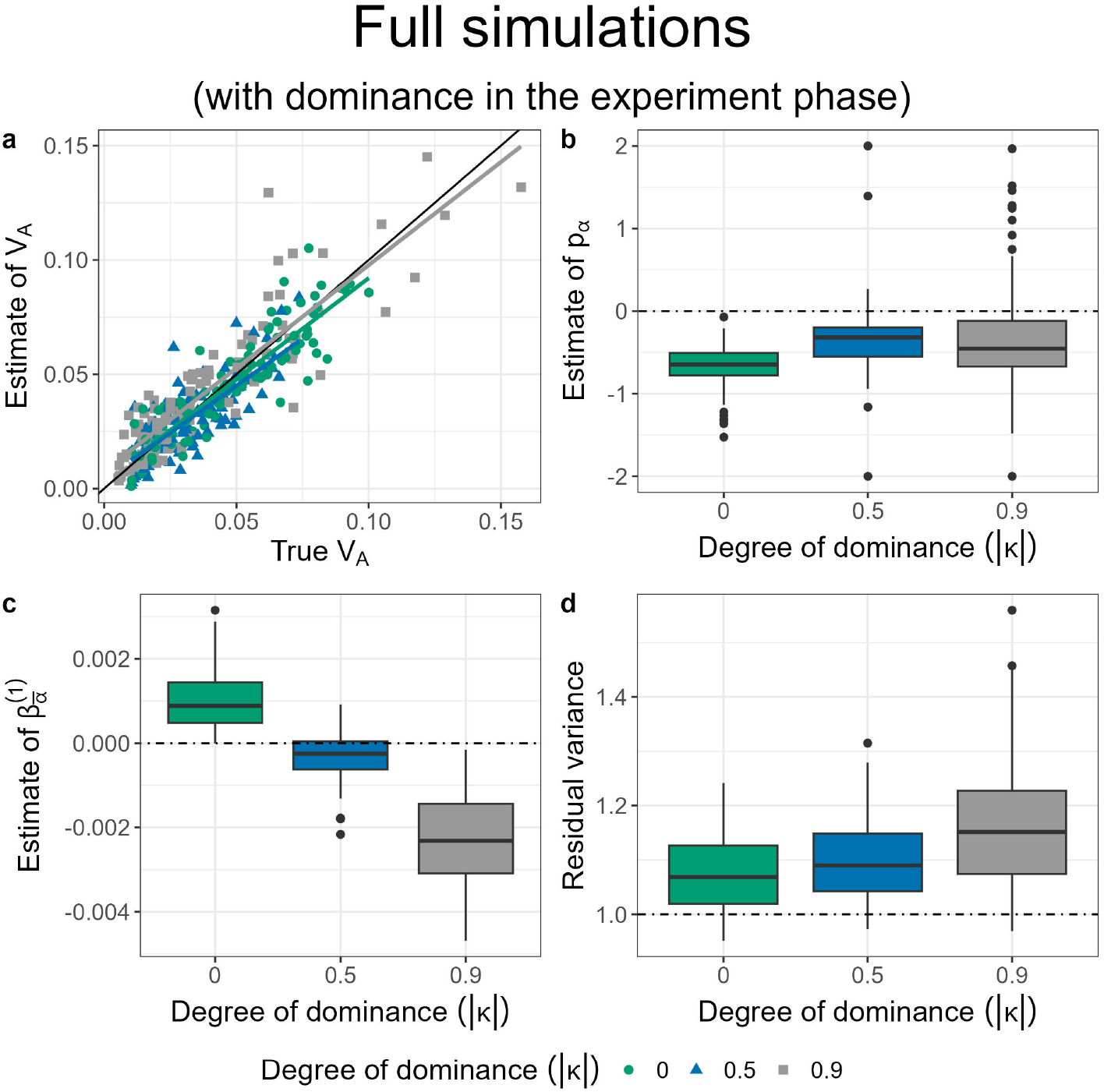
Results of full simulations (map lengths in the history phase = 0.5 morgans and experiment phase = 2 morgans, number of replicate populations = 10, population size = 1,000, number of generations = 3, the mean of the gamma distribution from which effect sizes for log absolute fitness are sampled for non-neutral mutations (*E*[| *η*|]) = 0.03), in which dominance effects were switched on only in the experiment phase using different degrees of dominance: |*κ*| = 0 (green circles), |*κ* | = 0.5 (blue triangles), and |*κ*| = 0.9 (grey squares). (a) A scatter plot of estimates of *V*_*A*_ vs true values of *V*_*A*_. The solid black line indicates the 1:1 line. The coloured lines represent regression lines for estimates of *V*_*A*_ vs true values of *V*_*A*_. The inference of *V*_*A*_ was obtained by modelling the mean and the (co)variance of the average effects for relative fitness as 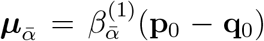 and 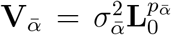, respectively. (b)-(d) Histograms of the estimates of 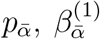, and the residual variance, respectively. The horizontal dashed lines indicate null expectations (0 for 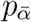 and 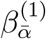, and 1 for the residual variance).

**Supplementary Figure 5:**
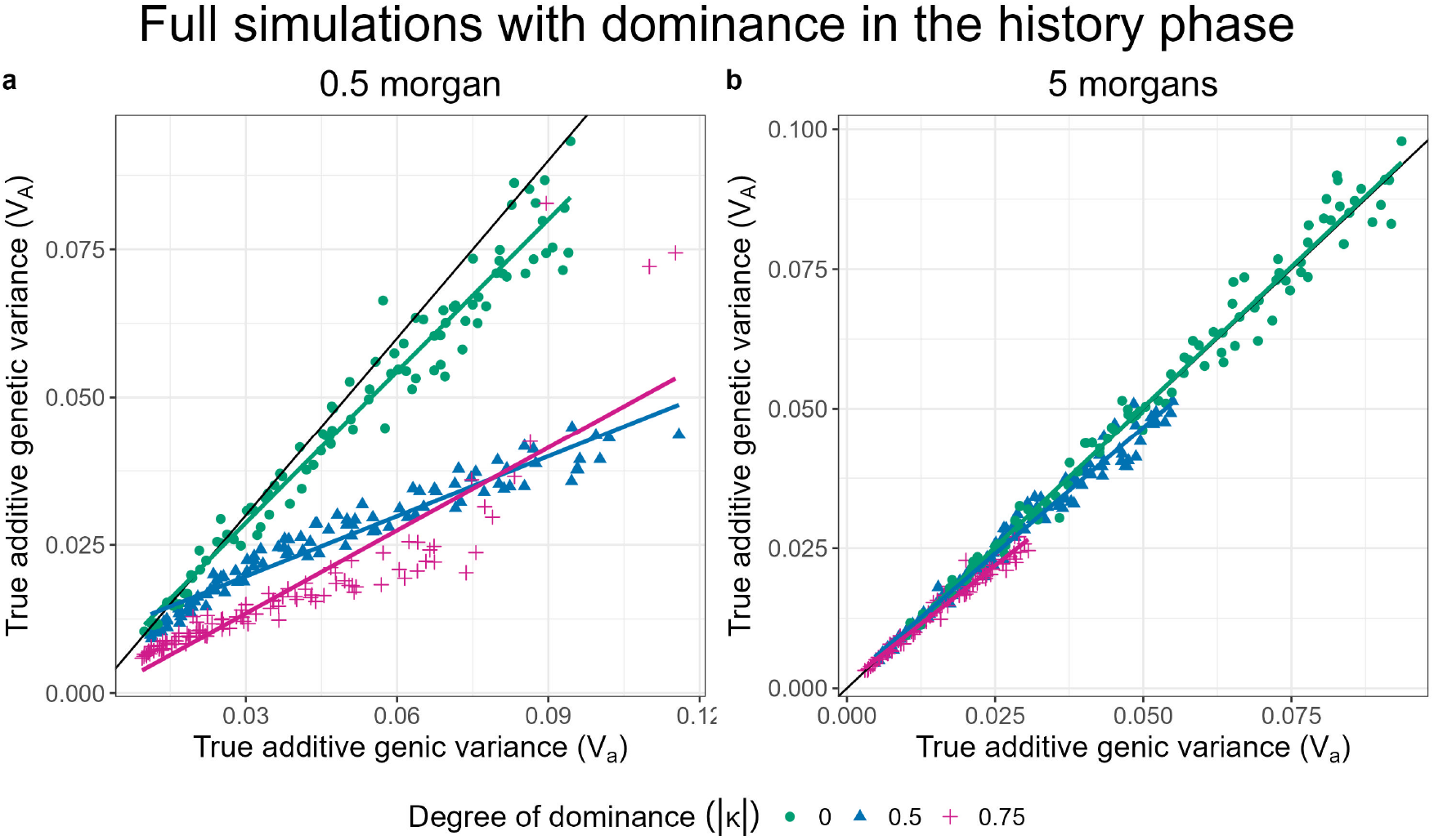
The true additive genetic variance for relative fitness (*V*_*A*_) plotted versus the true additive genic variance for relative fitness (*V*_*a*_) in full simulations with a burn-in phase of 25,000 generations performed using a map length in the history phase equal to either 0.5 morgan (a) or 5 morgans (b) with dominance effects switched on in the last 5,000 generations of the history phase. Three different degrees of dominance were used: | *κ*| = 0 (green circles), *κ* = 0.5 (blue triangles), and *κ* = 0.75 (magenta plus symbols). The mean of the gamma distribution from which effect sizes for log absolute fitness are sampled for non-neutral mutations (*E*[|*η*|]) was 0.03. The solid black lines indicates the 1:1 line. The coloured lines represent regression lines for true *V*_*A*_ vs true true *V*_*a*_.

**Supplementary Figure 6:**
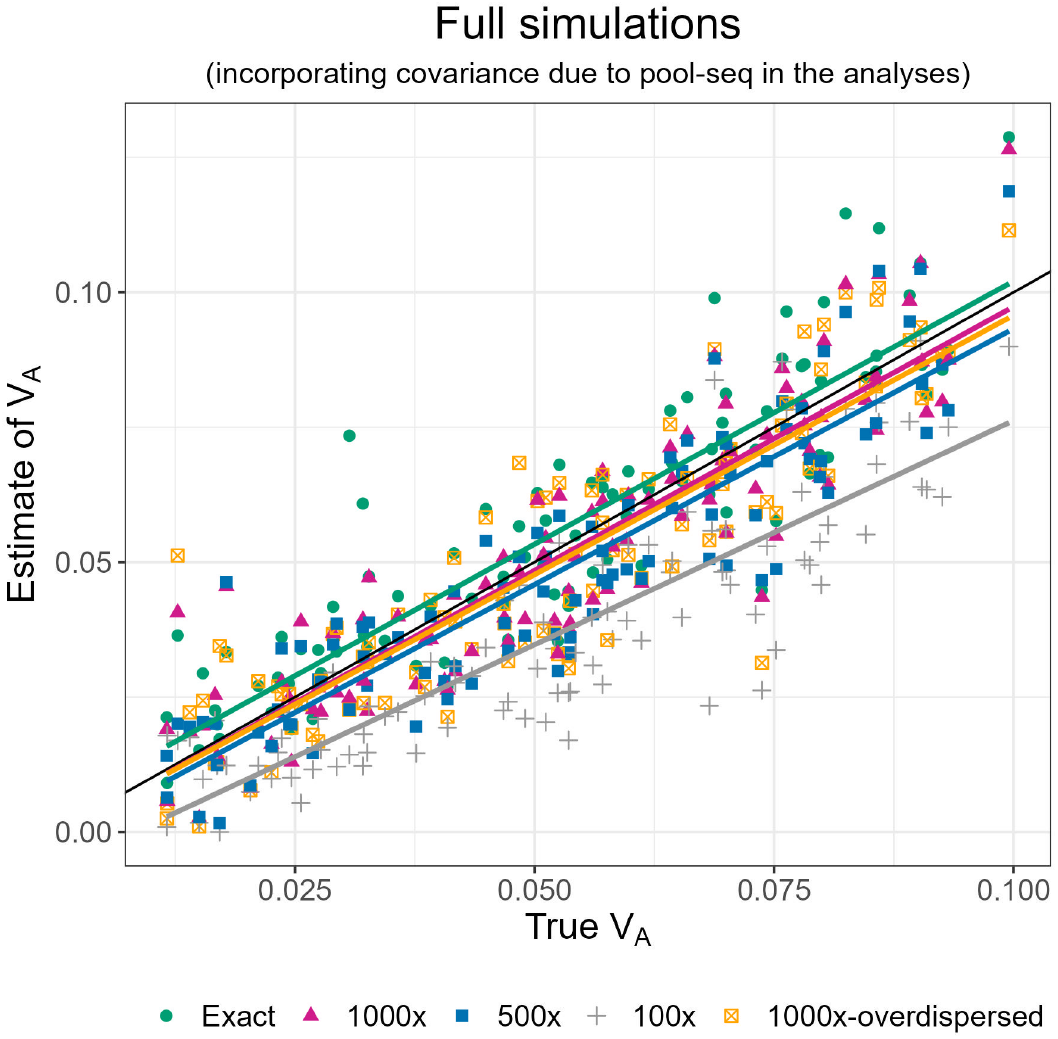
Scatter plots of estimates of *V*_*A*_ vs true values of *V*_*A*_ for full simulations with a burn-in phase of 25,000 generations (map length in the history phase = 0.5 morgan, map length in the experiment phase = 2 morgans, number of replicate populations = 10, population size = 1,000, number of generations = 3, the mean of the gamma distribution from which effect sizes for log absolute fitness were sampled for non-neutral mutations (*E*[|*η*|]) = 0.02, and no dominance (i.e. *κ* = 0)) using either exact allele frequencies in the experiment phase (green circles), or allele frequencies in the experiment phase obtained via simulated pool-seq implemented without any overdispersion in the number of reads mapping to an individual, at three different levels of coverage (expected number of reads mapping a segregating site: 1000x (magenta triangles), 500x (blue squares), and 100x (grey plus symbols)), as well as estimates obtained from simulated pool-seq implemented with overdispersion (*V*_*x*_ = *log*(2)) in the number of reads mapping to an individual, at 1000x coverage (orange empty boxes with crosses). Reads were modelled to be 37 base-pairs long. The solid black line indicates the 1:1 line. The coloured lines represent regression lines for estimates of *V*_*A*_ vs true values of *V*_*A*_. Estimates of *V*_*A*_ were obtained by incorporating the expected covariance structure due to pool-seq sampling in the models.

